# Extended nuclear glycosylation regulates RNA processing

**DOI:** 10.64898/2025.12.02.691741

**Authors:** Jon Lundstrøm, Michelle Fong, Annika Thorsell, Ekaterina Mirgorodskaya, Johannes Fuchs, Ujala Bashir, Jordi C.J. Hintzen, Chunsheng Jin, F. Ifthiha Mohideen, Vivian Lobo, Evgeniia Shcherbinina, Alesia A. Tietze, Lara K. Mahal, Aishe A. Sarshad, Daniel Bojar

## Abstract

In eukaryotes, glycans modify proteins in the secretory pathway and the extracellular space. Aside from nucleocytoplasmic *O*-GlcNAc, glycosylation is not considered a relevant post-translational modification in other cellular compartments. Here, we challenge this long-standing paradigm by showing that extended *O*-glycans are commonly found on intranuclear proteins. Through comprehensive genetic and biochemical analyses, we conclusively demonstrate that these *O*-glycans stem from the secretory pathway, yet are found on nuclear proteins across mammalian cell lines and primary cells. Using knock-out cell lines, we show mechanistically that nuclear glycans are shuttled to the nucleus via active vesicular transport. We identify several of these intranuclear glycoproteins as RNA-binding proteins, including KHSRP/FUBP2, RBM12, and RPP30. Lastly, we show that site-specific glycosylation of RPP30 is crucial for effective tRNA processing. Overall, our findings suggest a much broader role for glycosylation in regulating cellular functions and open up investigation into the role of glycans in more biological processes.

## Introduction

Glycosylation, which describes the addition of oligosaccharide chains to proteins and other biomolecules, is one of the most common classes of post-translational modification (PTM). It impacts the stability and function of proteins in the secretory pathway and beyond^1^. Glycosylation is dynamically regulated, structurally complex, and frequently dysregulated in disease^2^. Beyond the single example of *O*-GlcNAc, a strict monosaccharide-only PTM common in nucleocytoplasmic compartments^3^, glycosylation is thought to be irrelevant outside the secretory pathway and cell surface.

For decades, individual reports challenging the dogma of glycosylation as a secretory pathway phenomenon have accumulated, to the extent that this is even discussed in the *de facto* textbook of glycobiology^4^. The existence of glycan-binding proteins, such as galectins^5^, as well as glycan-degrading enzymes, such as sialidases^6^, in the nucleus and cytosol have long given rise to speculation. The biosynthesis of all activated monosaccharide precursors occurs in the cytosol and nucleus (e.g., CMP-sialic acid in nucleoli)^7^. The Human Protein Atlas localizes some glycan-binding proteins (e.g., Siglec-15) to the nucleus/nucleoli and nuclear proteins such as BIG1 (ADP-ribosylation factor guanine nucleotide-exchange factor 1) or DPY30 (Dpy-30 histone methyltransferase complex regulatory subunit) to the Golgi apparatus (“The multilocalizing proteome”)^8^. Photoproximity labeling has shown nuclear and cytosolic proteins close to sialylated proteins^9^, while Golgi sugar nucleotide transporters directly interact with nuclear proteins^10^. As described below, a cornucopia of glycoproteomics experiments have inadvertently reported soluble cytosolic and nuclear glycoproteins. Yet the prevailing view remains that extended glycosylation does not exist beyond the secretory pathway. Key arguments against existing data in conflict with the paradigm^4^ are contamination and misinterpretation of ER remnants that are contiguous with the nuclear envelope.

Numerous reports, using multiple methods and coming from different groups, have shown the existence of extended *O*-GalNAc type glycans in nuclei and other cellular compartments. The Irazoqui group has shown the presence of nuclear *O*-GalNAc type glycans, including the core 1 structure (Galβ1-3GalNAc), using various techniques including confocal microscopy, enzymatic assays, affinity chromatography, and mass spectrometry^11,12^. The Su group has used proximity labeling to identify nuclear *O*-GalNAc proteins^13^. *O*-GalNAc on S121 of p53 has been shown to affect protein stability, by the Zhang group^14^, supported by lectin blotting, glycosidase treatment, and click chemistry in GALE-knockout cells, to prevent confounding with *O*-GlcNAc. Other research documents the existence and biosynthesis of glycolipids in the nuclear envelope^6,15^. These (ignored) findings point to the existence of extended glycosylation in cellular compartments outside the secretory pathway, yet its existence, extent, and relevance remain contentious.

Independently, nuclear proteins without a signal peptide have been repeatedly localized in the trans-Golgi compartment, such as DPY30^16^ which is involved in histone methylation. These nuclear proteins exhibit interaction partners in both the nucleus and Golgi apparatus, suggesting an active exchange of material. Proteins involved in vesicular trafficking from the trans-Golgi, such as the guanine-nucleotide exchange protein ARFGEF1/BIG1, exhibit direct interactions with nuclear proteins and even contain a nuclear localization signal^17^. The localization of these shuttled proteins can be directly modulated by restricting Golgi-trafficking proteins such as BIG1 to the nucleus via an increase in cAMP levels^18^ or by a phospho-mimicking mutation^17^, while they can be restricted to the trans-Golgi by inhibiting microtubules that mediate vesicular transport^17^. These findings hint at a novel route of exchange between the trans-Golgi and the nucleus that could provide a mechanism for how nuclear proteins can be glycosylated.

The data we present here are explicitly designed to refute prior objections and incontrovertibly demonstrate the existence of extended nuclear *O*-glycans as well as establish a mechanistic model of the biosynthesis and trafficking of this new class of PTM. A rigorous purification procedure allows us to investigate pure mammalian nuclei from multiple cell lines and primary cells, without remnant ER. These nuclei exhibit clear lectin staining that can be removed by exoglycosidase treatment. Glycomics of these samples identifies extended *O*- but not *N*-glycans, further ruling out contamination. Following this up with glyco-enriched proteomics, we identify intra-nuclear proteins, enriched for RNA-binding proteins (RBPs), with extended *O*-glycans. CRISPR-Cas9 knockout (KO) screens and subsequent validation pinpoint the importance of both Golgi-localized nucleotide transporters and glycosyltransferases to nuclear glycosylation features, indicating a shared biosynthesis inside the secretory pathway. Disrupting vesicular transport via biochemical and genetic intervention abrogates nuclear glycosylation yet maintains cell surface glycosylation, highlighting a regulated mechanism for how glycoproteins reach the nucleus. Finally, we show that the glycosylation of RBPs such as the tRNA-processing RPP30 affect their function, indicating the role of extended nuclear glycosylation as a new layer of post-translational regulation and expanding the realm of glycobiology into a new subcellular compartment.

## Results

### Ultra-pure mammalian nuclei exhibit extended glycosylation

Two key requirements need to be met, in order to conclude that complex nuclear glycosylation is genuine, and overcome the paradigm of (exclusive) secretory pathway glycosylation (Fig. 1A): purity of nuclear preparation and presence of nuclear glycoproteins. To achieve the former and to exclude contamination by secretory pathway glycoproteins, we used and optimized a proven protocol of hypotonic lysis, differential centrifugation, and extensive washes^19–21^ to obtain pure nuclei of mammalian cells (Fig. 1B, Fig. S1, Fig. S2). Since the outer nuclear membrane is contiguous with the ER, we additionally optimized procedures^6^ to remove ER from our nuclei and demonstrated that our measured glycan signal did not stem from remnant ER (Fig. 1, Fig. S1, see Methods). Importantly, the resulting nuclei could no longer be stained in a Western blot for markers of the outer nuclear membrane (nesprin-4), ER (calnexin, calreticulin), or Golgi apparatus (GM130), while we still detected robust nuclear signal (Lamin A/C) and glycan signal (PNA; Fig. 1B). We thus conclude that our approach produces pure nuclei, without appreciable contamination from the secretory pathway.

**Figure 1.**
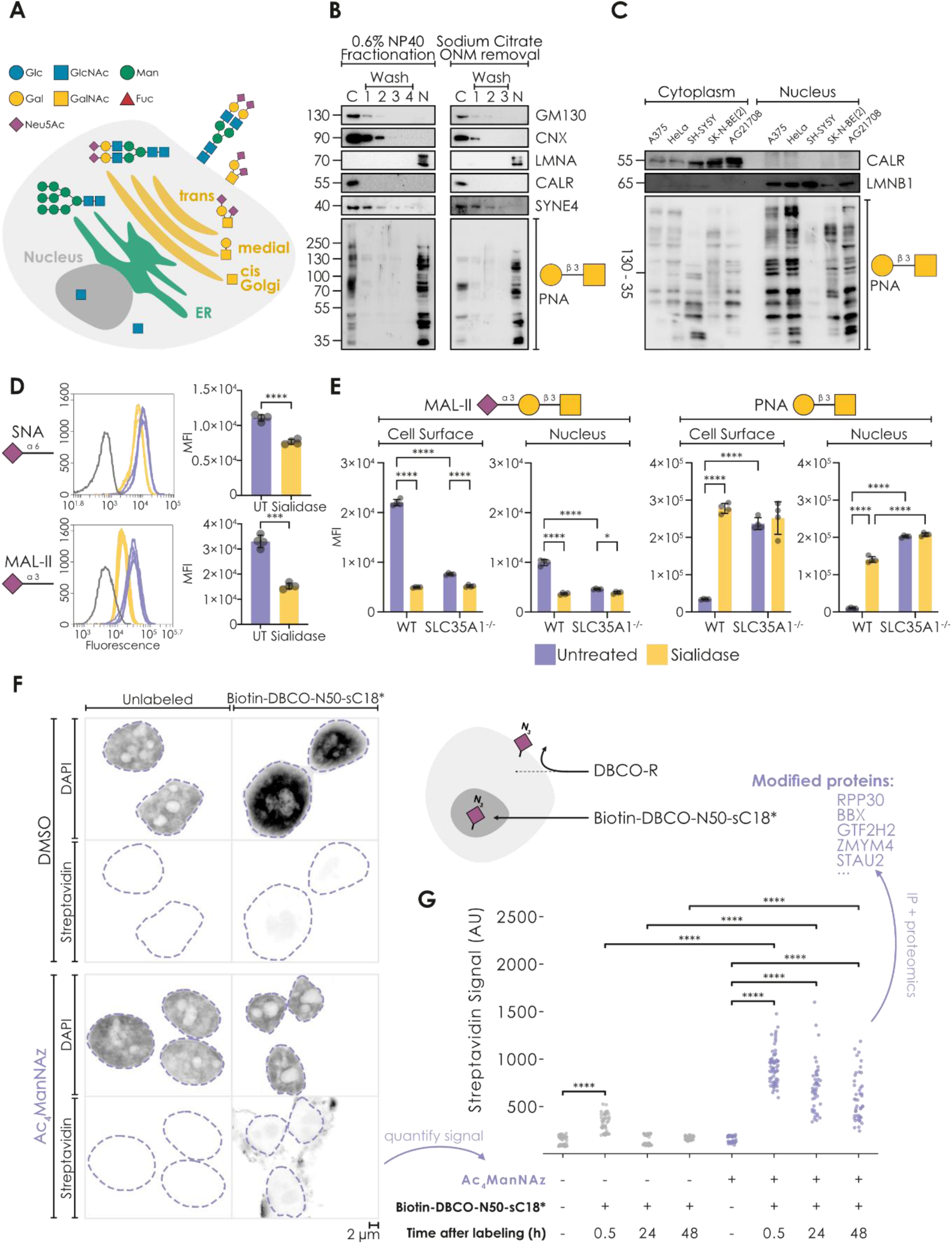
Extended glycans reside in mammalian nuclei. A) Schematic of conventional protein glycosylation and its subcellular distribution. B) Using a workflow involving either hypotonic lysis or ONM removal by sodium citrate, followed by extensive washing (1-4: subsequent washes), nuclei from A375 cells that lack detectable markers for Golgi apparatus (GM130), residual ER (Calnexin (CNX), Calreticulin (CALR)), and outer nuclear membrane (Nesprin-4 (SYNE4)) can be retrieved. C: cytoplasm, N: nucleus, ONM: outer nuclear membrane. Representative blots (n = 3). C) Extended nuclear glycosylation is a common feature of mammalian cell lines. Western blots of cytoplasm or nuclei from five human cell lines: A375 (melanoma), HeLa (cervical carcinoma), SH-SY5Y and SK-N-BE(2) (neuroblastoma), and AG21708 (primary dermal fibroblasts) stained with PNA or markers (ER: CALR; nucleus: Lamin B (LMNB1). Representative blots (n = 3). Subset of Fig. S4. D) Treating fixed and permeabilized A375 nuclei with sialidase abolishes the binding of sialic acid binding lectins MAL-II and SNA, as measured via flow cytometry (n = 4). Unstained controls are shown in grey in the histograms. UT: untreated. E) Knocking out sialic acid in the secretory pathway (via the CMP-sialic acid transporter SLC35A1) abrogates sialylation and exposes terminal galactosylation on nuclei and cell surface. Flow cytometry of HEK293T SLC35A1-/- or wildtype (WT) cells/nuclei stained with FITC-MAL-II (sialic acid) or PNA (T-antigen) (n = 4). Samples were treated with sialidase as a control (yellow bars). F) Confocal microscopy shows metabolic labelling of nuclear sialic acids. A375 cells were treated with 50 µM ManNAz for 72h to metabolically label sialic acids. Azides were detected 0.5 h after labeling with the Biotin-DBCO-N50-sC18* peptide via streptavidin staining. Controls with unlabeled cells and cells without peptide are shown. Scale bar: 2 µm. G) Time course of residence of nuclear-homing peptide in control / ManNAz-treated nuclei, measured via confocal microscopy (n = 44-67). Only the areas denoted by dashed lines in (F) have been used for quantification. Example nuclear proteins modified with the peptide are provided (full list in Table S2). All glycans in this work have been drawn with GlycoDraw22 and conform with the Symbol Nomenclature For Glycans (SNFG). Significant differences were established via a Welch’s t-test (two conditions) or a one-way ANOVA with Tukey’s multiple comparison test (>2 conditions). *p < 0.05, **p < 0.01, ***p < 0.001, ****p < 0.0001.

Next, we probed these purified nuclei for glycan signal with commonly used lectins^23^, which are specific for select glycan epitopes indicative of extended *O*-glycan structures. We report that, in A375 melanoma nuclei, we detect robust staining with lectins such as VVL (GalNAc), PNA (Galβ1-3GalNAc, T-antigen), SNA (Neu5Acα2-6), and MAL-II (Neu5Acα2-3Gal; Fig. S3). Since MAL-II is known to only bind its epitope in the context of *O*-GalNAc glycans^23^, we concluded that nuclear glycans contained extended sialylated *O*-glycans. Next, we wanted to ascertain whether this phenomenon could be observed in cell lines other than A375. We screened a panel of mammalian cell lines (A375: melanoma, HeLa: cervical carcinoma, SH-SY5Y and SK-N-BE(2): neuroblastoma, AG21708: primary dermal fibroblasts, HCT116: colon cancer, MCF7: breast cancer, A549: lung cancer, HEK293T: immortalized kidney embryonic cells, and E14: murine embryonic stem cells) using lectin blots (Fig. 1C, Fig. S4). Except for SH-SY5Y and MCF7 cells, the tested nuclei all produced robust, yet cell type-specific, staining behavior with lectins. We also note that the stained bands typically differed from those observed in the remaining lysate from the same cells.

To further confirm our findings, we next performed flow cytometry on extracted nuclei using lectins. This revealed clear nuclear signal, which was generally lower than the cell surface signal, in all tested cell lines (A375, HEK293T, THP-1, and non-dividing, PMA-differentiated THP-1 cells, Fig. 1D, Fig. S5). Importantly, since MAL-II and SNA both recognize sialylated epitopes on extended glycans, we treated extracted nuclei with sialidase, which removes sialic acid, as a control. In line with expectation, we observed a decrease in lectin binding, confirming that the signal was glycan-dependent (Fig. 1D, yellow bars). Coupled with the established purity of our nuclei, we thus were confident that this demonstrated the presence of extended glycan signal from ultra-pure nuclei.

To ensure that nuclear glycoconjugates were formed in living cells and not post-lysis, we next acquired a HEK293T*^SLC35A1-/-^* knockout (KO) cell line^24^. Deleting the sialic acid transporter yielded cells with no nucleotide-activated sialic acid in the secretory pathway, yet ample CMP-Neu5Ac in the rest of the cell, which would be available to sialyltransferases post-lysis but not when intact. In line with known results^24^, SLC35A1^-/-^ KO resulted in loss of cell surface sialylation (Fig. 1E), with a corresponding increase of galactose-terminated structures. We extend these observations by demonstrating a similar impact on nuclear sialylation (Fig. 1E), suggesting that nuclear glycoproteins are assembled in living cells, prior to lysis, and that their biosynthesis requires the secretory pathway.

Following up on this observation, we then set out to image nuclear glycans in intact cells. To this end, we took advantage of metabolic labeling techniques for sialic acid^25^ and peptide localization. We synthesized a probe peptide with cell-penetrating and nucleus-homing properties^26^ (N50-sC18*), functionalized with a DBCO moiety for copper-free click chemistry^25^ and a biotin handle for detection and affinity purification (Biotin-DBCO-N50-sC18*, Supplementary Methods). We metabolically labeled the sialosides in A375 cells with Ac_4_ManNAz, incorporating an azide into sialic acid (e.g., Neu5Az). A375 cells were chosen because the RENBP epimerase enzyme, which generates GlcNAc/Az from ManNAc/Az^27^ is generally not expressed in A375 cells^28^. In line with this, we could not detect the enzyme (Fig. S6), precluding labeling due to *O*-GlcNAc.

Using our nuclear-selective probe, we visualized extended nuclear glycans in intact cells (Fig. 1F). Quantifying the nuclear peptide signal over 48 hours then allowed us to show that nuclear signal was rapidly washed out under normal culture conditions yet was retained when cells were cultured with ManNAz (Fig. 1G, Fig. S7, Fig. S8), indicating successful reaction with nuclear glycoproteins. To ensure that our peptide indeed reacted with proteins, and not CMP-sialic acid, we next performed a streptavidin immunoprecipitation of this nuclear material and indeed detected many labeled proteins via Western blot (Fig. S9A).

To identify the sialylated nuclear proteins that were reacting with our designer peptide, we performed streptavidin-based affinity purification, followed by proteomics analysis of nuclei from DMSO control- and Ac_4_ManNAz-treated cells labeled with the peptide. This revealed 683 significantly different nuclear proteins between these two conditions (Fig. 1G, Fig. S9B, Table S1, Table S2), with 602 enriched nuclear proteins. Depleted proteins included cell adhesion and extracellular matrix (ECM)-receptor interaction proteins, known to be downregulated in the presence of ManNAz^29^, whereas enriched proteins reacting with our click-peptide included many RNA-binding proteins, with the most enriched protein being RPP30^30^, a soluble, nucleolar component of ribonuclease P (RNase P). Analyzing RPP30 with *O*-glycosylation site predictors such as NetOGlyc 4.0^31^ indeed yielded strong predicted glycosylation sites (up to a score of 0.87 on S55).

Following up on our results, we then validated our top hits (RPP30 and the transcription factor BBX) via Western blot in a fresh experiment (Fig. S10). We also ensured that our results do not stem from nonenzymatic Ac_4_ManNAz-driven *S*-glycosylation by repeating the peptide labeling and validation pull-downs using cells that were metabolically labeled with non-acetylated ManNAz (Fig. S11). Given that (i) peptide labeling occurred within living cells under physiological conditions and (ii) required not only glycosylated but sialylated proteins, we thus present these enriched proteins as the first map of the complex *O*-glycoproteome in the nucleus.

### Glycomics and enriched proteomics confirm extended *O*-glycans inside the nucleus

After demonstrating the presence of nuclear glycoproteins in living cells, we next set out to determine which exact glycan sequences could be found inside nuclei. For this, we used ultra-pure A375 nuclei (Fig. 1B, Fig. S1) and performed mass spectrometry-driven glycomics experiments. Confirming our earlier lectin-based experiments, we identified several extended *O*-GalNAc type glycans in A375 nuclei (Fig. 2A, Table S3), which we confirmed in HEK293T nuclei (Fig. S12, Table S4). While we observed core 1 *O*-glycan structures in both cell lines, only A375 exhibited substantial amounts of core 2 *O*-glycans. Further, A375 nuclei exhibited a stronger nuclear glycosylation signal than HEK293T nuclei overall, indicating cell type-specific differences in nuclear glycosylation patterns (Fig. 1C, Fig. S4). These structures are traditionally located in the trans-Golgi, far away from the nucleus, and hence are unlikely to stem from contaminants. Moreover, the absence of any detected mature (complex/hybrid) *N*-glycans further reinforces the idea of a distinct glycome pool and rules out the possibility of contamination.

**Figure 2.**
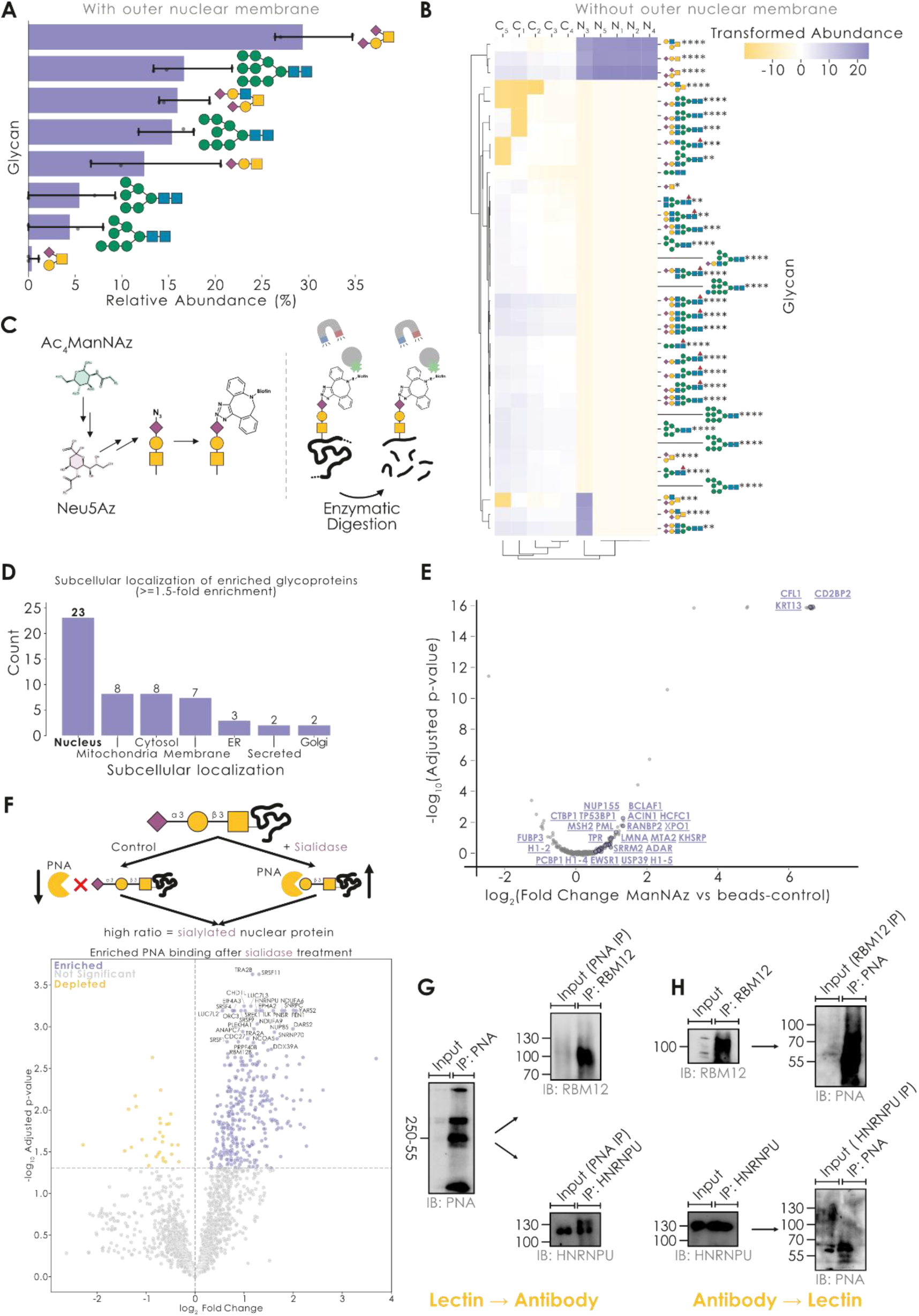
Strongly sialylated O-GalNAc glycans modify nuclear proteins. A) Glycomics of purified A375 nuclei (n = 3) revealed extended, sialylated O-GalNAc type glycans, yet no complex/hybrid N-glycans. B) For O-glycomics measurements of the secretory pathway (C) and ultra-pure nuclei with removed outer nuclear membrane (N) of A375 cells, we identified different sets of glycans in the different compartments (n = 5). Data are visualized via hierarchical clustering of CLR-transformed relative abundances using the get_heatmap function of glycowork32 (v1.5.0), followed by establishing differential expression via the get_differential_expression function of glycowork, indicated for each structure. C) Workflow for glycoprotein-enriched proteomics. D) Major subcellular localization, based on UniProt annotations, of glycoproteins enriched in ManNAz-treated nuclei (at least 1.5-fold enrichment compared to nuclei from cells grown without ManNAz, n = 1). E) Volcano plot of glycoprotein enrichment in nuclei after ManNAz treatment, with enriched nuclear proteins labeled in purple. F) Volcano plot of PNA capture enrichment after sialidase treatment, with enriched nuclear glycoproteins labeled in purple. G-H) Nuclear glycoproteins can be purified from human cells via tandem immunoprecipitation (IP). For the RNA-binding proteins RBM12 and HNRNPU, we show Western blots stained with either PNA or anti-RBM12/anti-HNRNPU. Lanes include both the elution from the first IP (anti-RBM12 or PNA) as well as the elution from the second IP. Tandem IPs are shown as lectin-then-antibody (G) and antibody-then-lectin (H). *p < 0.05, **p < 0.01, ***p < 0.001, ****p < 0.0001.

To further validate our glycomics results from A375 nuclei (Fig. 2A), we generated more stringently purified nuclei by removing the outer nuclear membrane (Fig. 2B, Fig. S1). This approach eliminated the mannose-rich, immature *N*-glycans characteristic of the ER, while, importantly, retaining the extended *O*-glycans, which seemed to be located inside the nucleus. We next compared the nuclear A375 *O*-glycome to its canonical secretory pathway glycome from paired samples, revealing a distinctly different glycome in the two locations (Fig. 2B, Table S5). We therefore conclude that the mammalian nucleus harbors a separate pool of glycans, namely extended *O*-GalNAc type glycans rich in sialic acid, with a particular enrichment of sialylated core 1 structures, such as the disialyl T antigen.

To better understand the functional implications of *O*-GalNAc glycosylation in the nucleus, we next continued to identify which nuclear proteins were post-translationally modified. We returned to a click chemistry-based enrichment strategy for this to facilitate enrichment (Fig. 2C). In contrast to the indirect peptide capture described above, we here aimed to directly biotinylate nuclear glycans, to avoid any peptide-specific effects. For this, we fed A375 cells with Ac4ManNAz to extract nuclei with Neu5Az-modified glycans, which we then directly reacted with a UV-cleavable biotin tag. Streptavidin-based enrichment then enabled the specific isolation of nuclear proteins carrying sialylated glycans. Since proteins can bind non-specifically to streptavidin beads, we compared this to a control of identically treated nuclei from cells not grown with Ac4ManNAz. Proteomics of enriched glycoproteins then identified 52 proteins with substantial enrichment (>1.5-fold) in Ac4ManNAz-treated nuclei compared to controls (Fig. 2D-E, Fig. S13, Table S6), including hits such as BCLAF1, TP53BP1, as well as various histones.

As an orthogonal strategy, we also compared the PNA-driven capture of nuclear glycoproteins with and without prior sialidase treatment (controlling for any unspecific PNA binding that should be invariant between conditions), and report 330 nuclear proteins that significantly increased in PNA capture after sialidase treatment (exposing the Galβ1-3GalNAc PNA binding epitope; Fig. 2F, Table S7, Table S8). Notably, this included many RNA-binding proteins, such as RBM12B, PRPF40B, or HNRNPU. We then went on to show that these glycosylated RBPs could be purified from human nuclei via tandem immunoprecipitation of RBM12/HNRNPU and the lectin PNA (Fig. 2G-H), which notably succeeded in both directions (antibody followed by lectin, and vice versa). We then validated additional hits from our PNA capture enrichment (Fig. 2F), such as TRA2B and NUP85, via such tandem IPs (Fig. S14).

Reasoning that published *O*-glycoproteomics measurements of whole cells or tissues may have inadvertently detected nuclear glycopeptides, which would have been ignored under the current paradigm, we curated a dataset of noncanonical glycopeptides (Table S9). This revealed thousands of non-secretory pathway glycopeptides, often from the same proteins or even the same sites measured repeatedly in different studies, which carried extended glycans (Fig. S15). Analyzing the sequence contexts of identified sites revealed strong similarities to canonical *O*-glycosylation sites (Fig. S16), which was in line with our own findings with examples such as RPP30 above. Reasoning that regulatory processes such as glycosylation might be enriched in particular biological processes, we then engaged in a gene set enrichment analysis of this dataset and revealed that RNA-related functions were strongly enriched among reported nuclear glycoproteins (Fig. S17). These results, aggregated from dozens of peer-reviewed studies and our own data, clearly show that extended nuclear glycosylation is a recurring phenomenon and that at least some of the sites seem remarkably conserved, indicating functional implications.

Given this consistent enrichment of RNA-binding proteins across our datasets, we next designed an experiment to probe the impact of glycosylation on a representative example: RBM12, a RBP implicated in cancer immune evasion^33^ and one of the enriched proteins identified in our datasets, which further displays distinct nuclear localization by immunofluorescence microscopy (Fig. S18A). This validation was designed as a co-immunoprecipitation between RBM12 and PNA as a lectin binding extended *O*-glycans (Fig. S18B). While the co-IP of RBM12 and PNA only yielded a modest increase in signal compared to controls, the addition of sialidase (cleaving sialic acid and thus exposing the PNA binding motif) substantially increased the signal of the RBM12-PNA interaction. This again indicated that RBM12 exhibited sialylated extended *O*-glycans in human nuclei, paving the way for studying functional implications of this in the future. We also note that only the nuclear pool of RBM12 was glycosylated, and not the cytoplasmic pool that was also detectable via Western blot, suggesting either subcellular selection for glycosylation or a role for glycosylation in regulating subcellular localization.

### Extended nuclear *O*-glycans are biosynthesized in the Golgi apparatus

A central question arising from the discovery of extended *O*-glycans in the nucleus concerns the mechanism of their biosynthesis, especially given that most of the modified nuclear proteins we identified here lack a traditional signal peptide. Results from our HEK293T*^SLC35A1-/-^*cell line in which the CMP-Neu5Ac transporter SLC35A1—required for sialylation in the Golgi—is genetically disrupted, clearly showed that nuclear sialylation levels were dependent on secretory pathway sialylation (Fig. 1E). Next, we aimed to further confirm this with other steps of the glycan biosynthesis pathway, probing whether the canonical biosynthesis machinery for structures such as the disialyl-T antigen (Fig. 3A) also constructed the glycans we observed in nuclei.

**Figure 3.**
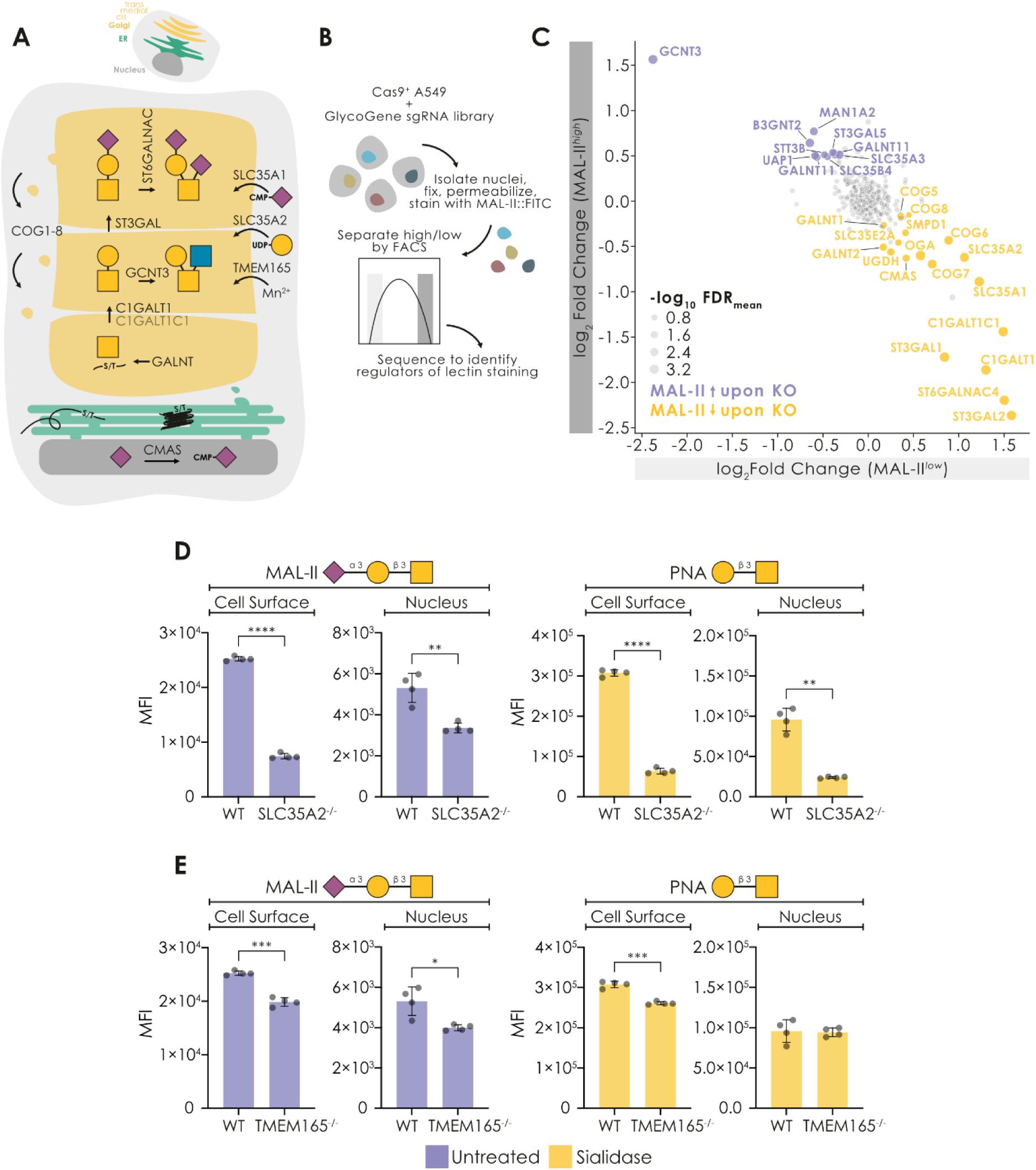
Nuclear glycans are synthesized in the Golgi apparatus. A) Schematic view of the canonical biosynthesis of the disialyl-T antigen, including relevant enzymes and transporters. B) Schematic view of the nuclear GlycoGene CRISPR/Cas9 KO screen workflow. C) Results of the glycosylation-focused CRISPR/Cas9 KO screen. Shown are the significant genes for which the KO either resulted in lower or higher MAL-II signal in A549 nuclei (n = 2). D-E) HEK293T cells with a knocked-out galactose transporter (HEK293TSLC35A2-/-; n = 4; D) or Mn2+ transporter (HEK293TTMEM165-/-; n = 4; E) show decreased binding of the sialic acid- and galactose-binding lectins MAL-II and PNA to their cell surface and nuclei, supporting secretory pathway biogenesis of nuclear glycans. Subset of Fig. S20. Significant differences were established via a Welch’s t-test. *p < 0.05, **p < 0.01, ***p < 0.001, ****p < 0.0001.

To this end, we performed a focused CRISPR/Cas9 knockout (KO) screen, using MAL-II staining of nuclei as an output signal (Fig. 3B). MAL-II binds the terminal Neu5Acα2-3 motif on *O*-glycans^23^, thus it can serve as a reporter for genes relevant for the vast majority of nuclear glycosylation (e.g., genes relevant for the construction of the core 1 structure will still affect the level of Neu5Acα2-3 and hence are detectable with this approach). We created an A549 CRISPR/Cas9 KO library focused on 347 glycosylation-related genes using an established guide library^34^. Sorting nuclei based on MAL-II staining, we identified multiple enzymes and Golgi proteins responsible for the nuclear *O*-glycome (Fig. 3C, Fig. S19A-D, Table S10, Table S11).

As expected, the sialic transporter SLC35A1 was among our top hits, confirming our previous data (Fig. 1E). In addition, we observed the galactose transporter SLC35A2. Consistent with our screen results, staining of the previously established^35^ HEK293T*^SLC35A2-/-^* cell line showed substantial decreases in the binding of both sialic acid- and galactose-specific lectins to nuclei and cell surface (Fig. 3D, Fig. S20A).

MAL-II binding also heavily relied on the isozymes ST3GAL1/2, α2,3-sialyltransferases that glycosylate *O*-glycans and which have known overlaps in their substrate specificity^36^. Further, the strict reliance on the core 1 biosynthesis machinery (C1GALT1 and its chaperone C1GALT1C1) reinforces our interpretation that the detected structures are *O*-GalNAc type glycans.

We also note that the entire lobe B of the COG complex^37^ (COG5-8), which is associated with Golgi-traversing vesicles^38^, was identified as hits here. COG defects are known to be associated with generally impaired glycosylation, e.g., in the context of COG-CDGs (congenital disorders of glycosylation)^39^. In order to differentiate between general vs nuclear-specific effects, we repeated the CRISPR/Cas9 KO screen but this time sorted based on cell surface, rather than intra-nuclear, MAL-II binding. Here, we still identified all expected hits related to *O*-GalNAc glycosylation, i.e., C1GALT1, C1GALT1C1, ST3GAL1/2, and SLC35A1/2, yet none of the COG components were significantly enriched (Fig. S19E-H, Table S12, Table S13).

In addition, several other studies applying the same GlycoGene CRISPR/Cas9 screen to lectin-binding of sialylated *O*-GalNAc glycans on the cell surface (i.e., E-selectin^40^, P-selectin^40^, HECA-452^40^, and ST3GAL-sCore2^41^) failed to find COG components. Further supporting this selectivity, siRNA-mediated knock-down of COG7 resulted in significantly decreased MAL-II and PNA staining in HEK293T nuclei, while we observed increased staining intensity at the cell surface (Fig. S21). This implicates a role for retrograde vesicular trafficking in nuclear glycosylation.

While the GlycoGene-focused screen served as a very sensitive instrument to probe nuclear glycan biosynthesis, we reasoned that genes beyond the 347 assayed could also contribute to this phenomenon. To demonstrate this with a gene absent from our CRISPR screen, we turned to an orthogonal approach to probe biosynthesis by examining a HEK293T*^TMEM165–/-^* cell line. TMEM165 encodes a Golgi-resident transmembrane protein that regulates manganese (Mn^2+^) homeostasis in the secretory pathway, where Mn^2+^ serves as an essential co-factor for many glycosyltransferases^42^. Loss of TMEM165 disrupts glycosylation by impairing the function of these enzymes, and we indeed observed a decrease in nuclear glycosylation levels in these cells, although less pronounced than those seen in the transporter knockouts (Fig. 3E, Fig. S20B). Overall, both the KO screen and our validation strongly support our initial finding that extended nuclear glycans are assembled in the secretory pathway, in particular within the Golgi apparatus.

### Nuclear glycoproteins are trafficked by a novel Golgi-nucleus transport pathway

Since nuclear glycan biosynthesis was Golgi-dependent, we next set out to show that nuclear proteins themselves were indeed transiting through the Golgi apparatus. For this, we used the well-established APEX2 proximity labeling system^43^ to transiently biotinylate proximal proteins in live cells for subsequent enrichment and characterization via proteomics. We used the established PAQR3-APEX2^44^ as a marker for the cis-Golgi proteome and constructed a trans-Golgi marker with the ST6GAL1-APEX2 construct. In both cases, the APEX2 protein was located in the lumenal part of the Golgi apparatus, i.e., any biotinylated protein would also need to be inside the Golgi (Fig. 4A, Fig. S22). Enriching biotinylated proteins via streptavidin beads (vs control samples which had not been biotinylated) revealed 834 statistically significantly enriched nuclear proteins transiting through the Golgi (Fig. 4B-C, Table S14, Table S15).

**Figure 4.**
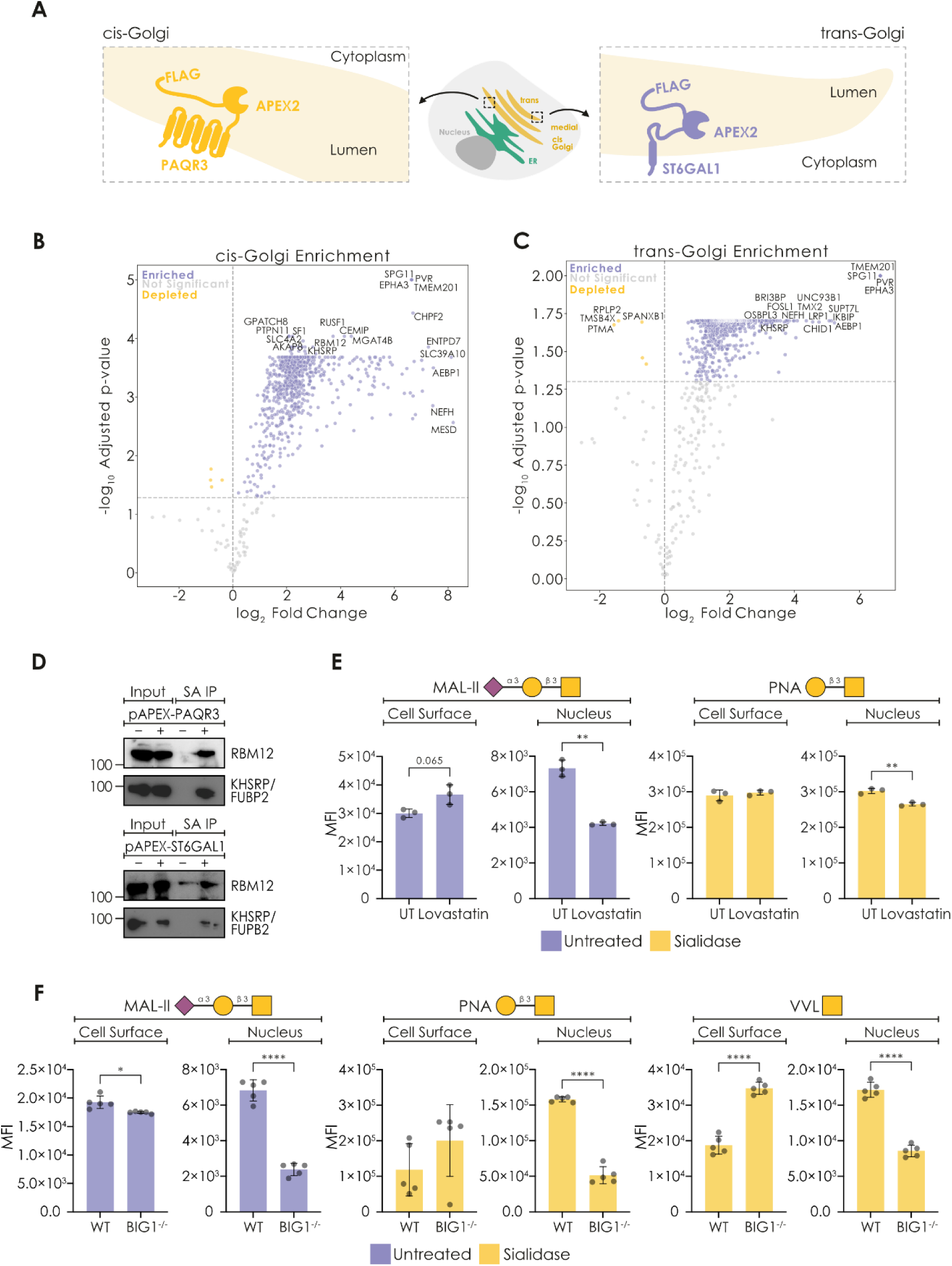
Nuclear glycoproteins are present in the Golgi apparatus and transported to the nucleus. A) Schematic view of the cis-Golgi marker PAQR3-APEX2 and the trans-Golgi marker ST6GAL1-APEX2. B-C) Tagging Golgi-resident proteins in live cells reveals nuclear proteins in the secretory pathway. PAQR3-APEX2 (B; cis-Golgi) and ST6GAL1-APEX2 (C; trans-Golgi) mediated biotinylation shows Golgi-enrichment of hundreds of proteins associated with the nuclear proteome (ncontrol = 3, nPAQR3-APEX2 = 5, nST6GAL1-APEX2 = 4). D) Western blots of streptavidin-IPs for the nuclear proteins KHSRP/FUBP2 and RBM12 after PAQR3-APEX2 and ST6GAL1-APEX2 mediated biotinylation. SA: streptavidin E) Disrupting vesicular transport from the trans-Golgi with lovastatin reduced nuclear glycosylation, yet did not impair surface glycosylation in HEK293T cells (n = 3). Subset of Fig. S26. F) Knocking out BIG1-dependent vesicular transport selectively abrogates nuclear glycosylation. Flow cytometry of HEK293T and HEK293TBIG1-/- cells/nuclei, stained with MAL-II, PNA, and VVL (n = 5). Subset of Fig. S27. Significant differences were established via a Welch’s t-test. *p < 0.05, **p < 0.01, ***p < 0.001, ****p < 0.0001.

We observed an exceptionally high concordance between nuclear cis-Golgi and trans-Golgi hits (86%; Fig. S23A), further supporting the conclusion that nuclear proteins transited the cis-Golgi and were processed up to the trans-Golgi, consistent with their glycosylation via the canonical biosynthesis machinery. Further, we could show that 74% of these nuclear proteins had been previously^45^ detected in an APEX-like ER screen using a peroxidase that was exclusively active in the secretory pathway (Fig. S24, Table S16), arguing for an ER entrance and/or exit via retrotranslocation for these nuclear proteins. This last point was also supported by our glycosylation-focused KO screen, since the KO of MAN1A2 (mannosidase alpha class 1A member 2)—increasing high-mannose *N*-glycans and subsequent ERAD-retrotranslocation—led to higher MAL-II staining in our nuclei (Fig. 3C). Additionally, we noted a substantial overlap between nuclear proteins identified in the Golgi apparatus and nuclear glycoproteins identified via our Biotin-DBCO-N50-sC18* peptide-aided mass spectrometry (Fig. S23B), further affirming our results and mapping the full route of nuclear glycosylation. Subsequent validation of well-studied nuclear RBPs, with the example of KHSRP/FUBP2 and RBM12 (enriched in both APEX2 experiments and click chemistry-aided mass spectrometry), indeed confirmed their co-localization with Golgi-resident APEX2 proteins via Western blots (Fig. 4D), while the main pool of KHSRP/FUBP2 still resided in the nucleus (Fig. S25) and nuclear RBM12 can be shown to be glycosylated (Fig. S18).

To find out whether there was a distinguishing feature of the nuclear proteins we identified in the secretory pathway, we then assembled a set of 1,214 multi-localizing nuclear proteins (Table S17) from across our datasets, to show that this comprised a characteristic subset of the entire nuclear proteome (Table S18). This also revealed that, even among nuclear proteins, our multi-localizing proteins were still enriched for RNA-binding proteins (p < 0.001, Fisher’s exact test). We also report a significant enrichment of Ca^2+^-binding EF hand motifs (p < 0.001, Fisher’s exact test) and ATP/GTP-binding P-loops (p < 0.001, Fisher’s exact test) in multi-localizing nuclear proteins, compared to the general pool (Table S19), arguing for specific regulation that recruits certain types of nuclear proteins to traverse the secretory pathway.

For nuclear proteins to be glycosylated in the secretory pathway, they have to not only enter this compartment but also return to the nucleus as their primary site of localization. We thus probed this trafficking, beginning with the Golgi-nucleus transport step, as we could here rely on previous studies that have shown initial evidence of exchange between the trans-Golgi region and the nucleus^16^, particularly regulated by BIG1^17,18^, a key factor in vesicular protein sorting. Leveraging the known impact of small molecule inhibitors on BIG1 functioning^17^, we first disrupted vesicular transport from the trans-Golgi using inhibitors such as lovastatin (inhibiting the prenylation of small GTPases), which left cell surface glycosylation mostly intact yet reduced nuclear glycosylation levels (Fig. 4E, Fig. S26). This could be partly rescued by mevalonate addition, which is the metabolite from the step after lovastatin inhibition (Fig. S26). Together, this data strongly indicates that nuclear glycoproteins are actively transported from the trans-Golgi via vesicles in a GTPase-dependent manner.

Hypothesizing the key role of BIG1 in this transport process, we next produced a HEK293T*^BIG1-/-^* KO cell line to disentangle the surface and nuclear glycomes (Fig. 4F, Fig. S27). Indeed, for commonly observed terminal epitopes of nuclear glycans (MAL-II: Neu5Acα2-3, PNA: Galβ1-3GalNAc, VVL: GalNAc), we observed significant decreases in HEK293T*^BIG1-/-^* KO in nuclei yet not on the cell surface, supporting our hypothesis that active transport from the trans-Golgi is key in establishing and maintaining the nuclear glycome.

### Glycosylation of RPP30 impacts tRNA processing

To determine whether glycosylation of nuclear proteins impacted function, we focused on the intranuclear, tRNA-processing protein RPP30, we mutated its hypothesized glycosylation site (RPP30^S55A^) and expressed both WT and mutated version as FLAG-tagged proteins in A375 cells. While RPP30^S55A^ still correctly localized to the nucleus (Fig. S28), we could show that it exhibited decreased glycosylation compared to RPP30^WT^ (Fig. 5A), which resulted in a reduced capacity to bind tRNAs (Fig. 5B-C, Table S20-21). This led to a global reduction of protein synthesis, measured via a SUnSET assay^46^ (Fig. 5D, Fig. S29), indicating that RPP30^S55A^ acted as a dominant negative and that glycosylation at Ser55 is crucial to facilitate effective tRNA processing. We thus identify RPP30 as the first protein- and site-specific function for extended nuclear glycosylation.

**Figure 5.**
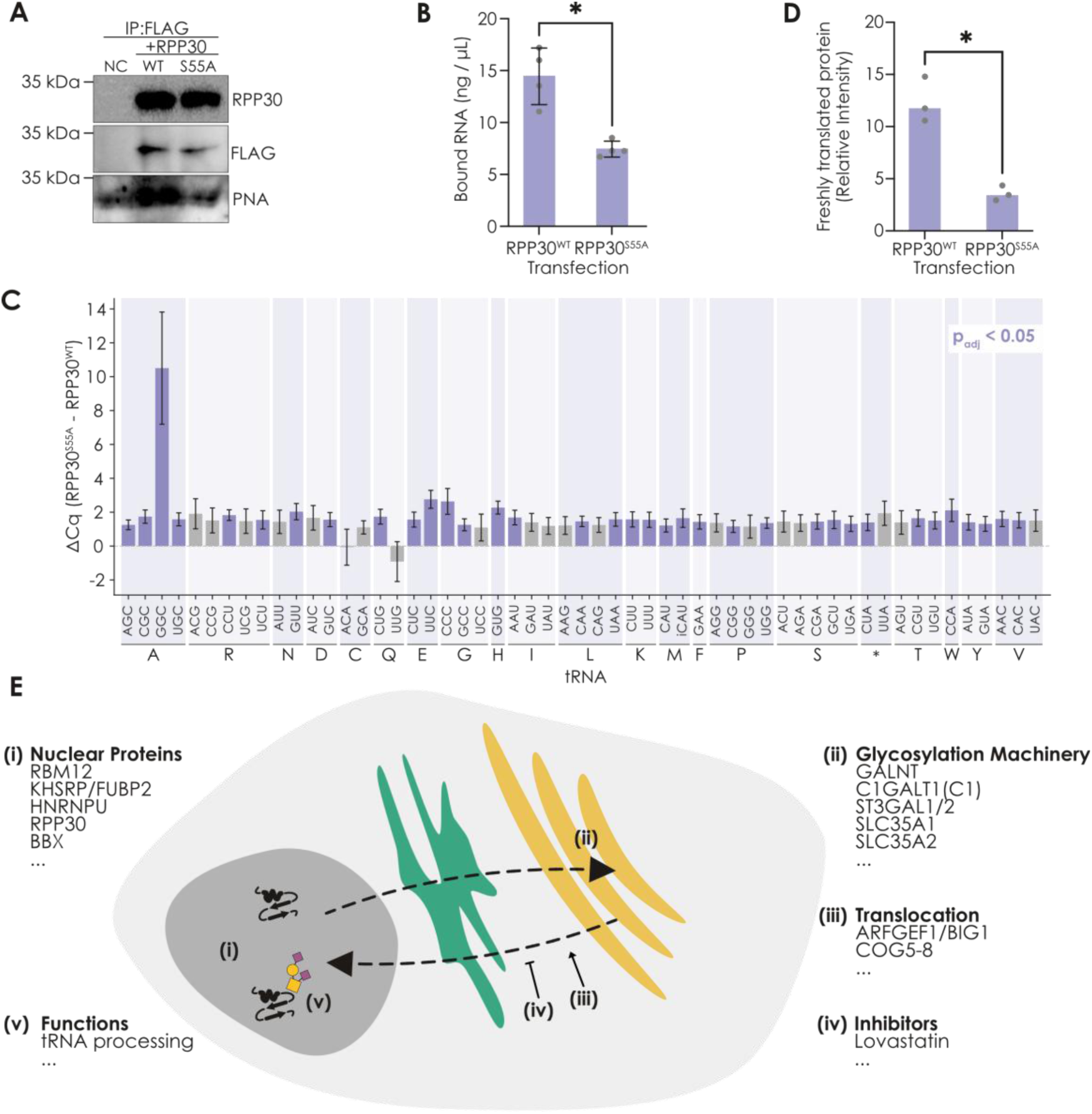
Extended nuclear glycosylation regulates tRNA processing. A) RPP30 is glycosylated at Ser55. Shown is a Western blot of an anti-FLAG immunoprecipitation from nuclear lysates from FLAG-tagged RPP30WT and RPP30S55A-expressing A375 cells, probed with either anti-RPP30, anti-FLAG, or PNA. B) Decrease in total RNA binding of RPP30S55A. Shown are bar graphs with overlaid scatter plots and error bars indicating standard deviation of Qubit-quantified RNA bound to immunoprecipitated RPP30WT and RPP30S55A. C) Deglycosylated RPP30 binds less tRNAs. Shown are bar graphs of the Cq difference with error bars indicating standard error of the difference of qRT-PCRs of human tRNAs in RNA from anti-FLAG immunoprecipitation from whole cell lysates from FLAG-tagged RPP30WT and RPP30S55A-expressing A375 cells. Significant differences after correction are colored purple. D) Deglycosylated RPP30 stymies global protein synthesis. From the same overexpression set-up as in (C) we quantified the amount of freshly translated proteins with a SUnSET assay46. Shown is a bar graph with overlaid scatter plot of anti-puromycin signal intensity on a Western blot for RPP30WT and RPP30S55A overexpression. E) Current working model of the production and trafficking of nuclear glycoproteins. *p < 0.05.

Overall, we conclude that our findings frame a new post-translational modification mechanism, in which nuclear proteins (after being translated) are transported into the Golgi apparatus, glycosylated, and returned to the nucleus via active vesicular transport (Fig. 5E).

## Discussion

Herein, our work demonstrates that extended *O*-linked glycans are a common and conserved class of post-translational modifications inside mammalian nuclei. Multiple methods as well as numerous controls incontrovertibly show that mature *O*-linked, but not mature *N*-linked, glycans can be identified on nuclear proteins, especially RNA-binding proteins. While some steps are not yet fully clarified, our data convincingly show that the biosynthesis of extended nuclear glycans occurs in the secretory pathway, coupled with novel transport routes between the Golgi and nucleus. This work leads to a deeper understanding of cellular physiology and paves the way for a great number of follow-up studies.

Several reasons exist for why this class of PTMs has been largely overlooked on nuclear proteins so far. First, *O*-glycans are harder to analyze than *N*-glycans, due to the lack of a simple enzyme removal step such as PNGaseF. Thus, much of the work in the field has focused on the *N*-glycome^47^. Second, based on fluorescence intensity in our flow cytometry experiments, we note that sialylated glycans appear less abundant in the nucleus than in the secretory pathway. We speculate that this stems from either a lack of densely glycosylated mucin domains in the nucleus or a low stoichiometry of occupied glycosylation sites, for the glycosites we identify here. This latter situation is common for *O*-GlcNAc sites, where this is still physiologically relevant.

While we did not do so explicitly, we urge future research to probe whether the conservation of nuclear *O*-glycans extends beyond mammals. If that is so, it will be exciting to see whether nucleocytoplasmic *O*-Fuc glycans, an analog of *O*-GlcNAc used in some non-mammalian eukaryotes^48^, could contribute to extended nuclear glycans, as they do in the case of thrombospondin repeats in the secretory pathway^49^.

Overall, we cannot help but notice that the strong enrichment of extended nuclear *O*-glycans on RNA-binding proteins mirrors the recent discovery of glycosylated RNA^50^—in more ways than one—and further ties the RNA world and glycobiology together. This also extends to our functional investigation of the RNA-processing RPP30. As a member of not only the tRNA-processing RNase P but also the rRNA-processing RNase MRP complex^51^, it indicates the impact of nuclear glycosylation on several types of RNA and could provide additional explanatory factors for the impact of RPP30^S55A^ on cellular translation.

We also again point out the striking enrichment of lobe B of the COG complex in our CRISPR/Cas9 KO screen. Previous research has found extremely symmetric impacts on surface glycosylation when knocking out members of lobe A or B^52^, arguing for a lobe B-specific role in the case of nuclear glycans.

Further, active exchange between the nucleus and the cytosol leads us to speculate that non-canonically extended *O*-glycans may not be restricted to the nucleus and may eventually be identified in other cellular compartments.

## Limitations of the study

Given the lower abundance of extended nuclear glycans, as well as potential sample loss during nuclear extraction, we likely only sample high-abundance glycoproteins/glycopeptides here and caution that this does not constitute the entire glycoproteome of the nucleus. Further, we cannot conclusively rule out the presence of additional, low-abundance, glycans with other sequences than those described here, such as neutral *O*-glycans with fucosylation and/or extended core structures.

## Methods

### Resource availability

#### Lead contact

Further information and requests for resources and reagents should be directed to and will be fulfilled by the Lead Contact, Daniel Bojar

#### Materials availability

All unique/stable reagents generated in this study are available from the Lead Contact.

#### Data and code availability

The glycomics MS raw files have been deposited in the GlycoPOST database under https://glycopost.glycosmos.org/preview/96766034968ba8857c0ebf, code 6584. Proteomics raw files have been deposited to the ProteomeXchange Consortium (http://proteomecentral.proteomexchange.org) via the PRIDE partner repository^53^ under the dataset identifiers PXD069188 and PXD069531. Curated literature glycoproteomics data and processed results from glycomics, proteomics, and other measurements can also be found in the supplementary tables. Any additional information required to reanalyze the data reported in this paper is available from the lead contact upon request.

### Experimental model and subject details

#### Mammalian cell culture

A375, MCF-7, A549, HeLa, HEK293, HEK293T, HEK293T*^BIG1-/-^*, HEK293T*^SLC35A1-/-^*, HEK293T*^SLC35A2-/-^*, and HEK293T*^TMEM164–/-^* were cultured in high-glucose Dulbecco’s Modified Eagle Medium (DMEM; Gibco, 11995065). SH-SY5Y and THP-1 cells were maintained in RPMI 1640 medium (Gibco, 11875093). HCT116 cells were cultured in McCoy’s 5A medium (Gibco, 11540646). SK-N-BE(2) cells were cultured in DMEM F:12 media (Gibco, 11554546). The respective growth media was supplemented with 10% heat-inactivated fetal bovine serum (FBS; Gibco, 11573397) and 100 U/mL Penicillin-Streptomycin (Gibco, 15140122). AG21708 cells were cultured in Eagle’s Minimum Essential Medium (EMEM) with Earle’s salts and non-essential amino acids with 2 mM L-glutamine (M0643, Sigma-Aldrich) supplemented with 15% FBS. E14 embryonic stem cells were cultured on 0.1% gelatin-coated plates with DMEM (Thermo Fisher, 12430) supplemented with 15% ESC-qualified fetal bovine serum (Thermo Fisher, SH30070.03), 1 mM sodium pyruvate (Thermo Fisher, 11360070), 1×non-essential amino acids (Gibco, 12599049), 1×L-glutaMAX (Gibco, 11500626), 100 µM 2-mercaptoethanol (Thermo Fisher, 21985023), and 1000 U/mL of a leukemia inhibitory factor (Sigma-Aldrich, ESG1107). All cells were maintained at 37°C in a humidified incubator with 5% CO_2_ and cultured in their respective growth media.

### Method details

#### Preparation of whole cell lysate

Cell pellets were resuspended in RIPA Lysis Buffer (150 mM NaCl, 5 mM EDTA pH 8.0, 50 mM Tris, 0.1% (v/v) NP-40, 0.5% Sodium Deoxycholate, 0.1% (v/v) SDS) supplemented with protease inhibitors. The lysates were incubated on ice for 30 minutes and then sonicated once at 60% amplitude for 15 seconds (Sonics, VCX130). The lysates were subsequently centrifuged at 21,300×*g* for 15 minutes at 4°C. The supernatant was then transferred to a new tube and stored at -20°C until further analysis.

#### Isolation of highly pure nuclei

The cell pellet was resuspended in Hypotonic Lysis Buffer (HLB; 10 mM Tris-HCl, 10 mM NaCl, 3 mM MgCl_2_, 0.6% (v/v) NP-40, and 10% (v/v) glycerol), supplemented with protease inhibitors. Following incubation, the sample was centrifuged at 200×*g* for two minutes at 4°C, and the supernatant, representing the cytoplasmic fraction, was carefully collected into a fresh tube for downstream applications. The remaining pellet, representing the nuclear fraction, was washed three times with HLB buffer, each time followed by centrifugation at 200×*g* for two minutes at 4°C.

For standard nuclear fractionation, the pellet was subsequently subjected to an additional wash and the pellet was then resuspended in Nuclei Lysis Buffer (NLB; 20 mM Tris-HCl, pH 7.5, 150 mM KCl, 3 mM MgCl_2_, 0.3% (v/v) NP-40, and 10% (v/v) glycerol), supplemented with protease inhibitors.

For preparation of the nuclear fraction lacking the outer nuclear membrane, the washed nuclear pellet was incubated in 2% sodium citrate for one hour at 25°C with gentle rotation at 300 rpm. Following incubation, samples were then centrifuged at 1200×*g* for 10 minutes at 4°C, and the supernatant containing the outer nuclear membrane was discarded. The resulting pellet was resuspended in NLB as described above.

In both protocols, equal volumes of HLB and NLB were used to ensure a consistent cytoplasmic-to-nuclear ratio. The nuclear fractions were then subjected to sonication at 60% amplitude for 15 seconds (Sonics VCX130). Cytoplasmic and nuclear samples were subsequently centrifuged at 21,300×*g* for 30 minutes at 4°C. The resulting supernatants were collected and stored at –20°C until further analysis.

#### Western blotting

Protein quantification of cell lysates was performed using the Pierce 660 nm Protein Assay (Thermo Fisher, 10177723) according to the manufacturer’s instructions. Protein lysates were denatured at 95°C for five minutes with 1×Laemmli SDS reducing reagent (Thermo Fisher, 15493939). Protein samples were separated by SDS-PAGE on 10% polyacrylamide gel cast using the Invitrogen SureCast gel system. Electrophoresis was carried out at 120 V until completion, followed by transfer to a nitrocellulose membrane using a semi-dry transfer system at 400 mA for 50 minutes. The membrane was blocked with 2% skim milk in Tris-buffered saline with Tween-20 (TBST; 20 mM Tris-HCl, 150 mM NaCl, 0.1% Tween 20) for 20 minutes. Membranes were incubated with primary antibodies specific to the target proteins or lectins, followed by washing three times with TBST for 10 minutes each. Appropriate secondary antibodies or streptavidin were applied, and the membrane was developed using SuperSignal chemiluminescent HRP substrates (Thermo Fisher, 11859290) and visualized using a ChemiDoc Imaging System. Antibodies and lectins are listed in Table S22.

#### Flow cytometry

For all experiments, live cells and isolated nuclei from the same batch of cells were compared. After harvesting cells, an aliquot of live cells was kept on ice during the isolation and fixation of nuclei. Intact, pure nuclei were obtained by resuspending a cell pellet in Homogenization Buffer (HB; 250 mM sucrose, 25 mM KCl, 5 mM MgCl_2_, 10 mM Tris buffer pH 8.0, 0.1% (v/v) Triton X-100) followed by incubation on ice for five minutes with gentle mixing. Nuclei were pellet by centrifugation at 1000×*g* for eight minutes at 4°C and washed in 1×PBS. Nuclei were fixed in Fixation/Permeabilization solution (00-5223-56 and 00-5123-43, Thermo Fisher) for 15 minutes at room temperature and washed twice and resuspended in Permeabilization Buffer (PERM; 00-8333-56, Thermo Fisher). α2-3/6/8-Sialidase (10269611001, Sigma-Aldrich) treatment was carried out using 0.01 U sialidase / 50 µL buffer / sample in PBS (live cells) or PERM (fixed nuclei) with 5 mM sodium acetate pH 5.2 (S7899, Sigma-Aldrich) for 60 minutes at 37°C. Cells and nuclei were incubated with 10 µg/mL lectin solution and 2 µg/mL streptavidin-fluorescein in PBS or PERM, respectively, for 20 minutes. Lectins are listed in Table S22. For live cells, surface staining was done together with LIVE/DEAD Fixable Far Red Dead Cell Stain Kit (L10120, Thermo Fisher, 1:1000 dilution). Data were acquired on an Accuri C6 Plus Flow Cytometer (BD) and analyzed using FlowJo 10.10.0 software.

#### Generation of APEX2 constructs

Protein expression plasmids for FLAG-APEX2-PAQR3 and FLAG-APEX2-ST6GAL1 were generated by subcloning into the pcDNA5-FRT/TO-3×Flag-APEX2 PML plasmid (Addgene #187583) using the Takara In-Fusion^®^ HD Cloning Kit, according to the manufacturer’s protocol. The corresponding gene fragments were PCR-amplified from cDNA using the primers listed in Table S22. APEX2-fusion plasmid constructs were expressed in Stellar™ *E. coli* HST08 competent cells (Takara, 636763) and purified using a miniprep kit (Qiagen, 27104), in accordance with the manufacturer’s protocol.

#### APEX2 labeling in live cells

A375 cells were transfected with 8 μg of APEX2 fusion plasmid DNA using Lipofectamine 2000 (Invitrogen, 11668027) according to the manufacturer’s protocol. The biotinylation reaction of the APEX2-expressing constructs were done by replacing the medium with fresh high-glucose DMEM containing 800 μM biotin-phenol (Iris Biotech, ls-3500.1000). The cells were incubated at 37°C with 5% CO_2_ for 30 minutes. Next, H_2_O_2_ (Sigma-Aldrich, H1009) was added to the cells at a final concentration of 1 mM and incubated for one minute at room temperature. The reaction media was quenched by washing the cells three times with 1×PBS containing 5 mM Trolox (Sigma-Aldrich, 238813), 10 mM sodium ascorbate (Sigma-Aldrich, A4034-100G), and 10 mM sodium azide (Sigma-Aldrich, S2002).

#### Biotin-DBCO-N50-sC18* labeling in live cells

Biotin-DBCO-N50-sC18* was synthesized and purified as described in the Supplementary Methods.

For metabolic azide-labeling of sialylated glycoconjugates, A375 cells were grown in the presence of 50 µM Ac_4_ManNAz (C33366, Thermo Fisher) for 72 hours. For microscopy, cells were grown in 8-well Lab-Tek II Chamber Slides (154534PK, Thermo Fisher), while for streptavidin pull-down and mass spectrometry, cells were grown in 90 mm culture dishes (20101, SPL Life Sciences). Culture medium was replaced with Opti-MEM (31985070, Thermo Fisher) and the cells were incubated at 37°C with 5% CO_2_ for two hours to wash away and deplete residual Ac_4_ManNAz. The cells were then incubated with 5 µM Cy5.5 DBCO (CCT-1046, Vector Laboratories) in Opti-MEM for two hours to block cell surface azide groups, before 5 µM Biotin-DBCO-N50-sC18* in Opti-MEM was added for another two hours. After labeling, the cells were incubated in normal culture medium until assayed. Streptavidin pull-down and mass spectrometry were performed on samples harvested one hour after labeling. The samples were analyzed as described below (*Streptavidin affinity purification*).

Microscopy was performed at 0.5 hours, 24 hours, and 48 hours after labeling. To limit background staining, cytoplasmic depletion was performed before fixation and staining. The adherent cells were washed once with 1×PBS and then treated with freshly prepared cytoplasmic depletion buffer (10 mM PIPES pH 6.8, 300 mM sucrose, 100 mM NaCl, 3 mM MgCl_2_, and 0.7% (v/v) Triton X-100) for four minutes at RT. The cells were washed twice with 1×PBS and fixed with 4% paraformaldehyde in 1×PBS for 15 minutes at RT. The cells were washed once with permeabilization/quench buffer (PQB; 50 mM NH_4_Cl and 0.1% (v/v) Triton X-100 in 1×PBS) before 15 minutes incubation in fresh PQB. Then, the cells were incubated with blocking buffer (0.2% (w/v) gelatin in 1×PBS) for >5 minutes. The cells were stained with 2 µg/mL streptavidin-fluorescein in blocking buffer for one hour in a humidified chamber. The cells were then washed three times with 1×PBS and three times with ddH_2_O before being mounted using VECTASHIELD Vibrance Antifade Mounting Medium with DAPI (H-1800, Vector Laboratories). Images were acquired using a LSM980 microscope with Airyscan 2 (Zeiss), using the Airyscan 2 super-resolution mode and a 63x oil immersion objective. The images were processed using the default settings for Airyscan Processing in ZEN Blue and analyzed using QuPath 0.5.1. Fluorescence intensity was measured inside nuclei that were manually segmented based on the DAPI signal.

#### RENBP PCR

mRNA from A375 cells was isolated using the Total RNA Purification Plus Kit (48300, Norgen Biotek) according to the manufacturer’s instructions. cDNA was synthesized using the GoScript Reverse Transcription System (A5000, Promega) and oligo(dT) primers. The RENBP and GAPDH genes were amplified using PCR Master Mix (M7502, Promega) and the following cycling conditions: 96°C for 3.0 min; 30 cycles of 96°C for 45 s, 55°C or 57°C (GAPDH and RENBP, respectively) for 45 s, 72°C for 90 s; and then 72°C for 5.0 min. For amplification of RENBP, the PCR reaction buffer was supplemented with 6% (v/v) DMSO.

#### *Streptavidin* affinity purification

Cell pellets were resuspended in RIPA buffer (150 mM NaCl, 5 mM EDTA (pH 8.0), 50 mM Tris, 0.1% (v/v) NP-40, 0.5% sodium deoxycholate, 0.1% SDS), supplemented with protease inhibitors. The lysate was incubated on ice for 30 minutes and then sonicated once at 60% amplitude for 15 seconds (Sonics, VCX130). The lysates were centrifuged at 21,300×*g* for 15 minutes at 4°C. Prewashed Streptavidin Magnetic Beads (Thermo Fisher, 88816) were added to the protein lysates and incubated at room temperature for two hours. The streptavidin beads were subsequently washed three times with RIPA buffer. A fraction of the beads was processed for Western blot analyses by boiling in 2×Laemmli SDS reducing reagent supplemented with 2 mM free biotin. The remaining beads were resuspended in 50 mM triethylammonium bicarbonate (TEAB) buffer and further processed for LC-MS/MS analysis at the Proteomics Core Facility, University of Gothenburg.

#### Click chemistry

Ultra-pure nuclear lysates from A375 cells were mildly alkylated with 15 mM iodoacetamide (A39271, Thermo Fisher) for 30 min in the dark with gentle mixing. The lysates were then reacted with 50 µM PC DBCO Biotin (CCT-1120, Vector Laboratories) for three hours at RT with gentle mixing. Chloroform-methanol precipitation was performed by adding 600 µL methanol, 150 µL chloroform, and 400 µL ddH_2_O to 200 µL lysate, with mixing by vortexing in between. After centrifugation for five minutes at 18,000×*g*, the upper aqueous phase was carefully removed. The protein precipitate was then washed twice in methanol and finally resuspended in bead binding buffer (50 mM Tris-HCl pH 7.4, 0.6% (v/v) SDS). The sample was clarified by centrifugation for five minutes at 18,000×*g*. The supernatant was passed through a 0.1 µm Ultrafree Centrifugal Filer (UFC30VV, Sigma-Aldrich) before it was added to pre-washed Dynabeads MyOne Streptavidin C1 beads (65001, Thermo Fisher) and incubated overnight at 4°C. The beads were then washed twice in wash buffer 1 (0.1% (w/v) deoxycholic acid, 1% (v/v) Triton X-100, 1 mM EDTA, 500 mM NaCl, 50 mM HEPES pH 7.5), twice in wash buffer 2 (0.5% (w/v) deoxycholic acid, 0.05% (v/v) NP-40, 1 mM EDTA, 250 mM LiCl, 10 mM Tris-Cl pH 7.4), and finally four times in 50 mM Tris-HCl pH 7.4 before further processing for proteomic analysis at the Proteomics Core Facility, University of Gothenburg.

#### RBP immunoprecipitation

Lysates (either from ultra-pure nuclei or cytoplasmic fractions) were incubated overnight at 4°C with 4–7 µg of anti-RBP antibody, followed by a two hour incubation with prewashed Dynabeads Protein G (10003D, Thermo Fisher) at 4°C. Beads were washed three times with NP-40 lysis buffer (20 mM Tris-HCl, pH 7.5; 150 mM NaCl; 2 mM EDTA; 1% NP-40), and bound proteins were eluted with 2% acetic acid and neutralized with 1 M NaOH. Eluates were subjected to Western blot or glycomics analysis (Proteomics Core Facility, University of Gothenburg). Antibodies and lectins are listed in Table S22.

#### Enriched *O*-Glycan Immunoprecipitation

Ultra-pure nuclear lysates from A375 cells were treated with of 0.05 U α2-3/6/8-Sialidase (10269611001, Sigma-Aldrich) / 100 µg of total protein. Reactions were performed in 20 mM sodium acetate buffer (pH 5.5; AM9740, Merck) at 37°C for 60 minutes with shaking at 300 rpm. Samples were subsequently incubated overnight at 4°C with 4–6 µg of biotin-conjugated PNA lectin on a rotator. In parallel, pre-washed Dynabeads Protein G (10003D, Thermo Fisher) were suspended in NP-40 lysis buffer (20 mM Tris-HCl, pH 7.5; 150 mM NaCl; 2 mM EDTA; 1% NP-40 (v/v)) and incubated overnight at 4°C with 3–6 µg of anti-biotin antibody. The antibody-coupled beads were washed once with NP-40 buffer and combined with the lectin-incubated lysate. Samples rotated for an additional four hours at 4°C to capture biotinylated lectin–protein complexes. Following incubation, the beads were washed three times with NP-40 lysis buffer; each wash involved five minutes of gentle rotation at room temperature. Proteins bound to the beads were eluted with 2% acetic acid (A6283, Merck). Eluates were collected using a magnetic stand, immediately transferred to fresh tubes, and neutralized to pH 7.8 with 1 M NaOH. A fraction of the neutralized eluate was boiled for five minutes at 95°C with 1xLaemmli buffer (J60660.AC, Thermo Fisher) and analyzed by Western blot to confirm enrichment of *O*-glycosylated proteins. Remaining elutes were either sent for proteomic analysis at Proteomics Core Facility, University of Gothenburg or processed further for RNA-binding protein (RBP) immunoprecipitation.

#### Reverse enriched *O*-Glycan RBP immunoprecipitation

In a complementary approach, sialidase-treated nuclear lysates were first subjected to immunoprecipitation with 5 µg of anti-RBP antibody, followed by incubation with Dynabeads Protein G as described above. After elution and neutralization, samples were incubated overnight at 4°C with 4-5µg of biotin-conjugated PNA lectin. Lectin-bound complexes were captured using anti-biotin antibody-coated Dynabeads G protein (10003D, Thermo Fisher). Final eluates were processed for Western blot analysis or proteomic analysis at the Proteomics Core Facility, University of Gothenburg. Antibodies and lectins are listed in Table S22.

#### Glycomics

Glycomics analysis started from lysate (either from ultra-pure nuclei or cytoplasmic fractions). Samples were dot-blotted onto a PVDF membrane. The dots were excised and incubated with 20 µL of 0.5 M NaBH_4_ and 50 mM NaOH at 50°C overnight, to yield reduced glycans, released by reductive beta-elimination. After desalting, the dried samples were dissolved in 10 µL of water.

Released glycans were then analyzed by LC-MS/MS in negative ion mode on a LTQ linear ion trap mass spectrometer (Thermo Electron, San José, CA), with an IonMax standard ESI source equipped with a stainless-steel needle kept at –3.5 kV. The oligosaccharides (3 µL per sample), dissolved in water, were separated on a column (10 cm × 250 µm) packed in-house with 5 µm porous graphitized carbon particles (Hypercarb, Thermo-Hypersil, Runcorn, UK). The oligosaccharides were injected onto the column and eluted with an acetonitrile gradient (Buffer A, 10 mM ammonium bicarbonate; Buffer B, 10 mM ammonium bicarbonate in 80% acetonitrile). The gradient (0-45% Buffer B) was eluted for 46 min, followed by a wash step with 100% Buffer B, and equilibrated with Buffer A in the next 24 min.

Compressed air was used as nebulizer gas. The heated capillary was kept at 270°C, and the capillary voltage was –50 kV. A full scan (*m/z* 380-2000, two microscans, maximum 100 ms, target value of 30,000) was performed, followed by data-dependent MS^2^ scans (two microscans, maximum 100 ms, target value of 10,000) with normalized collision energy of 35%, isolation window of 2.5 units, activation q=0.25, and activation time 30 ms). The threshold for MS^2^ was set to 300 counts. Data acquisition and processing were conducted with Xcalibur software (Version 2.0.7). For comparing glycan abundances between samples, individual glycan structures were quantified relative to the total content by integrating the extracted ion chromatogram peak area. The area under the curve (AUC) of each structure was normalized to the total AUC and expressed as a percentage. The peak area was processed by Progenesis QI (Nonlinear Dynamics Ltd., Newcastle upon Tyne, UK).

#### Proteomics

Details for the experiment are provided in Supporting Information. Relative quantification was performed to compare protein abundances in different conditions. Beads with attached proteins were washed twice with one mL 50 mM triethylammonium bicarbonate (TEAB), then dissolved in 50 µL 50 mM TEAB, reduced and alkylated. Samples were digested by LysC/Trypsin, peptides labelled with TMTpro 18-plex isobaric mass tagging reagents (Thermo Fisher Scientific) and pooled into one TMT-set. The set was purified by HiPPR Detergent Removal Resin and fractionated with High-pH Spin Column into 10 fractions (all Pierce, Thermo Fisher Scientific). Each fraction was analyzed on an Orbitrap Tribrid mass spectrometer equipped with the FAIMS Pro ion mobility system interfaced with nLC 1200 liquid chromatography system (all Thermo Fisher Scientific). Peptides were separated on a C18 35 cm column over 90 minutes and data were acquired with the SPS MS3 method. Raw files were processed and analyzed with Proteome Discoverer against UniProt Swiss-Prot *Homo Sapiens* or a nuclear database with selected proteins together with a contaminant database using Sequest as a search engine.

#### Neu5Az-enriched proteomics

Beads with bound proteins, from Ac₄ManNAz and DMSO treated cells, were washed (4×1000 µL 50 mM TEAB) and incubated with 0.5 µg Trypsin/Lys-C Mix (A40009, Thermo Fisher) in 150 µL 50 mM TEAB at 37°C for two hours. The supernatant, including a 25 µL TEAB wash, was collected to identify captured proteins (Supernatant; S). S was digested overnight with 0.2 µg Trypsin (90059, Thermo Fisher) at 37°C, reduced with 5.5 mM dithiothreitol (30 min, 37°C), alkylated with 11.3 mM iodoacetamide (30 min at room temperature), then treated with an additional 0.3 µg Trypsin (2 h, 37°C). Samples were desalted using Pierce spin columns (89851, Thermo Fisher), dried, and reconstituted in 20 µL of 2% (v/v) acetonitrile, 0.1% (v/v) TFA for the LC-MS/MS analysis.

Samples were analyzed on an Orbitrap Exploris 480 interfaced with Easy-nLC1200 (all Thermo Fisher). Peptides were trapped on an Acclaim Pepmap 100 C18 trap column (100 μm x 2 cm, particle size 5 μm, Thermo Fisher Scientific) and separated on an in-house packed analytical column (75 μm × 30 cm, 3 μm, Reprosil-Pur C18, Dr. Maisch) using a 78 min 5-45% ACN gradient in 0.2% formic acid at 300 nL/min. MS1 scans (*m/z* 380–1500) were acquired at 120K resolution, AGC target 225%, max injection time 25 ms. The most abundant precursors with charges 2–7 were isolated with a 1.4 *m/z* window and fragmented by higher-energy collision dissociation (HCD) at 30% NCE. Fragment spectra were acquired at 30K resolution, with AGC target set to 150% and maximum injection time of 80 ms. The intensity threshold was set to 1.0e4 and dynamic exclusion to 10 ppm for 30 s.

Data were analyzed in Proteome Discoverer 2.4 using Sequest HT and Minora feature detection. The data was searched against a custom database consisting of Swiss-Prot human proteome (April 2023) plus four streptavidin entries, with a precursor mass tolerance of 5 ppm and a fragment ion tolerance of 0.02 Da. Tryptic peptides with up to two missed cleavages were accepted. Methionine oxidation was set as variable modifications and cysteine carbamidomethylation as fixed modification. Target-decoy approach was used for Peptide Spectrum Match (PSM) validation. For precursor ion quantification, the Minora Feature Detector was used with min trace length of 5 points and max delta RT of isotope pattern multiplets of 0.2 min. Both spectral counting at the PSM level and precursor ion quantification were used for evaluating protein differences between enriched glycoproteins from Ac₄ManNAz and DMSO treated cells.

#### CRISPR-Cas9 knockout screen

Cas9 lentivirus was produced using Lenti-X 293T cells (Takara Bio, CA, USA), the SiC-V1-Scr expression plasmid (Addgene # 133042), and the 2^nd^ Generation Packaging Mix & Lentifectin Combo Pack (abm, BC, Canada) according to the manufacturer’s protocol. Next, lentivirus particles in the supernatant were concentrated using the Lenti-X Concentrator (Takara Bio, CA, USA) following the manufacturer’s guidelines. The pellet was resuspended in 200 μL of DMEM media (Thermo Fisher), and aliquots were stored at -80°C. The viral titer was measured by transducing A549 cells (ATCC) with different volumes of Cas9 lentivirus and performing flow cytometry two days post-transduction. The same protocol was applied to the GlycoGene CRISPR library lentivirus preparation, except that the Human GlycoGene CRISPR pooled library (Addgene #140961) was used as the expression plasmid.

Isogenic A549 Cas9 cells were generated following Kelkar et al.^54^, with some modifications. Briefly, SiC-V1-Scr lentivirus was added at a multiplicity of infection (MOI) of 2.7 to 2 x 10^5^ A549 cells in nutrient mix media (Thermo Fisher) supplemented with 10 μg/mL polybrene (Sigma), in a total reaction volume of 100 μL. The cells were incubated for 10 min and then transferred to a well of a 12-well plate containing 600 μL of media, supplemented with 10 μg/mL polybrene. Then the cells were incubated at 37°C for two hours, and 300 μL of media with 10 μg/mL polybrene was added. The media was changed after 24 h post-transduction, and the cells were expanded over the following days. On day 12, fluorescence activated cell sorting (FACS) was performed to bulk sort the A549 Cas9 cells (Cas9^+^). On day 20, the bulk-sorted cells were single-cell sorted into wells of a 96-well plate to obtain isogenic A549 Cas9 cells.

A549 library cells were produced similar to the above-mentioned transduction protocol with minor modifications. Briefly, 4 x 10^6^ isogenic A549 Cas9 cells were transduced with GlycoGene CRISPR pooled lentivirus at an MOI of 0.36 in a 150 mm tissue culture plate containing 30 mL of nutrient mix media (Thermo Fisher) supplemented with 10 μg/mL polybrene (Sigma). To initiate gene editing, 1 μg/mL doxycycline (Dox) (abm, BC, Canada) was added two days post-transduction. After four days post-transduction, Dox was removed.

Library-transduced cells were expanded and processed as described in the *flow cytometry* section. The cells were expanded until day 11, when FACS was performed to bulk-sort A549 Library cells (BFP+ Cas9+).

#### Nuclei

Briefly, isolated nuclei were fixed and stained with MAL-II-biotin at 10 µg/mL and subsequently streptavidin-fluorescein at 2 µg/mL, each for 20 minutes at RT. The nuclei were sorted on a FACSAria III Cell Sorter (BD) for top and bottom 20% of the fluorescein signal. The sorted nuclei were de-crosslinked by incubation in PureLink Genomic Lysis/Binding Buffer (K182302, Thermo Fisher) supplemented with Proteinase K and RNase overnight at 56°C with gentle mixing. Genomic DNA was isolated using PureLink Genomic DNA Mini Kit (K1820-01, Thermo Fisher) and subsequently prepared for sequencing.

#### Cells

Harvested A549 glycogene CRISPR library cells were washed with cold staining buffer (1x TBS; 100 mM NaCl, 1 mM CaCl_2_, 1 mM MgCl_2_, 20 mM Tris base pH 7.2, 1% (w/v) BSA), and 5 x 10^6^ cells were stained on ice for 15 minutes with biotinylated MAL-II (20 µg/mL in cold staining buffer, B-1265-1, Vector). Cells were then washed twice and incubated with streptavidin–fluorescein (2 µg/mL in cold staining buffer, SA-5001-1, Vector) for 15 minutes on ice. After two additional washes, cell pellets were passed through a cell strainer (352235, Corning) and resuspended in FACS buffer (HBSS (Gibco), 1% (v/v) FBS, 1 mM EDTA) prior to sorting on a FACSAria III Cell Sorter. The top and bottom 20% of fluorescein-positive cells were collected. Genomic DNA was extracted from both low- and high-binding populations after each sort using the PureLink Genomic DNA Mini Kit (K1820-01, Thermo Fisher).

The regions flanking the sgRNA were PCR amplified using 1 μg of gDNA as the template, 1 μM of NGSlib Fwd primer, 1 μM of NGSlib Rev primer, and 25 μL of NEBNext^®^ Ultra™ II Q5® Master Mix (New England Biolabs, MA, USA) in a 50 μL reaction volume. The resulting PCR product was run on a 1.2% (w/v) agarose gel electrophoresis, and the band corresponding to the expected molecular weight (307 bp) was gel purified using the NucleoSpin^®^ Gel and PCR Clean-up kit (Macherey-Nagel). Subsequently, the sequencing library was prepared with the gel-purified amplicons and the Native Barcoding Kit 24 V14 (Oxford Nanopore), following the manufacturer’s protocol. Sequencing was performed on a MinION platform with real-time basecalling, using a FLO-MIN114 (R10.4.1) flow cell, for a run time of 72 h and a Min Q score of 8 (nuclei) or 9 (cells). The NGS data were analyzed as explained in Kelkar et al.^54^, and Zhu et al.^40^, to determine the count output and quantify sgRNA enrichment and depletion using MAGeCK^55^.

#### COG siRNA knock-down

225 pmol control or COG4/6/7-targeting esiRNA were complexed with 22.5 µL Lipofectamine 2000 (11668019, Thermo Fisher) in a total volume of 1500 µL Opti-MEM (31985062, Thermo Fisher) for 15 min before being added to a subconfluent 10-cm dish containing HEK293T cells in 7.5 mL growth medium. After 48 hours, an aliquot of cells was collected for validation of knock-down efficiency via real-time qPCR, and the remaining cells processed for lectin flow cytometry. mRNA was isolated using the Total RNA Purification Plus Kit (48300, Norgen Biotek) according to the manufacturer’s instructions and cDNA was synthesized using the GoScript Reverse Transcription System (A5000, Promega) and oligo(dT) primers. Real-time qPCR was performed using iTaq Universal SYBR Green Supermix (1725121, Bio-Rad) and the CFX Connect Real-Time System (Bio-Rad). Gene expression was quantified using the 2-ddCt method with GAPDH as the reference. Oligo sequences and references are listed in Table S22.

#### Immunofluorescence staining

For fluorescence microscopy, cells were seeded onto 18 mm glass coverslips in 12-well plates. Cells were fixed with 4% (v/v) paraformaldehyde (Solveco, 1267) for 15 minutes at room temperature. After fixation, cells were washed three times with 1×PBS and permeabilized with 0.1% (v/v) Triton X-100 (Merck, 2341264) in 1×PBS for 10 minutes at room temperature. Cells were then blocked in 5% bovine serum albumin (BSA; Thermo Fisher, 11423164) in PBS for 30 minutes. Cells were incubated with primary antibodies overnight in a humidified chamber at 4°C. The following day, cells were washed three times with PBS and incubated with appropriate secondary fluorescent antibodies for one hour at room temperature in a humidified chamber. Cells were washed three times with 1×PBS and stained with 0.3 nM DAPI (Fisher Scientific, 15205739) for five minutes at room temperature. Finally, cells were washed three times with 1×PBS and mounted onto coverslips using ProLong Diamond Antifade reagent (Fisher Scientific, P36961).

To visualize the endoplasmic reticulum, A375 cell lysates were spread on poly-L-lysine–coated coverslips, air-dried, and immunostained with anti-Calnexin antibody, followed by fluorescent secondary antibody and DAPI counterstaining, as described above. Antibodies are listed in Table S22.

#### SUnSET assay

A375 cells were transfected with 3 µg FLAG-tagged RPP30^WT^ or RPP30^S55A^ using Lipofectamine 2000 (11668027; Invitrogen), according to the manufacturer’s instructions. 48 hours post-transfection, to measure global protein translation rates, cells were treated with water or puromycin (5 µg/mL; P9620; Sigma-Aldrich) for 15 min at 37°C, after which medium was aspirated and cells were washed once with ice-cold PBS. Cell numbers were determined at collection, and equivalent cell number (1.0x10^5^ cells) were used across all conditions. Lysates and Western blots were prepared as described above. Puromycin incorporation into nascent polypeptide chains was detected by immunoblotting with an anti-puromycin antibody (MABE343; Sigma Aldrich), serving as a proxy for translation rate. Signal intensity across the entire puromycin-positive smear was quantified using ImageJ and compared between conditions.

#### tRNA qRT-PCR

RNA immunoprecipitation was performed to isolate RNA associated with FLAG-tagged RPP30^WT^ or RPP30^S55A^ proteins. Lysate from overexpressed FLAG-tagged RPP30^WT^ or RPP30^S55A^ A375 cells were prepared in RIPA buffer (150 mM NaCl, 5 mM EDTA (pH 8.0), 50 mM Tris, 0.1% (v/v) NP-40, 0.5% sodium deoxycholate, 0.1% SDS), supplemented with protease inhibitors, as described above. Clarified lysates were incubated with prewashed anti-FLAG M2 magnetic beads at 4°C with rotation overnight.

Following incubation, beads were washed three times with RIPA buffer. RNA was extracted using TRIzol reagent (15596026; Thermo Fisher Scientific) according to the manufacturer’s instructions, RNA pellets were resuspended in RNase-free water. RNA was quantified using Qubit RNA HS assay (Q32852; Invitrogen).

The abundance of 55 mature tRNAs was quantified using a qPCR method originally described by Ou et al.^56^. Briefly, 5 µL of eluted RNA was mixed with 1 µL of 100 mM Tris-HCl pH 9.0 and incubated at 37°C for 40 min to deacetylate mature tRNAs. 1 µL of 10×TE pH 7.5 was added for pH adjustment, before annealing with 20 pmol U-adaptor in 2 µL for 3 min at 90°C, followed by addition of 1 µL 50 mM Tris-HCl pH 8.0 and further incubation at 37°C for 20 min. Ligation was performed using 1 U T4 RNA Ligase 2 (M0239S, New England Biolabs) with the total volume adjusted to 20 µL and incubation at 37°C for 1 hour. U-adaptor-ligated tRNAs were then annealed with a mixture of specific tRNA reverse primers (0.44 µM final concentration; RT mix 1: primers 1-30; RT mix 2: primers 31-55) by incubation at 65°C for 5 min. cDNA was then synthesized using PrimeScript RT Master Mix (RR036A, Takara) according to the manufacturer’s instructions. qRT-PCR was performed using iTaq-Universal SYBR Green Supermix (1725121, Bio-Rad) in 10 µL reactions containing 500 nM forward and reverse primers and 2 µL cDNA (diluted 11-fold). Each sample was analyzed in technical duplicates using the CFX Connect Real-Time System (Bio-Rad) to monitor amplification. Thermal cycling conditions included initial denaturation at 95°C for 10 min followed by 50 cycles of 10 s at 95°C, 30 s at 58°C, and 40 s at 72°C and a final melt curve analysis.

#### Data curation

We searched the academic literature for studies performed with (i) *O*-glycoproteomics, (ii) on whole cells or tissues, and (iii) not using enrichment methods specific for *O*-GlcNAc. This was followed by a filtering script to filter out any hits containing the following substrings in any of their GO terms pertaining to subcellular localization, as retrieved from UniProt: [“Cell membrane”, “Secreted”, “Golgi”, “ndoplasmic”, “vesicle”, “Lysosome”, “Single-pass”, “Multi-pass”]. Hits without any subcellular localization GO terms were also removed. Then, a manual pass followed, in which each remaining hit was rigorously investigated as to whether its GO terms indicate an association with the secretory pathway. This procedure finally resulted in 5,288 glycopeptide reports from noncanonical subcellular compartments, comprising 1,322 unique proteins, with 2,299 unique glycosylation sites, from 31 peer-reviewed publications (Table S9).

To specifically search for putative nuclear proteins in our data, we assembled a custom database from the “Nucleoplasm” and “Nucleoli” segments in the Human Protein Atlas^8^, resulting in 7,406 unique human proteins (Table S18) that was used in proteomics experiments in this work. Any findings resulting from this database were also reproduced in full searches against all human proteins.

#### Quantification and statistical analysis

All glycan-related analyses were conducted using glycowork (version 1.5.0). Comparing two groups was done via two-tailed Welch’s t-tests. Comparing more than two groups was done via one-way ANOVA with Tukey’s multiple comparison test. Testing for enrichment via contingency tables was performed via Fisher’s exact test. In all cases, significance was defined as p < 0.05. All multiple testing was corrected with the Benjamini-Hochberg procedure. All statistical testing has been done in Python 3.11.3 using the statsmodels package (version 0.14.0) and the scipy package (version 1.11.0rc1). Effect sizes were calculated as Cohen’s *d* using glycowork (version 1.5.0).

## Acknowledgments

HEK293T*^SLC35A1-/-^* KO cells were a kind gift from Dr. Ritva Tikkanen (University of Giessen, Germany). HEK293T*^SLC35A2-/-^*KO cells were a kind gift from Dr. Mariusz Olczak (University of Wrocław, Poland). This work was funded by a Branco Weiss Fellowship – Society in Science (to D.B.), by the Knut and Alice Wallenberg Foundation (to D.B. and PAR 2020/228 to A.A.S.), the IngaBritt and Arne Lundberg Research Foundation (to D.B.), the Jeansson Foundations (to D.B.), the Swedish Society for Medical Research (to A.A.S.), the Swedish Research Council (grant no. 2019-01855 and 2025-02255 to A.A.S. and 2022-03825 to D.B.), the University of Gothenburg, Sweden (to A.A.S. and D.B.), the European Research Council (to D.B., SWEETSWAP, grant no. 101219123), and the Canada Excellence Research Chairs Program (CERC in Glycomics, L.K.M.). Proteomic analysis was performed at the Proteomics Core Facility, Sahlgrenska academy, Gothenburg University, with financial support from SciLifeLab and BioMS. We acknowledge the Center for Cellular Imaging at the University of Gothenburg and the National Microscopy Infrastructure, NMI (VR-RFI 2019–00217) for providing assistance in microscopy. We thank Dr. Iwona Nowak for technical support generating the APEX2-Golgi constructs. We thank C.K. Wong for his help with MAGeCK.

## Author contributions

A.A.S., D.B., and J.L., conceptualization; A.A.S., D.B., E.S., J.L., M.F., U.B., and V.L. formal analysis; C.J. glycomics; A.T., E.M., J.F. proteomics; A.A.T. and J.C.J.H. chemical peptide synthesis; F.I.M., J.L., and L.K.M. CRISPR/Cas9 knockout screen; D.B. data curation; D.B. and J.L. writing–original draft. A.A.S., A.A.T., A.T., D.B., E.M., E.S., F.I.M., J.F., J.C.J.H., J.L., L.K.M., M.F., U.B., and V.L. writing–review & editing; A.A.S., D.B., J.L., M.F., and U.B. visualization; A.A.S., A.A.T., D.B., and L.K.M. supervision; A.A.S., D.B., and L.K.M. funding acquisition; A.A.S., D.B., J.L., M.F., and V.L. methodology; A.A.S., J.L., M.F., and U.B. validation.

## References

1. Varki, A. (2017). Biological roles of glycans. Glycobiology 27, 3–49. 10.1093/glycob/cww086.

2. National Research Council (US) Committee on Assessing the Importance and Impact of Glycomics and Glycosciences (2012). Transforming Glycoscience: A Roadmap for the Future (National Academies Press) 10.17226/13446.

3. Yang, X., and Qian, K. (2017). Protein O-GlcNAcylation: emerging mechanisms and functions. Nat Rev Mol Cell Biol 18, 452–465. 10.1038/nrm.2017.22.

4. West, C.M., Slawson, C., Zachara, N.E., and Hart, G.W. (2022). Nucleocytoplasmic Glycosylation. In Essentials of Glycobiology, A. Varki, R. D. Cummings, J. D. Esko, P. Stanley, G. W. Hart, M. Aebi, D. Mohnen, T. Kinoshita, N. H. Packer, J. H. Prestegard, et al., eds. (Cold Spring Harbor Laboratory Press).

5. Haudek, K.C., Spronk, K.J., Voss, P.G., Patterson, R.J., Wang, J.L., and Arnoys, E.J. (2010). Dynamics of galectin-3 in the nucleus and cytoplasm. Biochimica et Biophysica Acta (BBA) - General Subjects 1800, 181–189. 10.1016/j.bbagen.2009.07.005.

6. Wang, J., Wu, G., Miyagi, T., Lu, Z.-H., and Ledeen, R.W. (2009). Sialidase occurs in both membranes of the nuclear envelope and hydrolyzes endogenous GD1a. Journal of Neurochemistry 111, 547–554. 10.1111/j.1471-4159.2009.06339.x.

7. Münster, A.-K., Weinhold, B., Gotza, B., Mühlenhoff, M., Frosch, M., and Gerardy-Schahn, R. (2002). Nuclear Localization Signal of Murine CMP-Neu5Ac Synthetase Includes Residues Required for Both Nuclear Targeting and Enzymatic Activity. Journal of Biological Chemistry 277, 19688–19696. 10.1074/jbc.M201093200.

8. Uhlén, M., Fagerberg, L., Hallström, B.M., Lindskog, C., Oksvold, P., Mardinoglu, A., Sivertsson, Å., Kampf, C., Sjöstedt, E., Asplund, A., et al. (2015). Tissue-based map of the human proteome. Science 347, 1260419. 10.1126/science.1260419.

9. Meyer, C.F., Seath, C.P., Knutson, S.D., Lu, W., Rabinowitz, J.D., and MacMillan, D.W.C. (2022). Photoproximity Labeling of Sialylated Glycoproteins (GlycoMap) Reveals Sialylation-Dependent Regulation of Ion Transport. J. Am. Chem. Soc. 144, 23633–23641. 10.1021/jacs.2c11094.

10. Wiktor, M., Wiertelak, W., Maszczak-Seneczko, D., Balwierz, P.J., Szulc, B., and Olczak, M. (2021). Identification of novel potential interaction partners of UDP-galactose (SLC35A2), UDP-N-acetylglucosamine (SLC35A3) and an orphan (SLC35A4) nucleotide sugar transporters. Journal of Proteomics 249, 104321. 10.1016/j.jprot.2021.104321.

11. Cejas, R.B., Lorenz, V., Garay, Y.C., and Irazoqui, F.J. (2019). Biosynthesis of O-N-acetylgalactosamine glycans in the human cell nucleus. Journal of Biological Chemistry 294, 2997–3011. 10.1074/jbc.RA118.005524.

12. Cejas, R.B., Garay, Y.C., de la Fuente, S., Lardone, R.D., and Irazoqui, F.J. (2020). Core 1 O-N-acetylgalactosamine (O-GalNAc) glycosylation in the human cell nucleus. Biological Chemistry 401, 1041–1051. 10.1515/hsz-2019-0448.

13. Long, Y., Li, Z., Wang, L., Ao, X., Zhang, Z., Chen, Q., Zhu, D., Liu, X., Liu, R., Chen, B., et al. (2024). Highly efficient identification of nucleocytoplasmic O-glycosylation by the TurboID-based proximity labeling method in living cells. Biotechnology Journal 19, 2300090. 10.1002/biot.202300090.

14. Xu, Z., Ku, X., Tomioka, A., Xie, W., Liang, T., Zou, X., Cui, Y., Sato, T., Kaji, H., Narimatsu, H., et al. (2020). O-linked N-acetylgalactosamine modification is present on the tumor suppressor p53. Biochimica et Biophysica Acta (BBA) - General Subjects 1864, 129635. 10.1016/j.bbagen.2020.129635.

15. Ledeen, R., and Wu, G. (2011). New findings on nuclear gangliosides: overview on metabolism and function. Journal of Neurochemistry 116, 714–720. 10.1111/j.1471-4159.2010.07115.x.

16. Xu, Z., Gong, Q., Xia, B., Groves, B., Zimmermann, M., Mugler, C., Mu, D., Matsumoto, B., Seaman, M., and Ma, D. (2009). A role of histone H3 lysine 4 methyltransferase components in endosomal trafficking. Journal of Cell Biology 186, 343–353. 10.1083/jcb.200902146.

17. Citterio, C., Jones, H.D., Pacheco-Rodriguez, G., Islam, A., Moss, J., and Vaughan, M. (2006). Effect of protein kinase A on accumulation of brefeldin A-inhibited guanine nucleotide-exchange protein 1 (BIG1) in HepG2 cell nuclei. Proc. Natl. Acad. Sci. U.S.A. 103, 2683–2688. 10.1073/pnas.0510571103.

18. Kuroda, F., Moss, J., and Vaughan, M. (2007). Regulation of brefeldin A-inhibited guanine nucleotide-exchange protein 1 (BIG1) and BIG2 activity via PKA and protein phosphatase 1γ. Proc. Natl. Acad. Sci. U.S.A. 104, 3201–3206. 10.1073/pnas.0611696104.

19. Lobo, V., Nowak, I., Fernandez, C., Correa Muler, A.I., Westholm, J.O., Huang, H.-C., Fabrik, I., Huynh, H.T., Shcherbinina, E., Poyraz, M., et al. (2024). Loss of Lamin A leads to the nuclear translocation of AGO2 and compromised RNA interference. Nucleic Acids Research 52, 9917–9935. 10.1093/nar/gkae589.

20. Gagnon, K.T., Li, L., Janowski, B.A., and Corey, D.R. (2014). Analysis of nuclear RNA interference in human cells by subcellular fractionation and Argonaute loading. Nat Protoc 9, 2045–2060. 10.1038/nprot.2014.135.

21. Huynh, H.T., Shcherbinina, E., Huang, H., Rezaei, R., and Sarshad, A.A. (2024). Biochemical Separation of Cytoplasmic and Nuclear Fraction for Downstream Molecular Analysis. Current Protocols 4, e1042. 10.1002/cpz1.1042.

22. Lundstrøm, J., Urban, J., Thomès, L., and Bojar, D. (2023). GlycoDraw: a python implementation for generating high-quality glycan figures. Glycobiology, cwad063. 10.1093/glycob/cwad063.

23. Bojar, D., Meche, L., Meng, G., Eng, W., Smith, D.F., Cummings, R.D., and Mahal, L.K. (2022). A Useful Guide to Lectin Binding: Machine-Learning Directed Annotation of 57 Unique Lectin Specificities. ACS Chem. Biol., acschembio.1c00689. 10.1021/acschembio.1c00689.

24. Banning, A., Zakrzewicz, A., Chen, X., Gray, S.J., and Tikkanen, R. (2021). Knockout of the CMP–Sialic Acid Transporter SLC35A1 in Human Cell Lines Increases Transduction Efficiency of Adeno-Associated Virus 9: Implications for Gene Therapy Potency Assays. Cells 10, 1259. 10.3390/cells10051259.

25. Baskin, J.M., Prescher, J.A., Laughlin, S.T., Agard, N.J., Chang, P.V., Miller, I.A., Lo, A., Codelli, J.A., and Bertozzi, C.R. (2007). Copper-free click chemistry for dynamic *in vivo* imaging. Proc. Natl. Acad. Sci. U.S.A. 104, 16793–16797. 10.1073/pnas.0707090104.

26. Gronewold, A., Horn, M., and Neundorf, I. (2018). Design and biological characterization of novel cell-penetrating peptides preferentially targeting cell nuclei and subnuclear regions. Beilstein J. Org. Chem. 14, 1378–1388. 10.3762/bjoc.14.116.

27. Luchansky, S.J., Yarema, K.J., Takahashi, S., and Bertozzi, C.R. (2003). GlcNAc 2-Epimerase Can Serve a Catabolic Role in Sialic Acid Metabolism*. Journal of Biological Chemistry 278, 8035–8042. 10.1074/jbc.M212127200.

28. Lobo, V., Shcherbinina, E., Westholm, J.O., Nowak, I., Huang, H.-C., Angeletti, D., Anastasakis, D.G., and Sarshad, A.A. (2024). Integrative transcriptomic and proteomic profiling of the effects of cell confluency on gene expression. Sci Data 11, 617. 10.1038/s41597-024-03465-z.

29. Han, S.-S., Shim, H.-E., Park, S.-J., Kim, B.-C., Lee, D.-E., Chung, H.-M., Moon, S.-H., and Kang, S.-W. (2018). Safety and Optimization of Metabolic Labeling of Endothelial Progenitor Cells for Tracking. Sci Rep 8, 13212. 10.1038/s41598-018-31594-0.

30. Castello, A., Fischer, B., Eichelbaum, K., Horos, R., Beckmann, B.M., Strein, C., Davey, N.E., Humphreys, D.T., Preiss, T., Steinmetz, L.M., et al. (2012). Insights into RNA Biology from an Atlas of Mammalian mRNA-Binding Proteins. Cell 149, 1393–1406. 10.1016/j.cell.2012.04.031.

31. Steentoft, C., Vakhrushev, S.Y., Joshi, H.J., Kong, Y., Vester-Christensen, M.B., Schjoldager, K.T.-B.G., Lavrsen, K., Dabelsteen, S., Pedersen, N.B., Marcos-Silva, L., et al. (2013). Precision mapping of the human O-GalNAc glycoproteome through SimpleCell technology. EMBO J 32, 1478–1488. 10.1038/emboj.2013.79.

32. Thomès, L., Burkholz, R., and Bojar, D. (2021). Glycowork: A Python package for glycan data science and machine learning. Glycobiology, cwab067. 10.1093/glycob/cwab067.

33. Han, H., Shi, Q., Zhang, Y., Ding, M., He, X., Liu, C., Zhao, D., Wang, Y., Du, Y., Zhu, Y., et al. (2024). RBM12 drives PD-L1-mediated immune evasion in hepatocellular carcinoma by increasing JAK1 mRNA translation. Oncogene 43, 3062–3077. 10.1038/s41388-024-03140-y.

34. Zhu, Y., Groth, T., Kelkar, A., Zhou, Y., and Neelamegham, S. (2021). A GlycoGene CRISPR-Cas9 lentiviral library to study lectin binding and human glycan biosynthesis pathways. Glycobiology 31, 173–180. 10.1093/glycob/cwaa074.

35. Wiertelak, W., Pavlovskyi, A., Maszczak-Seneczko, D., Szulc, B., and Olczak, M. (2023). The glycosylation defect in solute carrier SLC35A2/SLC35A3 double knockout cells is rescued by SLC35A2–SLC35A3 and SLC35A3–SLC35A2 hybrids. FEBS Letters 597, 2345–2357. 10.1002/1873-3468.14714.

36. Zhang, N., Lin, S., Cui, W., and Newman, P.J. (2022). Overlapping and unique substrate specificities of ST3GAL1 and 2 during hematopoietic and megakaryocytic differentiation. Blood Advances 6, 3945–3955. 10.1182/bloodadvances.2022007001.

37. Blackburn, J.B., D’Souza, Z., and Lupashin, V.V. (2019). Maintaining order: COG complex controls Golgi trafficking, processing, and sorting. FEBS Letters 593, 2466–2487. 10.1002/1873-3468.13570.

38. Willett, R., Blackburn, J.B., Climer, L., Pokrovskaya, I., Kudlyk, T., Wang, W., and Lupashin, V. (2016). COG lobe B sub-complex engages v-SNARE GS15 and functions via regulated interaction with lobe A sub-complex. Sci Rep 6, 29139. 10.1038/srep29139.

39. Blackburn, J.B., D’Souza, Z., and Lupashin, V.V. (2019). Maintaining order: COG complex controls Golgi trafficking, processing, and sorting. FEBS Letters 593, 2466–2487. 10.1002/1873-3468.13570.

40. Zhu, Y., Groth, T., Kelkar, A., Zhou, Y., and Neelamegham, S. (2021). A GlycoGene CRISPR-Cas9 lentiviral library to study lectin binding and human glycan biosynthesis pathways. Glycobiology 31, 173–180. 10.1093/glycob/cwaa074.

41. Engineering glycosyltransferases into glycan binding proteins using a novel mammalian surface display platform (2025). 10.21203/rs.3.rs-5004188/v1.

42. Foulquier, F., Amyere, M., Jaeken, J., Zeevaert, R., Schollen, E., Race, V., Bammens, R., Morelle, W., Rosnoblet, C., Legrand, D., et al. (2012). TMEM165 Deficiency Causes a Congenital Disorder of Glycosylation. The American Journal of Human Genetics 91, 15–26. 10.1016/j.ajhg.2012.05.002.

43. Lam, S.S., Martell, J.D., Kamer, K.J., Deerinck, T.J., Ellisman, M.H., Mootha, V.K., and Ting, A.Y. (2015). Directed evolution of APEX2 for electron microscopy and proximity labeling. Nat Methods 12, 51–54. 10.1038/nmeth.3179.

44. Cao, Q., Wang, Z., Wan, H., Xu, L., You, X., Liao, L., and Chen, Y. (2018). PAQR3 Regulates Endoplasmic Reticulum-to-Golgi Trafficking of COPII Vesicle via Interaction with Sec13/Sec31 Coat Proteins. iScience 9, 382–398. 10.1016/j.isci.2018.11.002.

45. Lyu, Z., Sycks, M.M., Espinoza, M.F., Nguyen, K.K., Montoya, M.R., Galapate, C.M., Mei, L., and Genereux, J.C. (2022). Monitoring Protein Import into the Endoplasmic Reticulum in Living Cells with Proximity Labeling. ACS Chem. Biol. 17, 1963–1977. 10.1021/acschembio.2c00405.

46. Piecyk, M., Fauvre, J., Duret, C., Chaveroux, C., and Ferraro-Peyret, C. (2024). SUrface SEnsing of Translation (SUnSET), a Method Based on Western Blot Assessing Protein Synthesis Rates in vitro. BIO-PROTOCOL 14. 10.21769/BioProtoc.4933.

47. Chen, S., Qin, R., and Mahal, L.K. (2021). Sweet systems: technologies for glycomic analysis and their integration into systems biology. Critical Reviews in Biochemistry and Molecular Biology 56, 301–320. 10.1080/10409238.2021.1908953.

48. Bandini, G., Agop-Nersesian, C., Van Der Wel, H., Mandalasi, M., Kim, H.W., West, C.M., and Samuelson, J. (2021). The nucleocytosolic O-fucosyltransferase SPINDLY affects protein expression and virulence in Toxoplasma gondii. Journal of Biological Chemistry 296, 100039. 10.1074/jbc.RA120.015883.

49. Hofsteenge, J., Huwiler, K.G., Macek, B., Hess, D., Lawler, J., Mosher, D.F., and Peter-Katalinic, J. (2001). C-Mannosylation and O-Fucosylation of the Thrombospondin Type 1 Module. Journal of Biological Chemistry 276, 6485–6498. 10.1074/jbc.M008073200.

50. Flynn, R.A., Pedram, K., Malaker, S.A., Batista, P.J., Smith, B.A.H., Johnson, A.G., George, B.M., Majzoub, K., Villalta, P.W., Carette, J.E., et al. (2021). Small RNAs are modified with N-glycans and displayed on the surface of living cells. Cell 184, 3109–3124.e22. 10.1016/j.cell.2021.04.023.

51. Smith, E.M., Ly, J., Haug, S., and Cheeseman, I.M. (2026). RNase MRP subunit composition and role in 40S ribosome biogenesis. Nat Struct Mol Biol 33, 20–33. 10.1038/s41594-025-01690-7.

52. Pokrovskaya, I.D., Willett, R., Smith, R.D., Morelle, W., Kudlyk, T., and Lupashin, V.V. (2011). Conserved oligomeric Golgi complex specifically regulates the maintenance of Golgi glycosylation machinery. Glycobiology 21, 1554–1569. 10.1093/glycob/cwr028.

53. Perez-Riverol, Y., Bai, J., Bandla, C., García-Seisdedos, D., Hewapathirana, S., Kamatchinathan, S., Kundu, D.J., Prakash, A., Frericks-Zipper, A., Eisenacher, M., et al. (2022). The PRIDE database resources in 2022: a hub for mass spectrometry-based proteomics evidences. Nucleic Acids Res 50, D543–D552. 10.1093/nar/gkab1038.

54. Kelkar, A., Groth, T., and Neelamegham, S. (2022). Forward Genetic Screens of Human Glycosylation Pathways Using the GlycoGene CRISPR Library. Current Protocols 2, e402. 10.1002/cpz1.402.

55. Li, W., Xu, H., Xiao, T., Cong, L., Love, M.I., Zhang, F., Irizarry, R.A., Liu, J.S., Brown, M., and Liu, X.S. (2014). MAGeCK enables robust identification of essential genes from genome-scale CRISPR/Cas9 knockout screens. Genome Biology 15, 554. 10.1186/s13059-014-0554-4.

56. Ou, X., Ma, B., Zhang, R., Miao, Z., Cheng, A., Peppelenbosch, M.P., and Pan, Q. (2020). A simplified qPCR method revealing tRNAome remodeling upon infection by genotype 3 hepatitis E virus. FEBS Letters 594, 2005–2015. 10.1002/1873-3468.13764.

