## Supplemental Figures for "Extended nuclear glycosylation regulates RNA processing"

#### Supplementary Figures

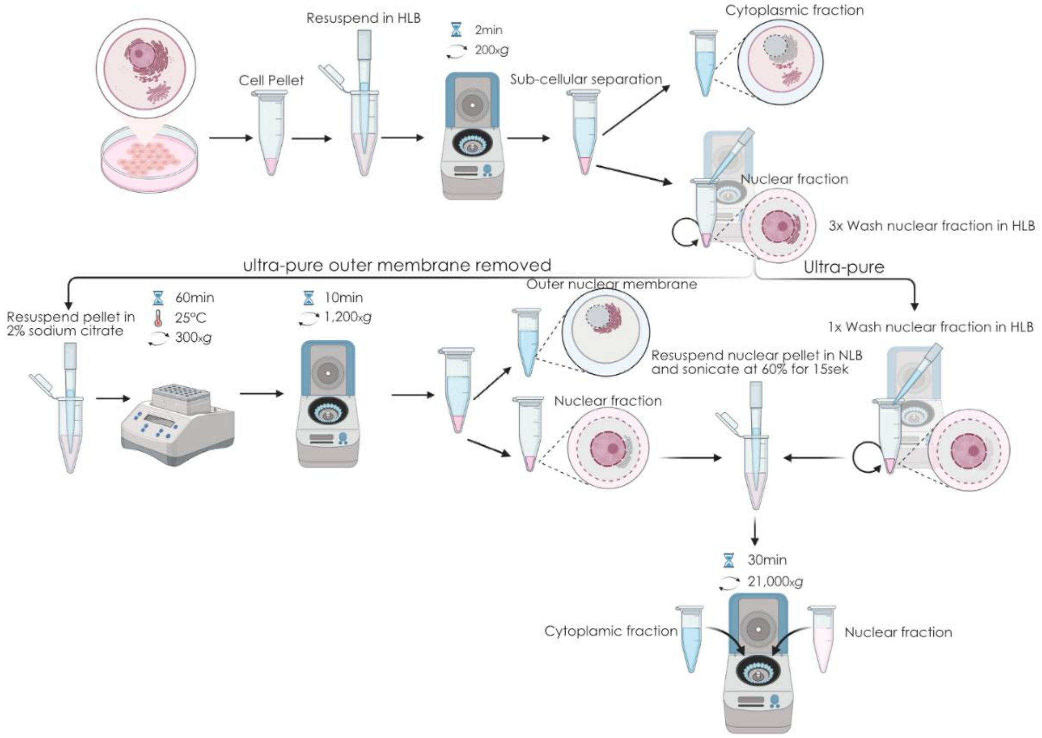

**Figure S1**. **Preparation of ultra-pure nuclear lysates.** Schematic showing the experimental workflow for the isolation of ultra-pure nuclear lysates, including the steps for removal of the outer nuclear membrane (Methods; Isolation of highly pure nuclei). The figure was prepared using BioRender.

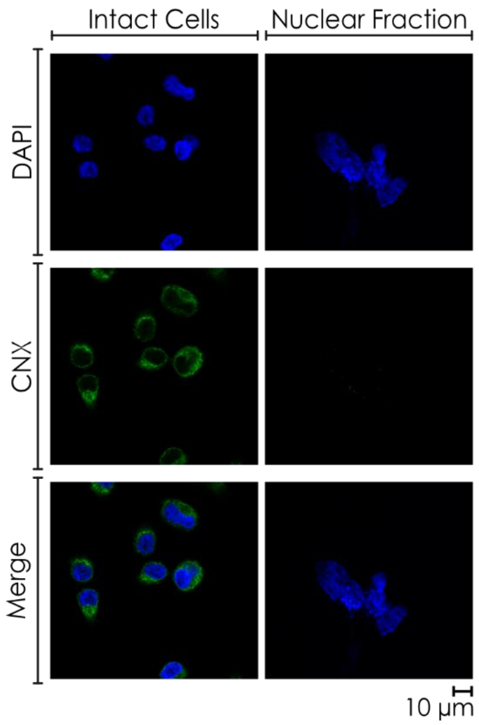

**Figure S2. Ultra-pure nuclear lysates are free of calnexin contamination.** Fluorescence microscopy showing detection of calnexin (CNX, green) in intact cells, but not around ultra-pure nuclei (DAPI, blue; Methods; Isolation of highly pure nuclei). Scale bar = 10 µm.

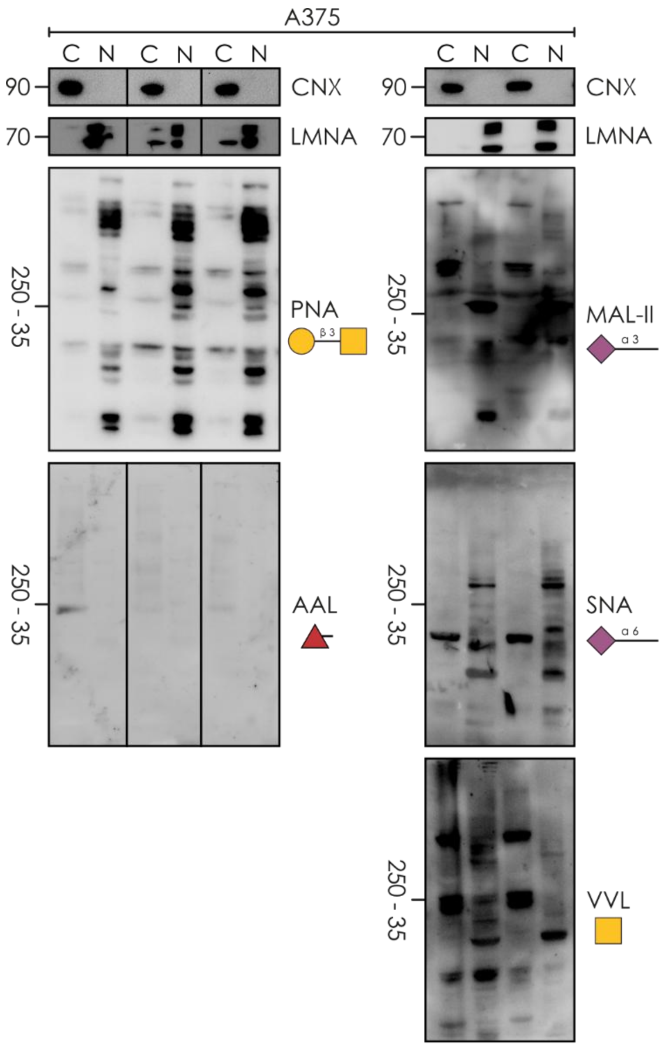

**Figure S3. Lectin blot detection of extended glycosylation in ultra-pure nuclear lysates of A375 cells.** Ultra-pure nuclei of A375 cells can be stained with several lectins that bind to extended *O*-GalNAc type glycans, including PNA, AAL, MAL-II, SNA, and VVL. Markers include Calnexin/CNX (ER) and Lamin A/LMNA (nucleus). C: cytoplasm, N: nucleus.

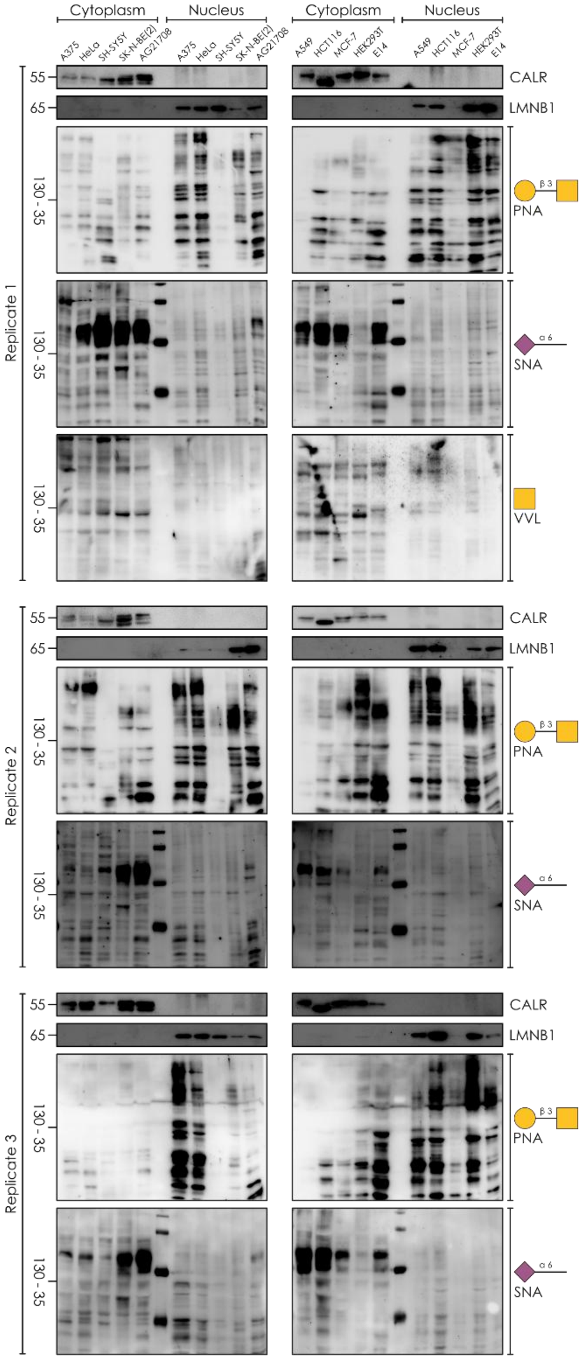

**Figure S4. Lectin blot detection of extended glycosylation in purified nuclear lysates of additional mammalian cell lines.** The tested cell lines include A375 (melanoma, human), HeLa (cervical carcinoma, human), SH-SY5Y (neuroblastoma, human), SK-N-BE (neuroblastoma, human), AG21708 (primary dermal fibroblasts, human), A549 (lung carcinoma, human), HCT116 (colorectal carcinoma, human), MCF-7 (breast adenocarcinoma, human), HEK293T (immortalized kidney embryonic cells, human), and E14 (embryonic stem cells, mouse). n = 3 (PNA and SNA), n = 1 (VVL). Markers include Calreticulin/CALR (ER) and Lamin B/LMNB1 (nucleus). We note that labile sialic acid is partially lost during Western blot sample preparation, yielding higher PNA staining and lower SNA/MAL-II staining than in our other analysis methods (i.e., glycomics/flow cytometry).

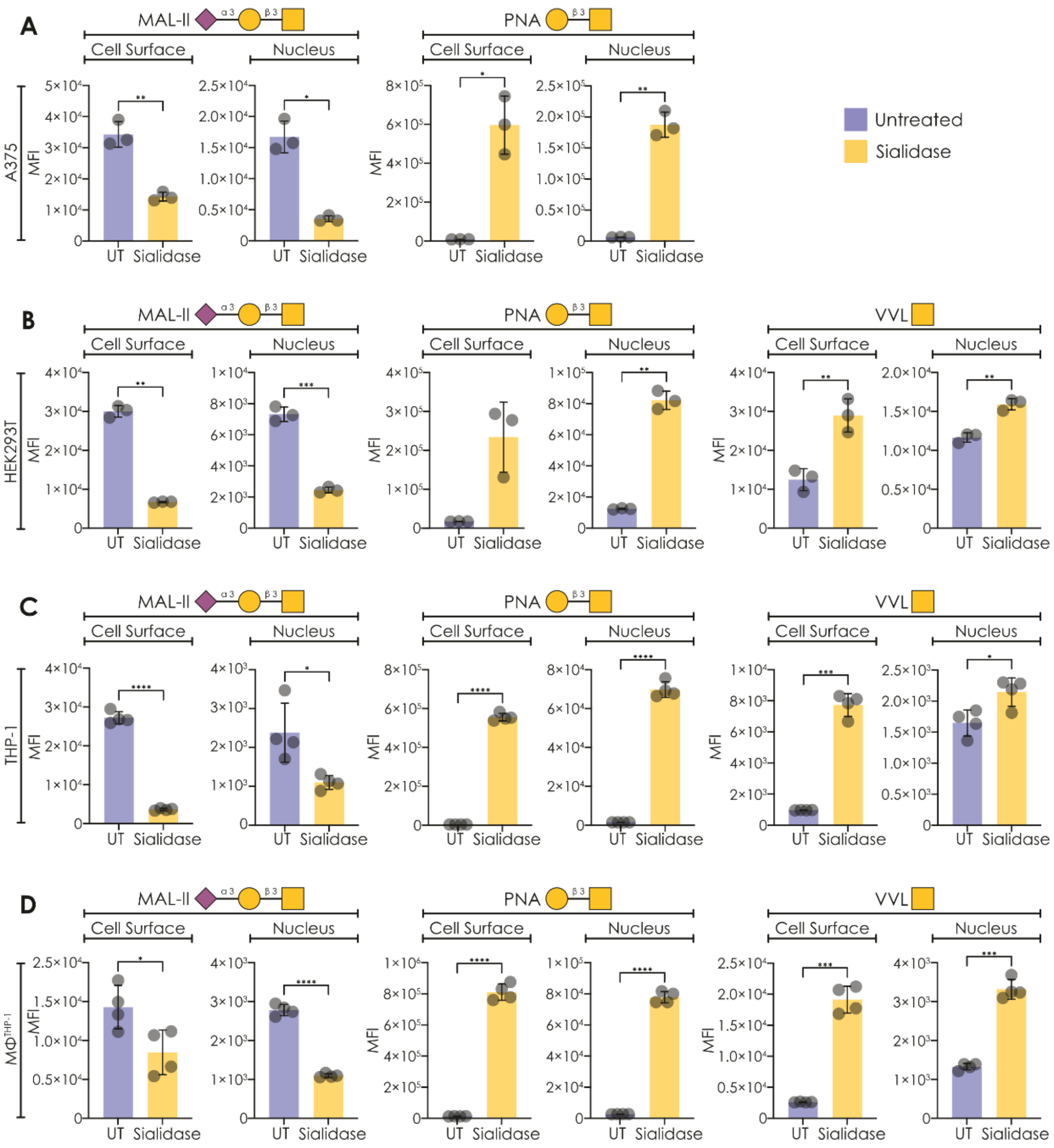

**Figure S5. Measuring extended *O*-glycosylation in the nuclei of diverse cell lines. A-D)** Glycosylation features on cell surfaces and pure nuclei of A375 (n = 3; A), HEK293T (n = 3; B), THP-1 (n = 4; C), and PMA-differentiated THP-1; $M\Phi^{THP\text{-}1}$ (n = 4; D) were quantified via flow cytometry using the lectins MAL-II (Neu5Acα2-3 on *O*-linked glycans), PNA (core 1 *O*-glycans), and VVL (terminal GalNAc), both with and without the application of sialidase. Significant differences were established via a Welch's t-test. *p < 0.05, **p < 0.01, ***p < 0.001, ****p < 0.0001.

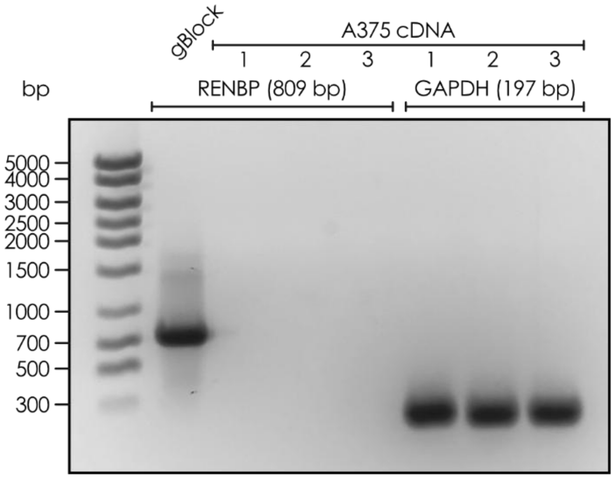

**Figure S6. RENBP is not expressed in A375 cells.** cDNA from A375 cells was analyzed for expression of RENBP and GAPDH with PCR and visualized on an agarose gel. A synthetic gBlock gene fragment was used as a positive control for verification of the RENBP primers (n = 3).

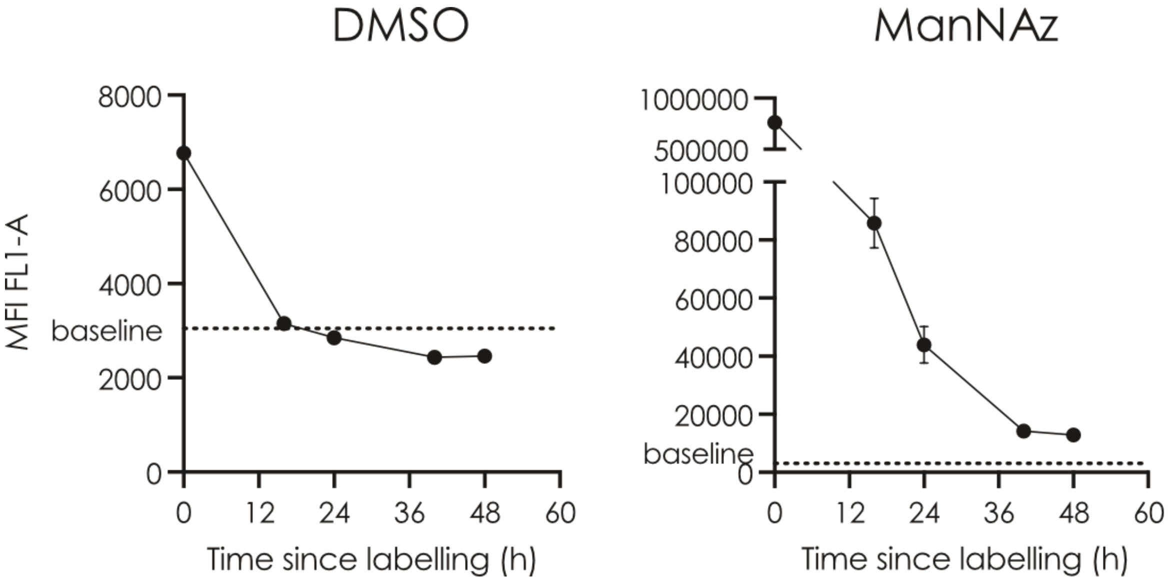

**Figure S7. Click reaction with glycoproteins increases peptide half-life in cells.** Cells grown with 50 µM ManNAz or DMSO (as control) for 72 h were labeled with 5 µM of Biotin-DBCO-N50-sC18* (n = 3) for two hours before being washed away. Then, for several timepoints during 48 hours after the labeling, FITC fluorescence (indicating peptide presence) was monitored via flow cytometry and compared between the two conditions. While DMSO-grown cells washed out any peptide already in the first timepoint, ManNAz-grown cells still exhibited detectable peptide after 48 hours, indicating successful click chemistry reactions.

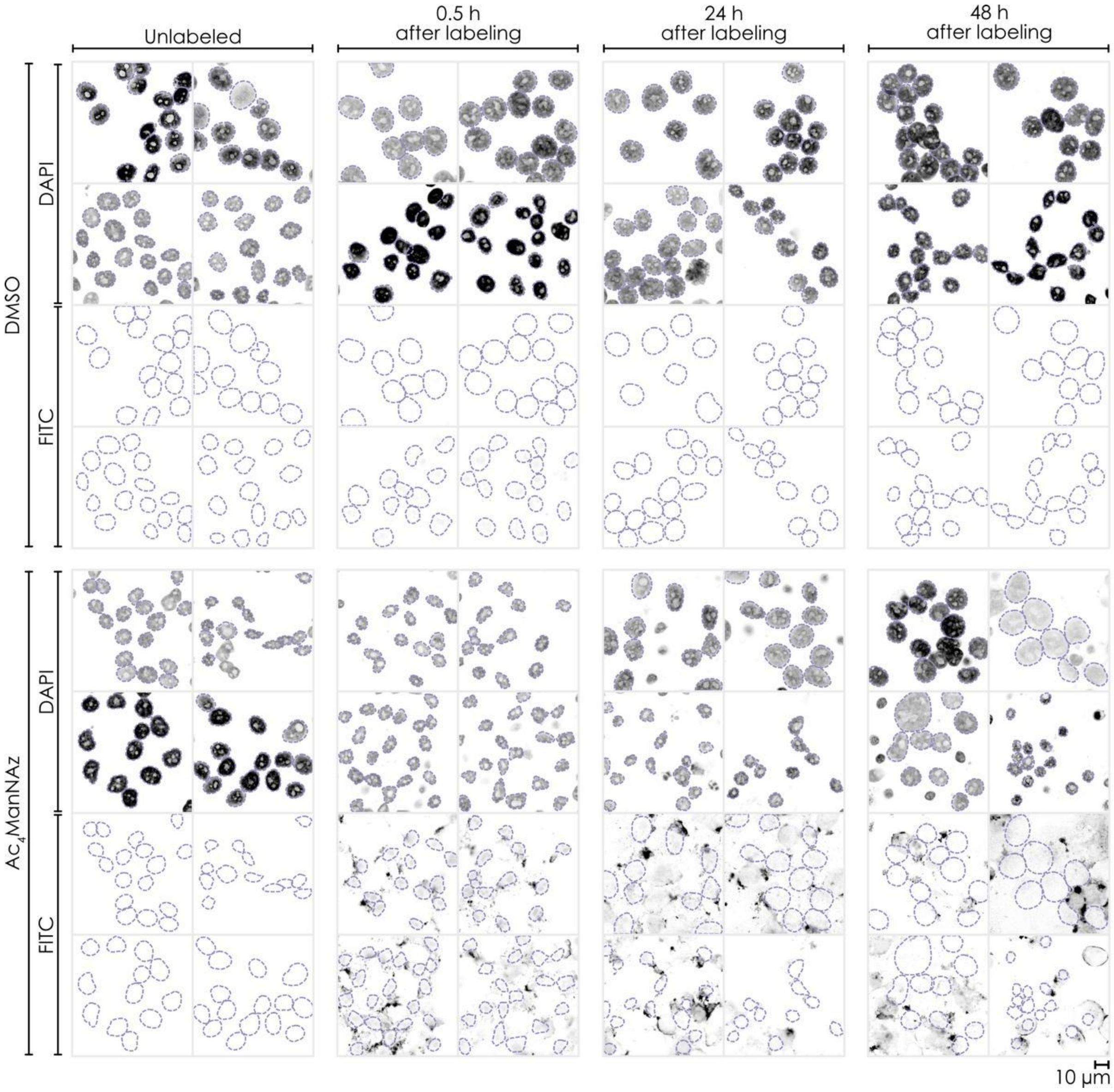

**Figure S8. Confocal imaging of cell-penetrating, nucleus-homing peptide in intact cells**. Images of DMSO-grown (control) cells and ManNAz-grown (click chemistry) cells were taken at timepoints 0.5 h, 24 h, and 48 h after labeling with Biotin-DBCO-N50-sC18*. Unlabeled cells were included as a control. Nuclei were segmented based on DAPI staining. The biotin-tagged peptide was imaged via staining with streptavidin-fluorescein. Prolonged staining of nuclei only in ManNAz-grown cells suggested a successful click chemistry reaction between the DBCO-conjugated peptide and Neu5Az-containing nuclear glycoproteins. Scale bar: 10 µm.

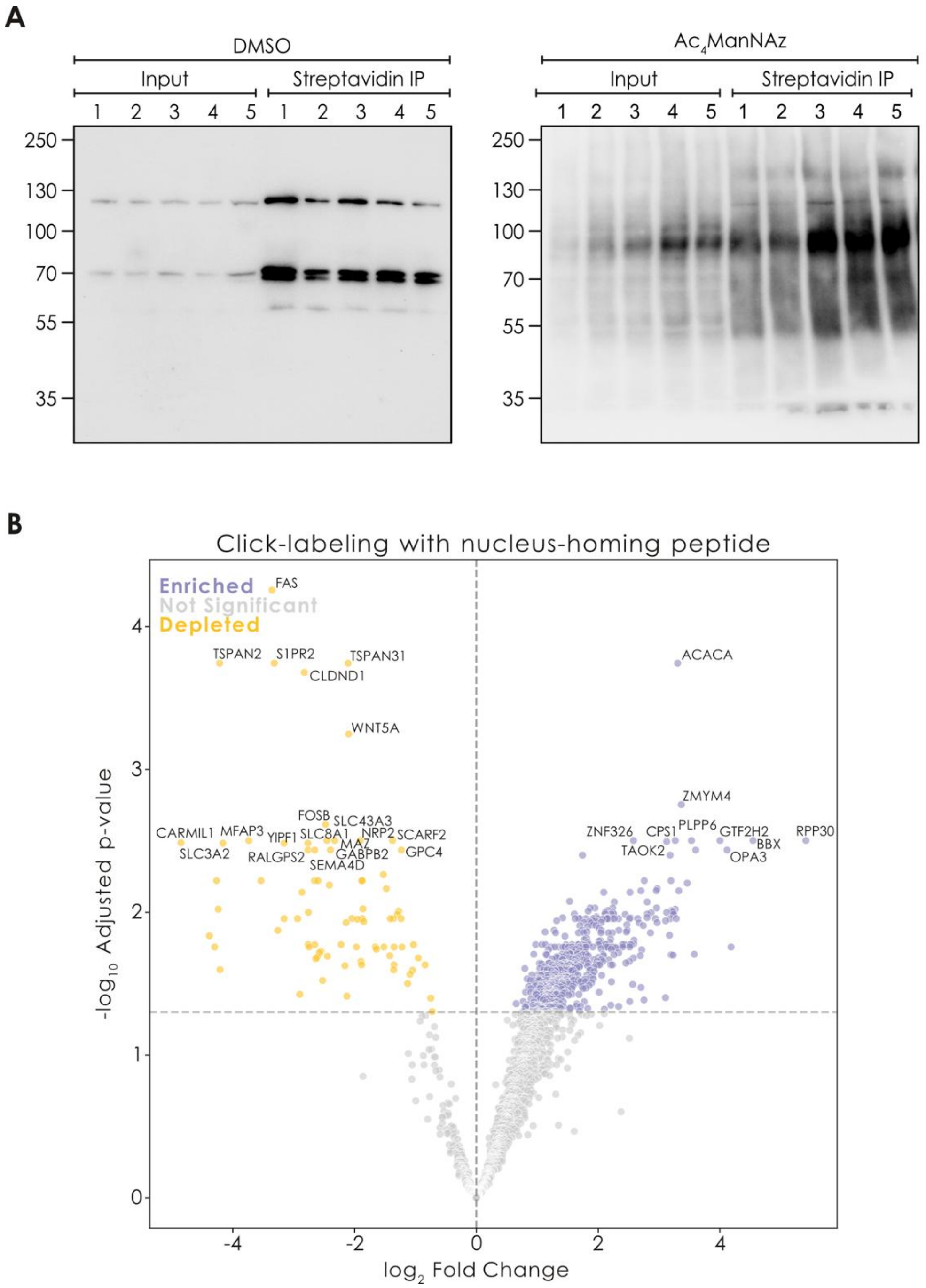

**Figure S9. Nucleus-homing peptide click-reacts with sialylated nuclear glycoproteins. A)** Western blot of peptide-labeled A375 cells grown with either DMSO or 50 µM ManNAz, both as input and immunoprecipitation with streptavidin from five replicates. Since our designer peptide was biotinylated, streptavidin capture indicated a successful click reaction. Bands in the DMSO-control stemmed from naturally biotinylated proteins that are outcompeted in the ManNAz condition. **B)** Volcano plot of proteomics data of streptavidin IP between DMSO and ManNAz grown cells. Significance was established via Welch's t-test and Benjamini-Hochberg correction for multiple testing. A horizontal line indicates the significance cut-off of $p_{adj} = 0.05$. Enriched proteins were assumed to be reacting with the click-peptide, since no click-reaction could occur in the DMSO condition.

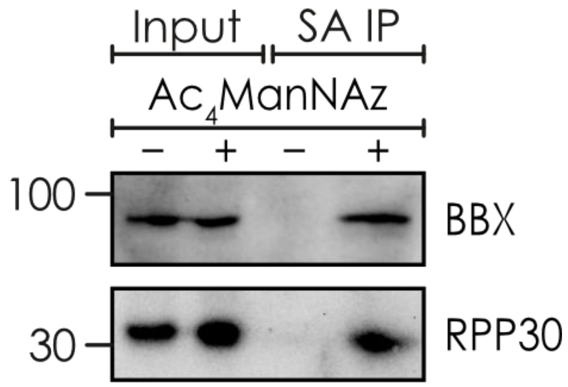

**Figure S10. Nuclear RPP30 and BBX carry sialylated, extended *O*-glycans.** Western blots of streptavidin-IPs for the nuclear proteins BBX and RPP30 after labeling of A375 cells grown in the presence or absence of $Ac_4ManNAz$ with Biotin-DBCO-N50-sC18*. SA IP: streptavidin immunoprecipitation.

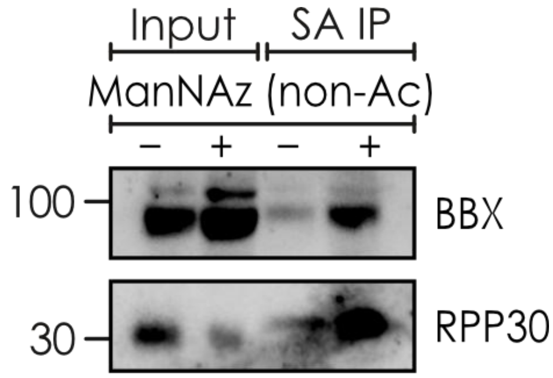

**Figure S11. Click chemistry results are not due to non-enzymatic *S*-glycosylation.** Western blots of streptavidin-IPs for the nuclear proteins BBX and RPP30 after labeling of A375 cells grown in the presence or absence of non-acetylated ManNAz with Biotin-DBCO-N50-sC18*. SA IP: streptavidin immunoprecipitation.

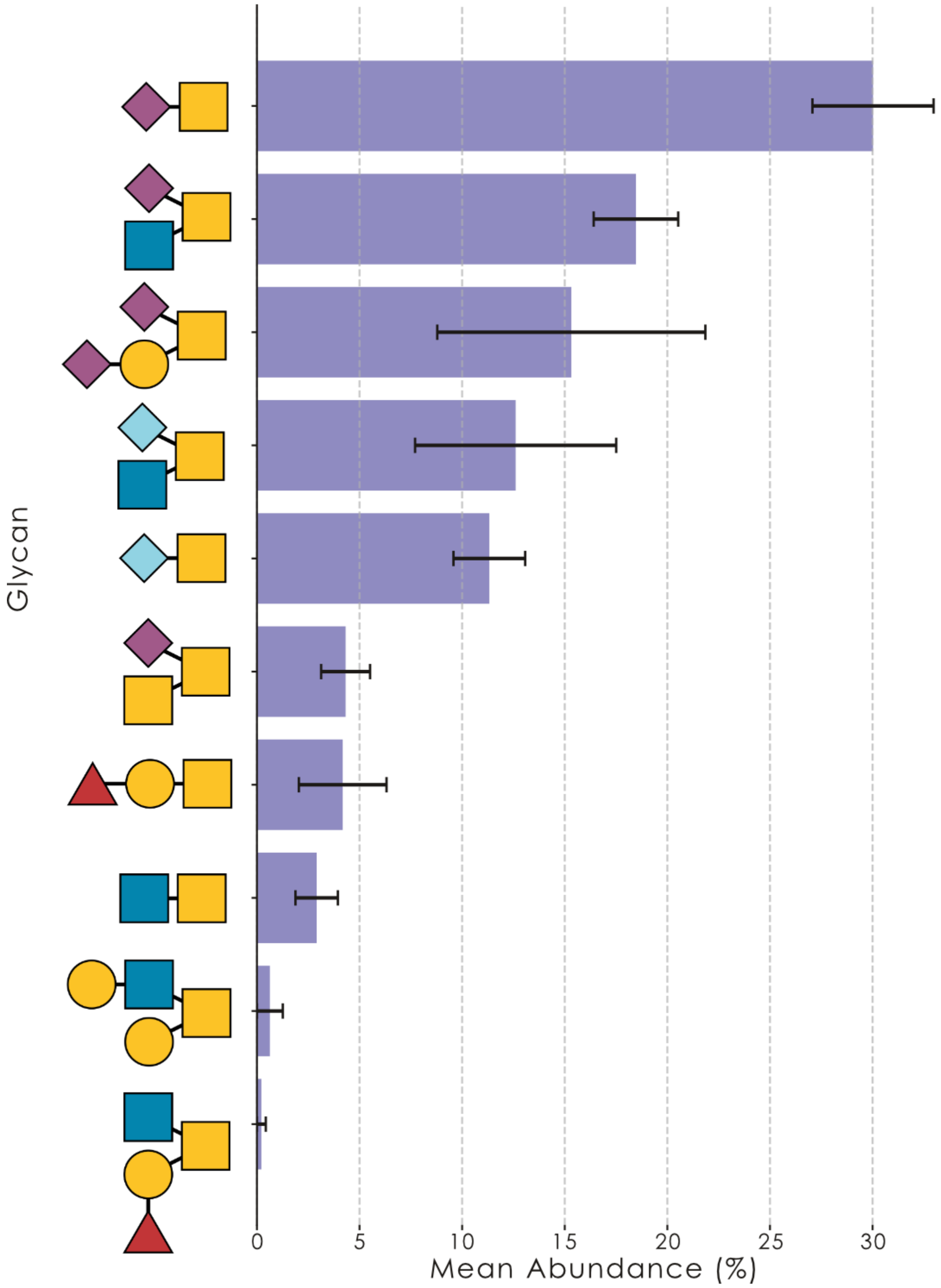

**Figure S12. Glycomics of ultra-pure HEK293T nuclei.** *O*-glycans in purified HEK293T nuclei (n = 9) were released via reductive elimination and measured via mass spectrometry. The results are shown as relative abundances in bar graphs that include the standard deviation, as well as the respective glycans in the SNFG depiction.

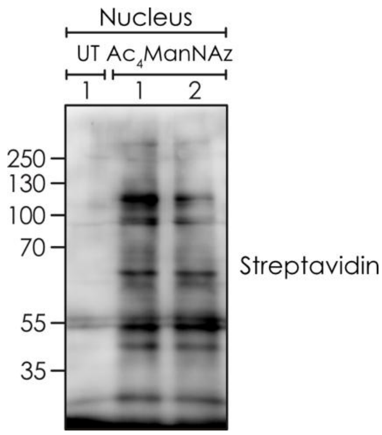

**Figure S13. ManNAz/Neu5Az is incorporated into nuclear glycoproteins.** We extracted ultra-pure nuclei from A375 cells grown with DMSO (UT) or added ManNAz, followed by click chemistry-driven biotinylation. Shown is a Western blot, in which this biotinylation was detected via streptavidin in two replicates, where only nuclei from ManNAz-grown cells exhibited distinct protein bands, presumably stemming from sialylated nuclear glycoproteins.

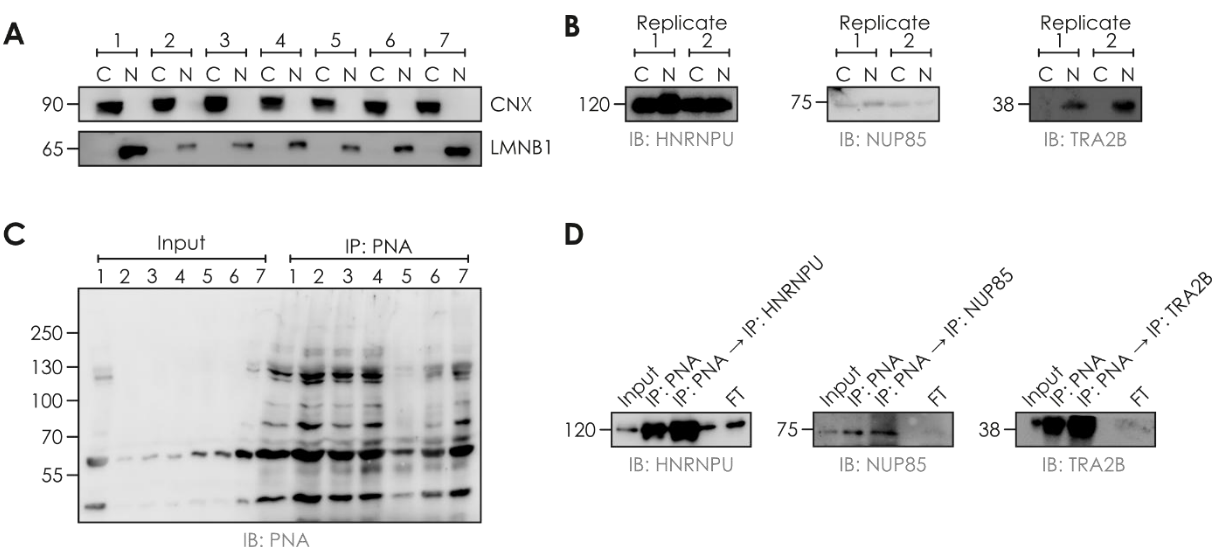

**Figure S14. Tandem purifications of nuclear glycoproteins. A-B)** We fractionated A375 cells into cytoplasm (C) and ultra-pure nuclei (N), using Calnexin (CNX) as a marker for the ER and Lamin B1 (LMNB1) as a marker for the nucleus (A). Then, we assessed the cytoplasmic vs nuclear localization of three nuclear proteins identified in our PNA capture enrichment (Fig. 2F), HNRNPU, NUP85, and TRA2B (B). **C-D)** Starting with an immunoprecipitation of sialidase-treated, ultra-pure nuclei with the lectin PNA (C), we pooled the elution of fractions 1-4 and 7 and performed a second immunoprecipitation against HNRNPU, NUP85, or TRA2B, respectively (D).

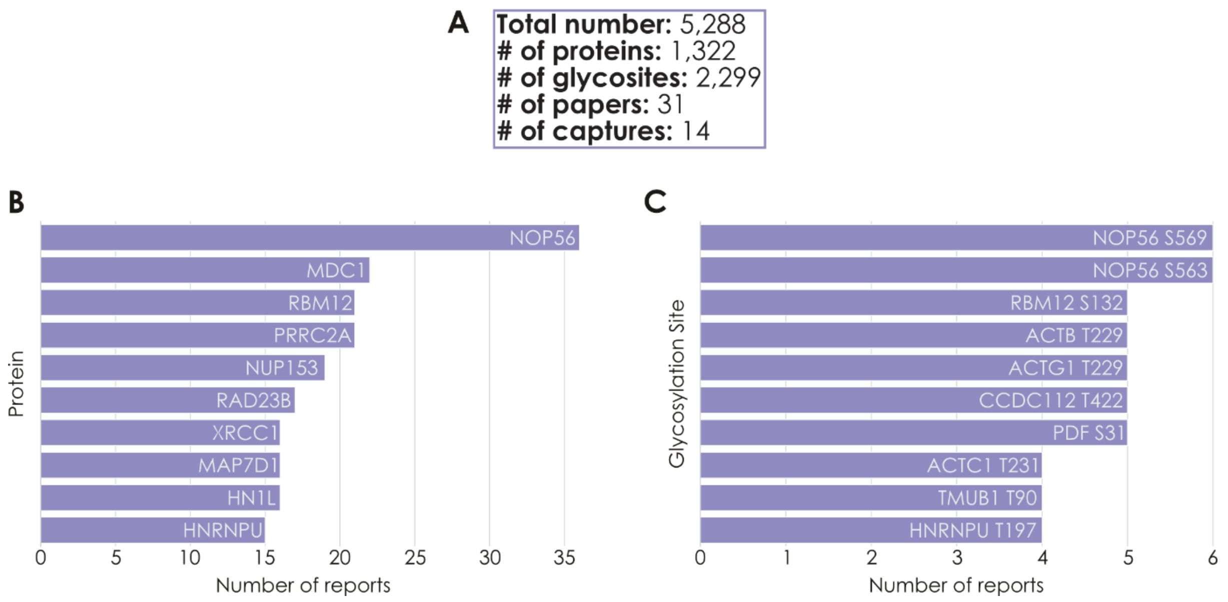

**Figure S15. Conserved nuclear glycoproteins are inadvertently reported in literature whole-cell/-tissue *O*-glycoproteomics data. A)** Overview of noncanonical literature *O*-glycoproteomics data (see Table S9 for full list). **B)** Ten representative noncanonical *O*-glycoproteins and how often they have been reported in the literature (including several glycopeptides from the same study). **C)** Ten representative noncanonical *O*-glycosylation sites and how often they have been reported in the literature.

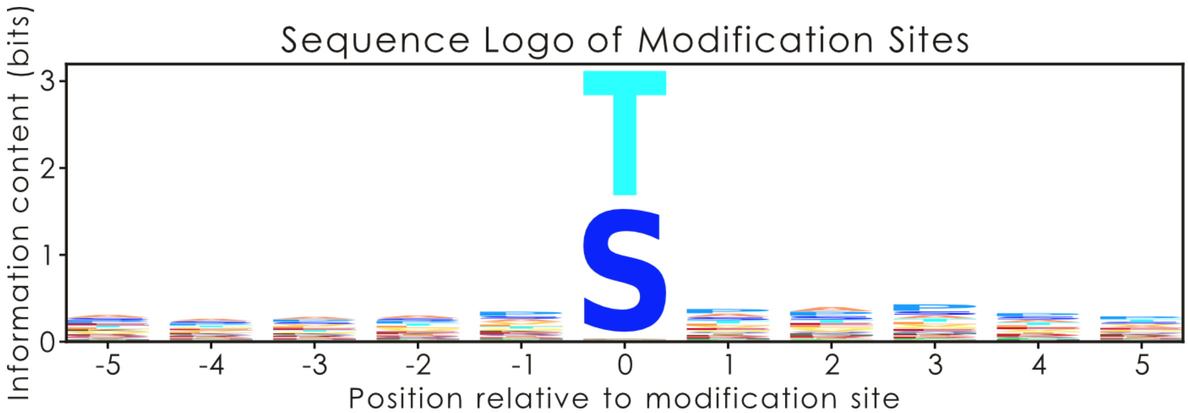

**Figure S16. Logo plot of amino acid sequence around noncanonical *O*-glycosylation sites.** Using all unique human glycosylation sites from our literature dataset (Table S9), we obtained their sequon window (+/- five amino acids around the glycosylation site) from their UniProt sequence. Shown is the logo plot for all 11 sequon positions, with individual amino acids color-coded and scaled by their information content (maximum possible information with 20 amino acids: 4.32). Overall, we observe enrichment for proline and small amino acids, such as alanine. Logo plots were visualized via Logomaker (version 0.8).

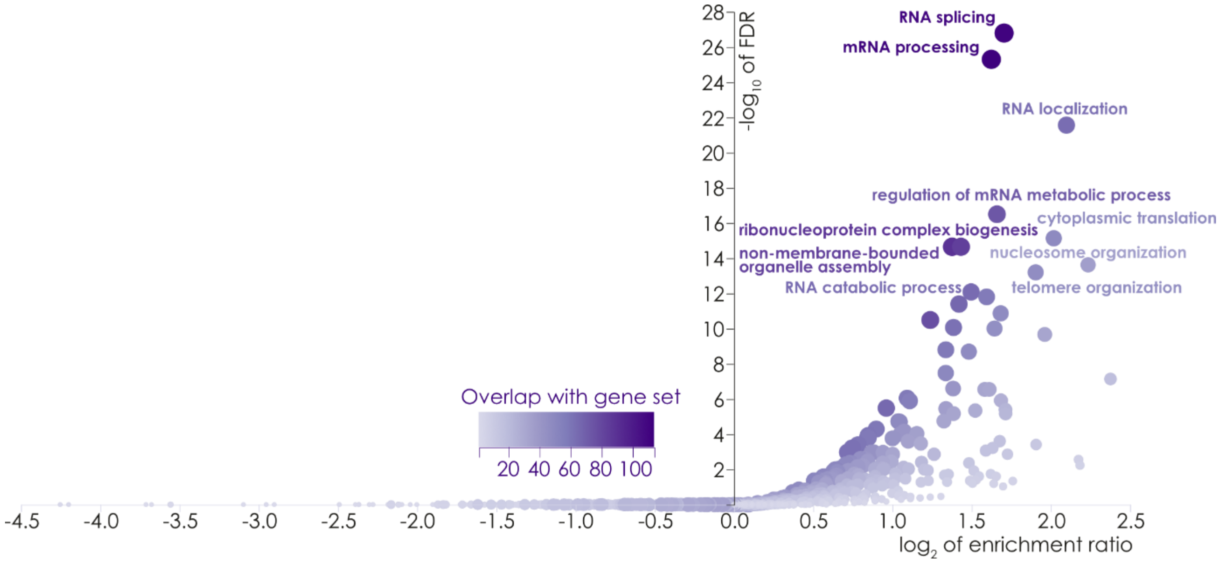

**Figure S17. Gene set enrichment analysis of literature-reported nuclear glycoproteins.** For overlooked glycopeptides from nuclear proteins in regular *O*-glycoproteomics studies in the academic literature (Table S9), we assessed their enrichment in functional groups via WebGestalt.

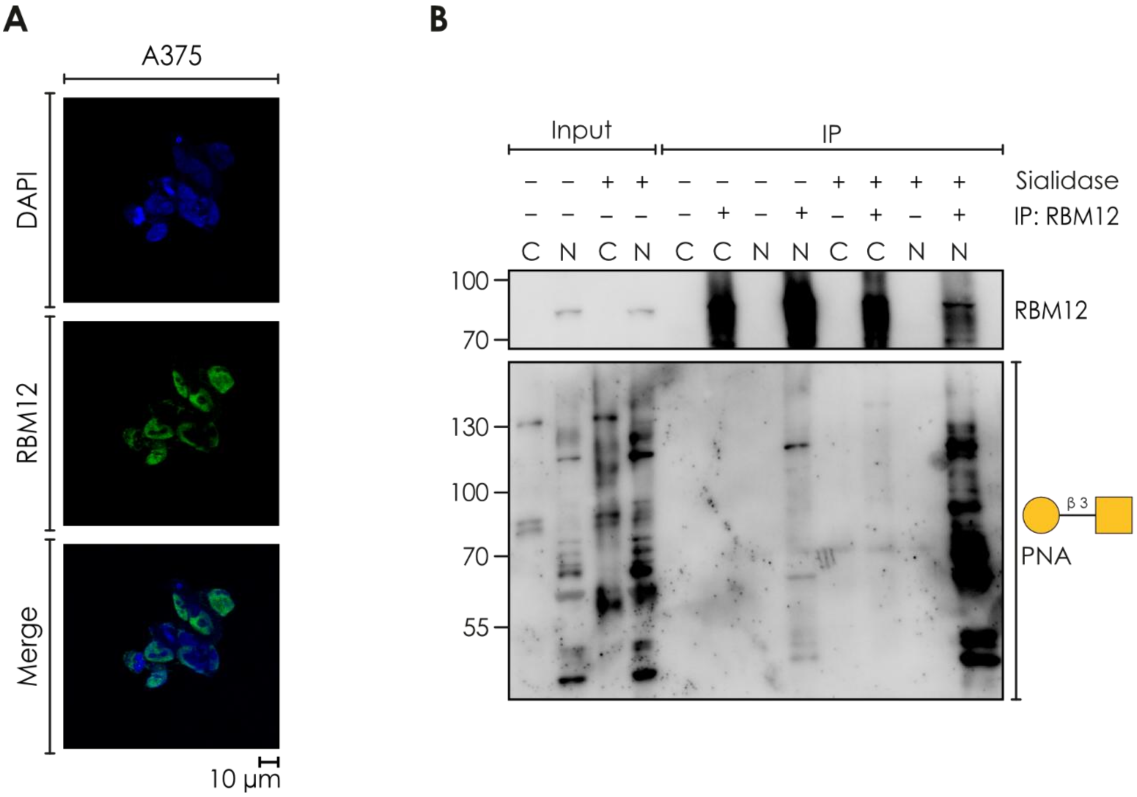

**Figure S18. RBM12/PNA co-immunoprecipitation reveals extended *O*-glycans. A)** Immunofluorescence images show that RBM12 primarily displays nuclear localization in A375 cells. Scale bar = 10 µm. **B)** Western blots against RBM12 and PNA for nuclear RBM12/PNA co-IP from A375 cells before and after sialidase treatment. We note that post-IP, the PNA-stained bands only occur with the nuclear pool of RBM12, not with its cytosolic fraction, potentially showcasing regulation of subcellular localization. Representative Western blot from three independent replications. C: cytoplasm, N: nucleus

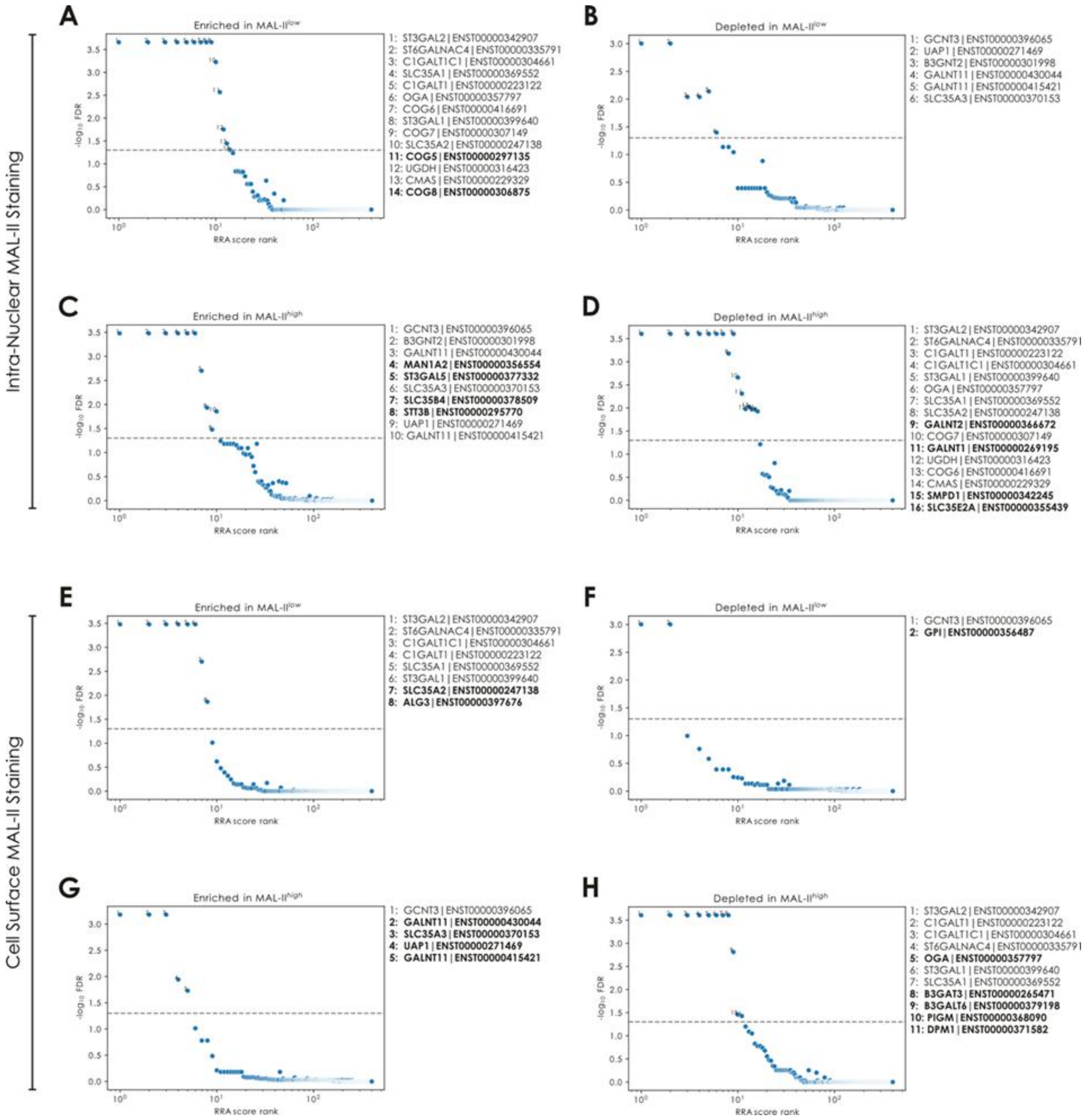

**Figure S19. Robust ranking aggregation (RRA) results of depleted gRNAs in the glycosylation-focused CRISPR/Cas9 KO screen of A549 nuclei and whole cells. A-B)** The genes for which gRNAs were significantly enriched (A) or depleted (B) in the MAL-II$^{low}$ population of A549 nuclei (n = 2). **C-D)** The genes for which gRNAs were significantly enriched (A) or depleted (B) in the MAL-II$^{high}$ population of A549 nuclei (n = 2; expected to be the inverse of gene enrichment/depletion in A-B). **E-F)** The genes for which gRNAs were significantly enriched (E) or depleted (F) in the MAL-II$^{low}$ population of A549 cells (n = 2). **G-H)** The genes for which gRNAs were significantly enriched (G) or depleted (H) in the MAL-II$^{high}$ population of A549 cells (n = 3). Genes in bold were only detected as significantly enriched/depleted in either the high- or low-staining population.

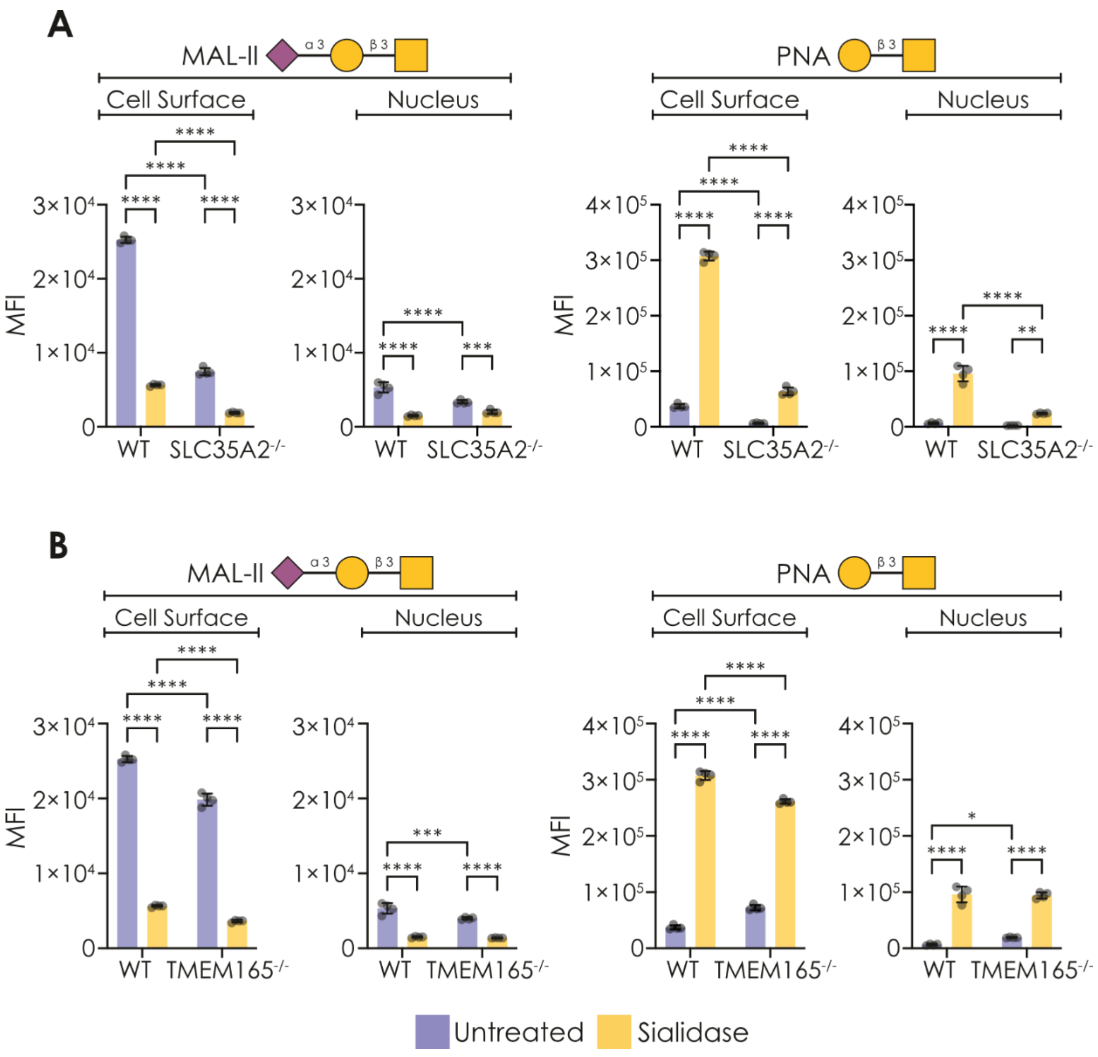

**Figure S20. Nuclear glycans are synthesized in the Golgi apparatus. A-B)** HEK293T cells with a knocked-out galactose transporter (HEK293T$^{SLC35A2-/-}$, n = 4; A) or $Mn^{2+}$ transporter (HEK293T$^{TMEM165-/-}$, n = 4; B) show decreased binding of the sialic acid- and galactose-binding lectins MAL-II and PNA to their cell surface and nuclei, supporting secretory pathway biogenesis of nuclear glycans. Significant differences were established via a one-way ANOVA with Tukey's multiple comparison test. *p < 0.05, **p < 0.01, ***p < 0.001, ****p < 0.0001.

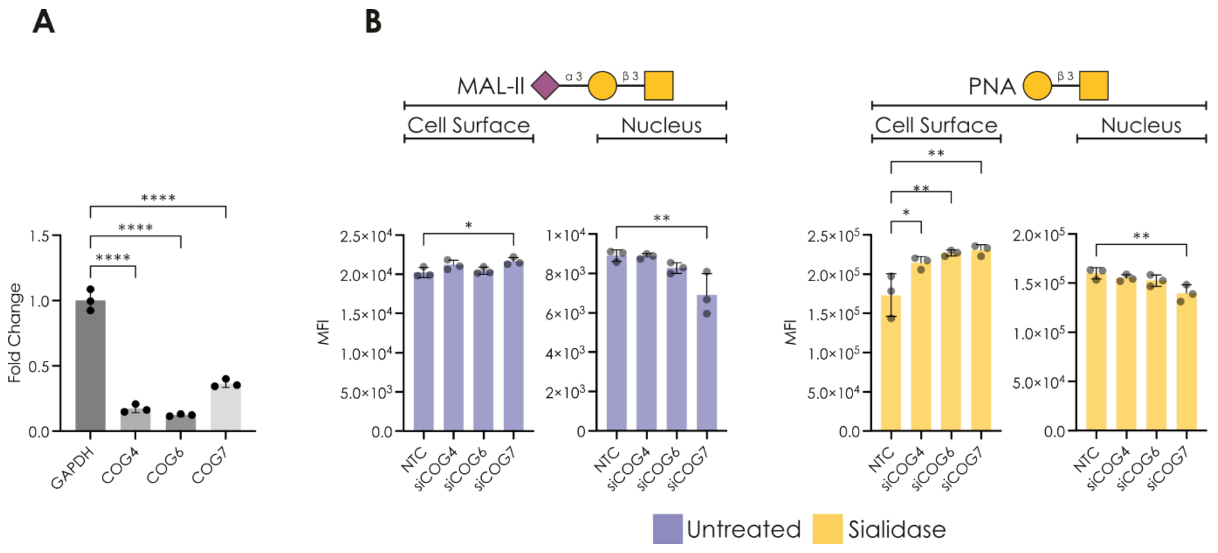

**Figure S21. siRNA-mediated depletion of COG7 selectively impairs nuclear glycosylation. A)** Real-time qPCR quantification of gene expression upon transfection of COG4, 6, or 7-targeting esiRNA in comparison to a non-targeting control (n = 3). **B)** Depletion of COG7 reduced nuclear glycosylation, yet did not impair surface glycosylation in HEK293T cells (n = 3). Significant differences were established via a one-way ANOVA with Tukey's multiple comparison test. NTC: non-targeting control. *p < 0.05, **p < 0.01, ***p < 0.001, ****p < 0.0001.

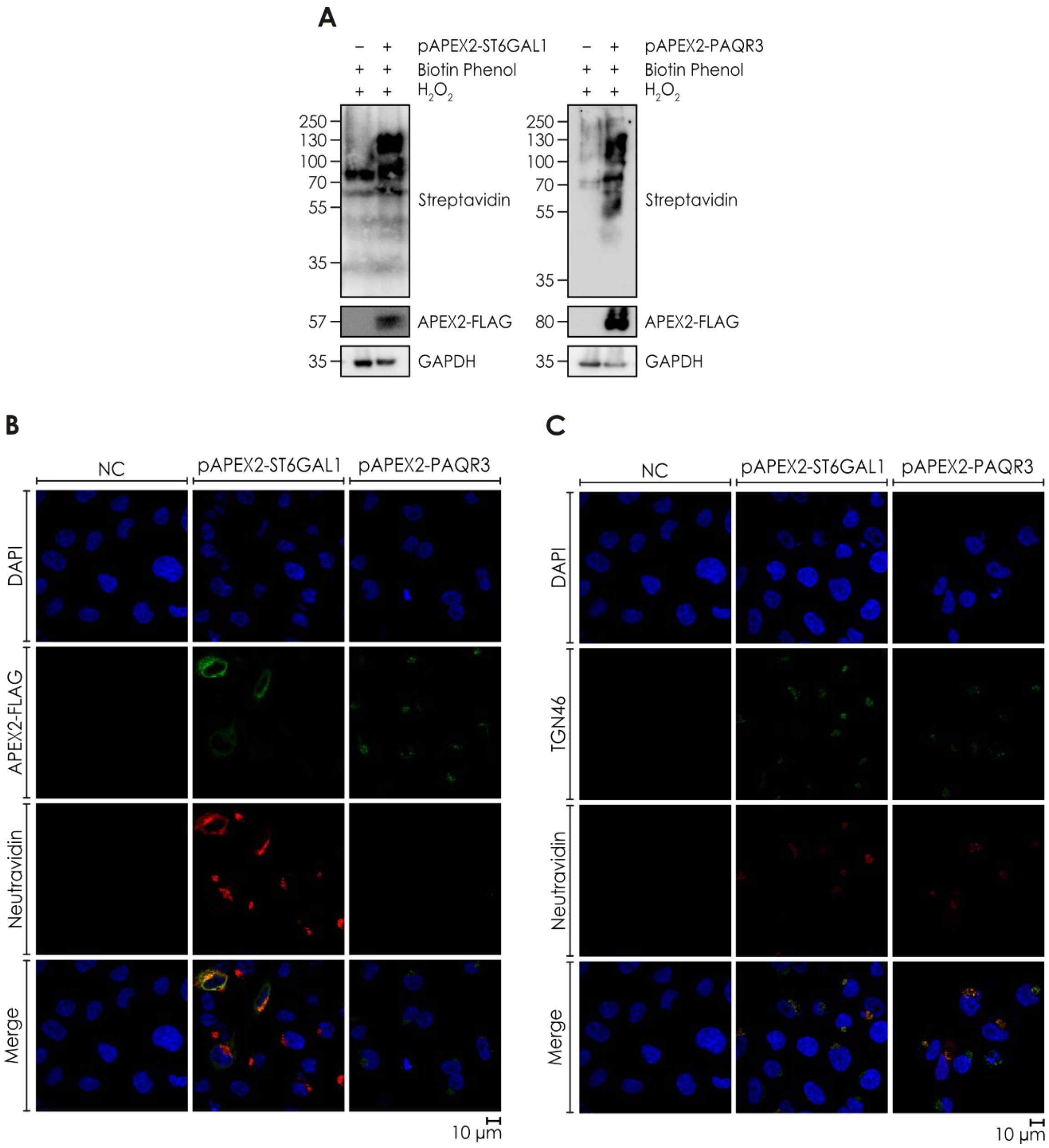

**Figure S22. The APEX2 fusion constructs localize to the Golgi for biotinylation of resident proteins. A)** Western blots showing the detection of biotinylated proteins in A375 cells upon expression of the APEX2 fusion constructs and the addition of hydrogen peroxide and biotin phenol. **B)** Fluorescence microscopy showing detection of biotinylated proteins (neutravidin) in the vicinity of the APEX2 fusion constructs (APEX2-FLAG). **C)** Fluorescence microscopy showing colocalization of biotinylated proteins (neutravidin) with the trans-Golgi marker TGN46. Scale bar = 10 µm.

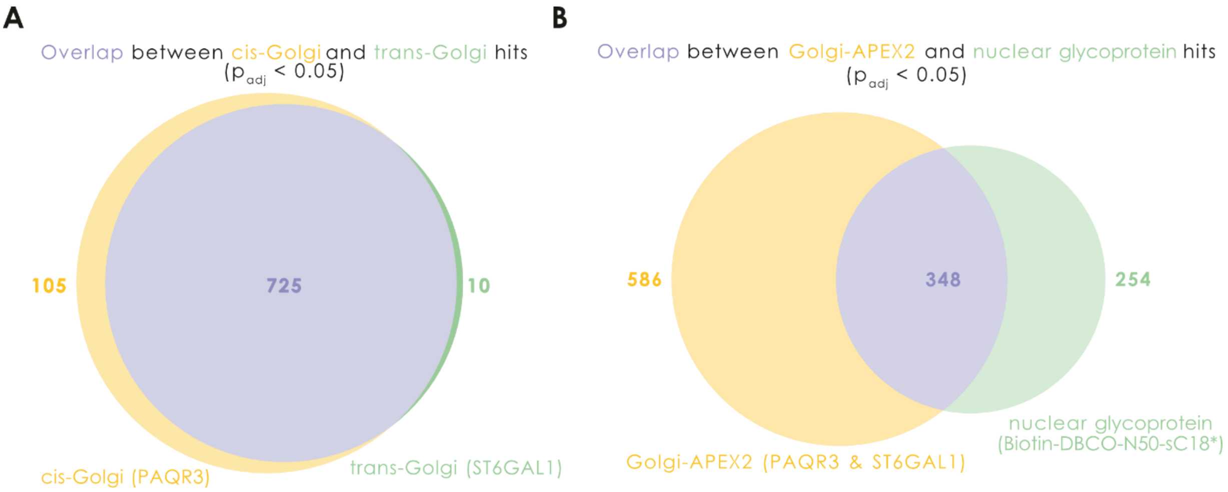

**Figure S23. Tracking nuclear glycoproteins through the secretory pathway and into the nucleus. A-B)** Only considering significant hits ($p_{adj} < 0.05$), we analyzed which nuclear proteins were enriched in both cis-Golgi (PAQR3-APEX2) and trans-Golgi (ST6GAL1-APEX2) probes (A) or in both the Golgi-APEX2 probes and the click chemistry-mediated enrichment of nuclear glycoproteins from Fig. 2E/Fig. S8 (B). Results are shown as Venn diagrams.

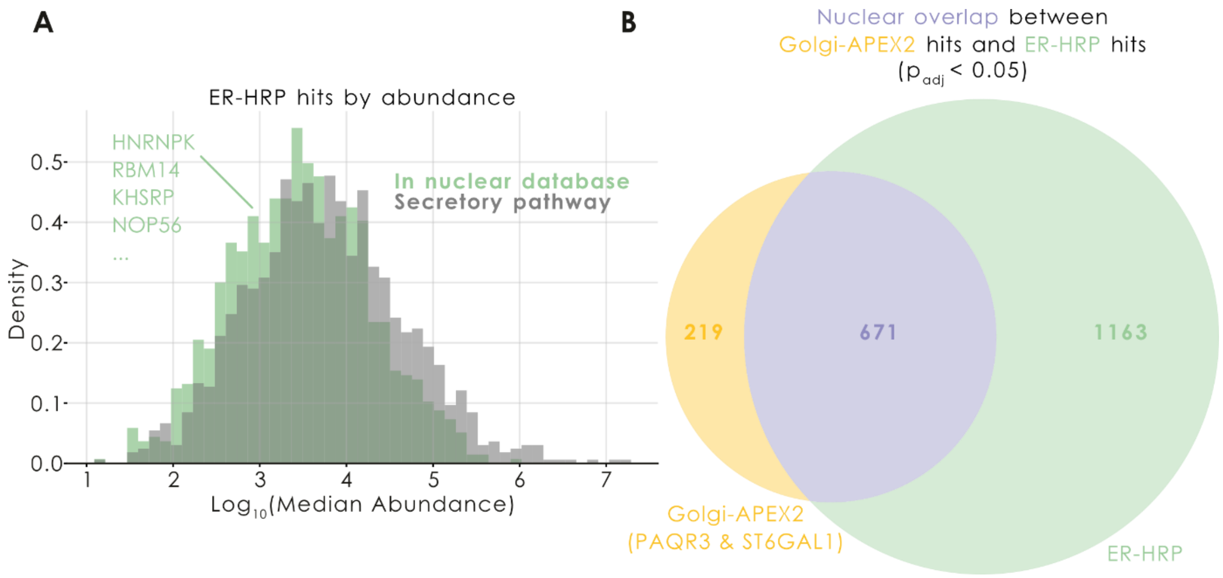

**Figure S24. Nuclear proteins transit the endoplasmic reticulum. A)** Using a dataset (Lyu et al., *ACS Chem Biol*, 2022) where lumenal ER proteins had been labeled by an ER-HRP construct (similar to APEX2, yet only active in the secretory pathway), we show a histogram of the median quantified abundance of these 3,884 ER proteins. Using our nuclear proteome database (Table S18), we then annotated 1,834 of these proteins (47.2%) as tentatively nuclear and distinguished their abundances from the classic secretory pathway hits. Example hits from this nuclear annotation are shown in the figure. **B)** Comparing the 1,834 nuclear hits with the significant hits ($p_{adj} < 0.05$) from our cis-/trans-Golgi APEX2 experiment (Fig. S23), shown via a Venn diagram, demonstrates substantial overlap between ER- and Golgi-transiting nuclear proteins.

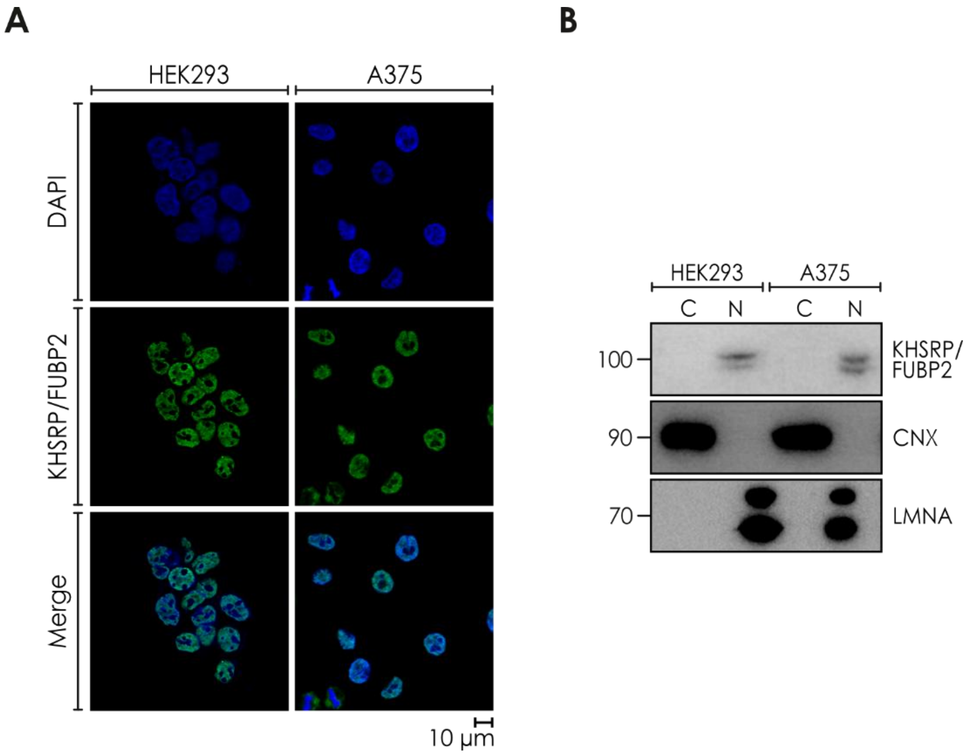

**Figure S25. KHSPR/FUBP2 localizes to the nucleus. A)** Fluorescence microscopy showing distinct nuclear localization of KHSRP/FUBP2 in HEK293 and A375 cells. Scale bar = 10 µm. **B)** Western blot detection of KHSRP/FUBP2 in ultra-pure nuclear lysates, with the outer nuclear membrane removed, of HEK293 and A375 cells. Markers include Calnexin/CNX (ER) and Lamin A/LMNA (nucleus). C: cytoplasm, N: nucleus

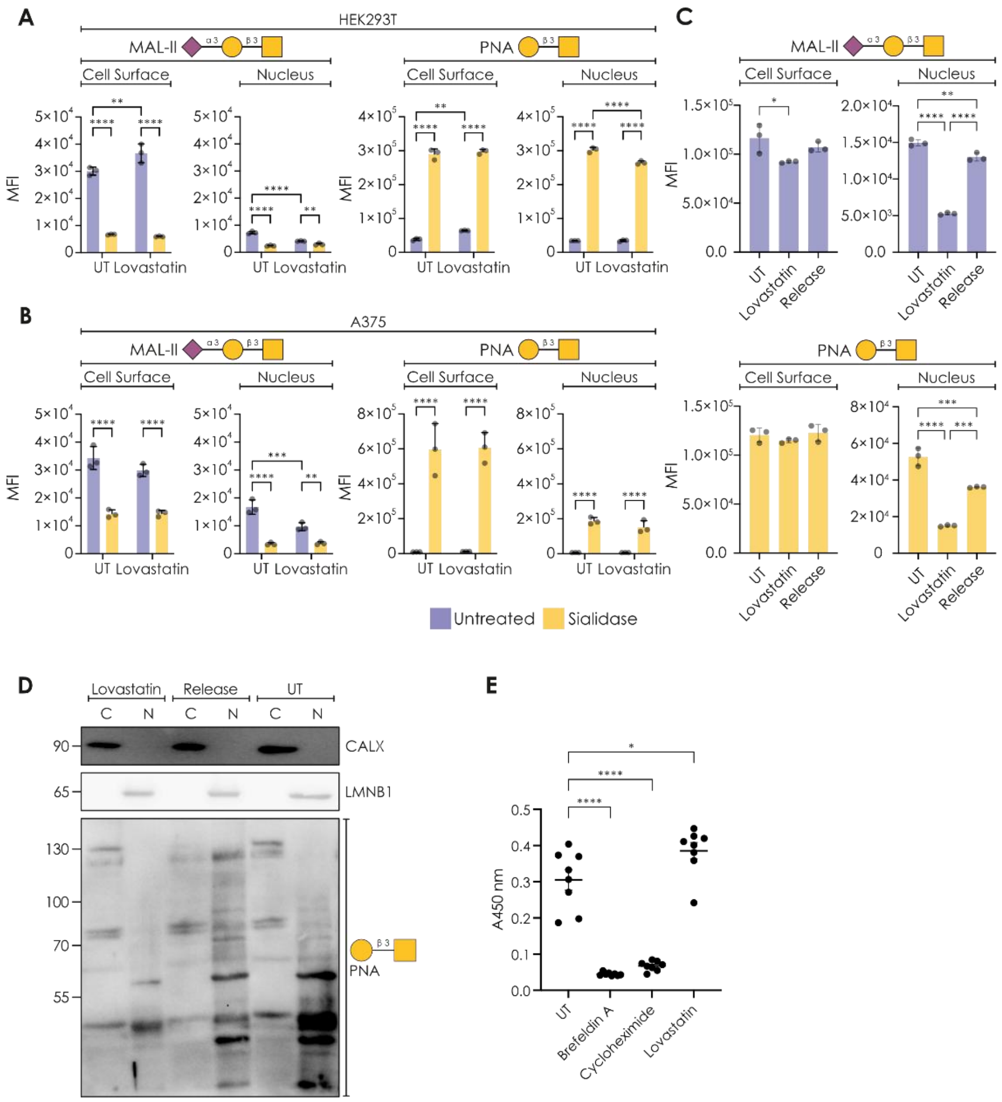

**Figure S26. Interrupting cytoskeleton dynamics selectively abrogates nuclear glycosylation. A-B)** Disrupting vesicular transport from the trans-Golgi with lovastatin reduced nuclear glycosylation, yet did not impair surface glycosylation in HEK293T cells (A; n = 3) and A375 cells (B; n = 3). **C-D)** For A375 cells, we fractionated cytoplasm (C) and nucleus (N) from DMSO-control cells (UT), cells grown under 5 µM lovastatin treatment for 24 hours (Lovastatin), or cells grown under 5 µM lovastatin treatment for 24 hours and then 1 mM mevalonate release (Release). Then, fractionations were probed via flow cytometry (C) and Western blot (D) for fractionation purity (calnexin and lamin b) and sialylation (MAL-II and PNA). Lovastatin treatment selectively decreased nuclear glycosylation levels, which could be rescued by mevalonate release treatment. **E)** Microwell plate-based detection of secreted sialoglycoproteins. A375 cells were labeled with $Ac_4ManNAz$ (50 µM, 72 h) and subsequently grown in Opti-MEM medium with or without 3 µg/mL Brefeldin A, 25 µM cycloheximide, or 3 µM lovastatin. The supernatant was collected after 24 hours and assayed in an ELISA-like setup: Azide-tagged glycoproteins in the supernatant were captured in a 96-well plate, reacted with DBCO-biotin for two hours at RT, and detected by the addition of horseradish peroxidase (HRP)-streptavidin for 30 minutes at 37°C. Stop solution (0.5 M $H_2SO_4$) was added before the absorbance was measured at 450 nm. We note that lovastatin only reduced nuclear but not secretory glycosylation. Significant differences were established via a one-way ANOVA with Tukey's multiple comparison test. Markers include Calnexin/CNX (ER) and Lamin B/LMNB1 (nucleus). C: cytoplasm, N: nucleus, UT: untreated. *$p < 0.05$, **$p < 0.01$, ***$p < 0.001$, ****$p < 0.0001$.

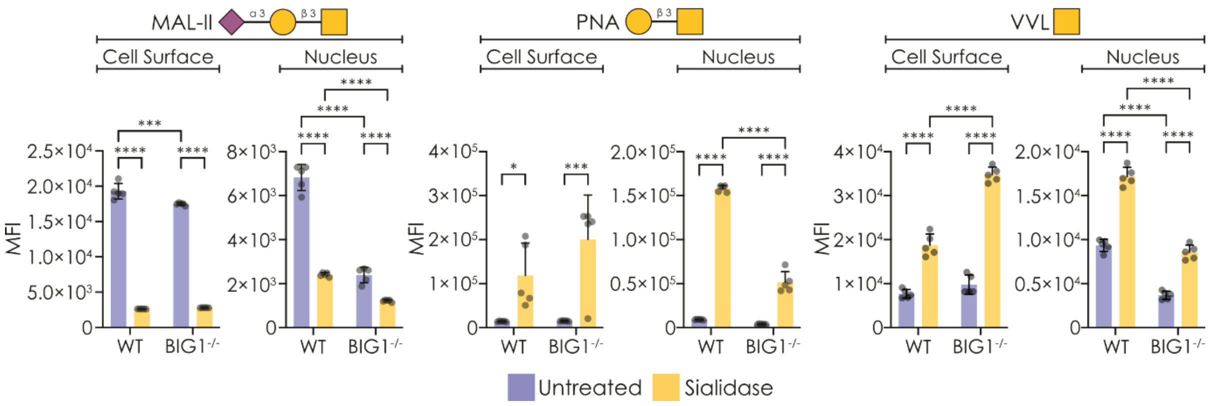

**Figure S27. Disrupting BIG1-mediated vesicular transport selectively abrogates nuclear glycosylation.** Knocking out BIG1-dependent vesicular transport selectively abrogates nuclear glycosylation. Flow cytometry of HEK293T and HEK293T$^{BIG1-/-}$ cells/nuclei, stained with MAL-II, PNA, and VVL (n = 5). Significant differences were established via a one-way ANOVA with Tukey's multiple comparison test. *p < 0.05, **p < 0.01, ***p < 0.001, ****p < 0.0001.

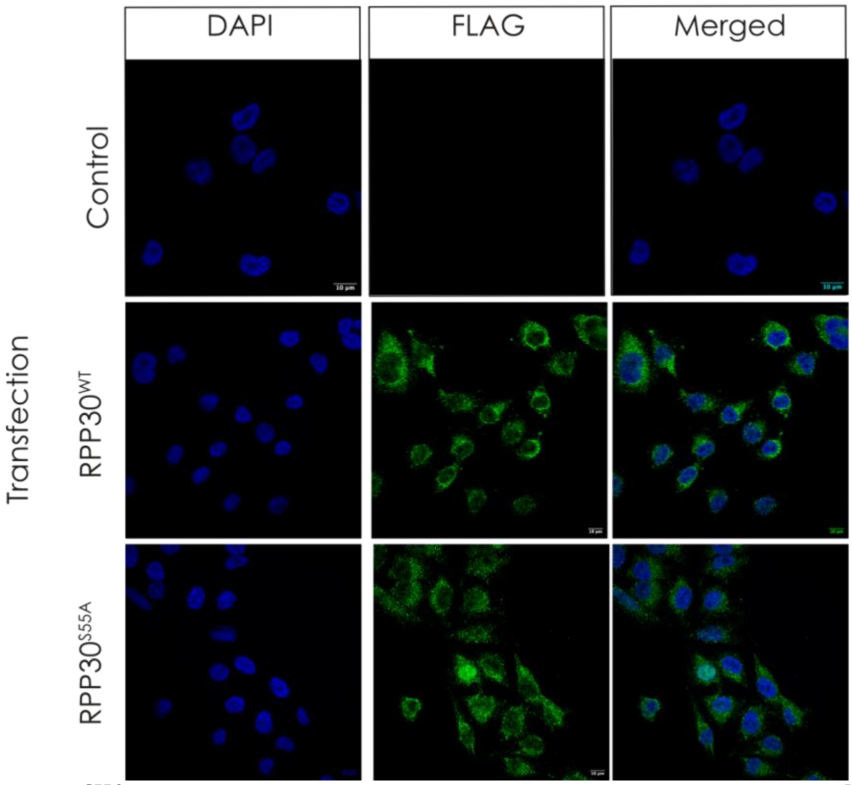

**Figure S28. RPP30$^{S55A}$ exhibits the same subcellular localization as RPP30$^{WT}$.** Shown are immunofluorescence confocal microscopy images from untransfected A375 cells or A375 expressing FLAG-RPP30$^{WT}$ or FLAG-RPP30$^{S55A}$. Nuclei were stained with DAPI and RPP30 expression with anti-FLAG. Scale bar = 10 µm.

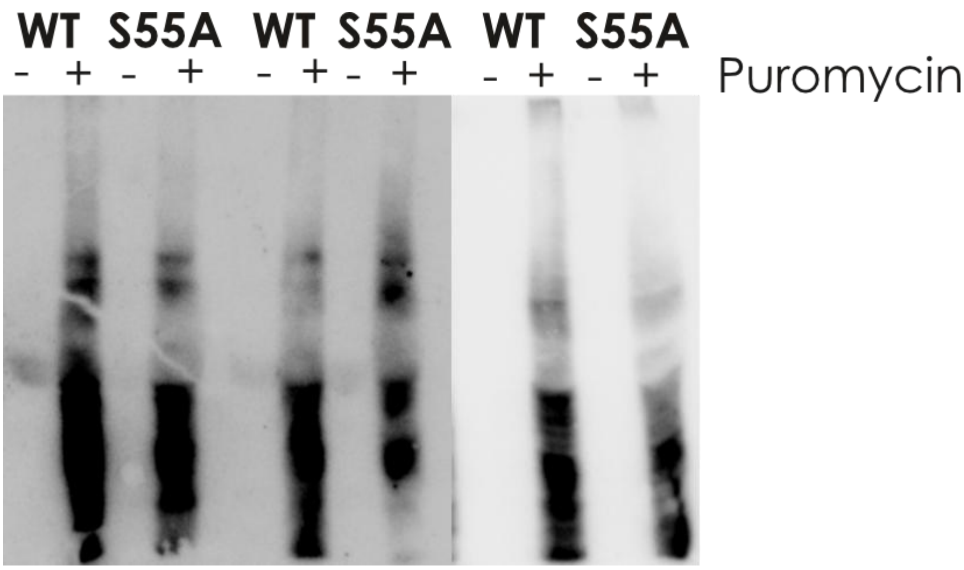

**Figure S29. Overexpressing RPP30$^{S55A}$ slows down global protein synthesis rate.** Shown is a Western blot of A375 whole-cell lysates overexpressing either RPP30$^{WT}$ (WT) or RPP30$^{S55A}$ (S55A) after treating cells for 15 minutes with water (-) or 5 µg/mL puromycin (+). After loading equal lysate volumes from matched cell counts (100,000 cells per sample), we probed samples with an anti-puromycin antibody to quantify freshly synthesized protein levels via a SUnSET assay. Per condition, we have analyzed three biological replicates (n = 3).

### Supplementary Methods

#### Proteomic sample preparation

Relative quantification was performed to compare protein expression to identify glycosylated proteins in the nucleus. The beads with attached proteins were washed twice with 1 mL 50 mM triethylammonium bicarbonate (TEAB), dissolved in 50 µL 50 mM TEAB, reduced (5 mM tris(2-carboxyethyl)phosphine (TCEP), 30 min, 37°C) and alkylated (10 mM iodoacetamide (IAA), 30 min, room temperature). Samples were digested by addition of 0.3 µg LysC/Trypsin (Promega, 37°C) for three hours. Supernatant was removed, and beads were washed with 40 µL 50 mM TEAB and combined. Peptide samples were digested overnight after extra addition of 0.3 µg LysC/Trypsin and labelled using TMTpro 18-plex isobaric mass tagging reagents (Thermo Fisher Scientific). The labelled samples were pooled into one TMT-set and purified using HiPPR Detergent Removal Resin (Thermo Scientific), according to the manufacturer´s instructions. The TMT-set was fractionated with High-pH Spin Column into 10 fractions using a gradient from 8% to 50% acetonitrile, 0.1% triethylamine in water (Pierce, Thermo Scientific). The fractions were evaporated in speed vac system and reconstituted in 15 µL 3% acetonitrile, 0.1% trifluoroacetic for LC-MS3 analysis.

#### LC-MS3 analysis

The fractions (5 µL) were analysed on an Orbitrap Lumos (Golgi-APEX2; Table S14) or Eclipse (Click peptide, Peptide pulldown, PNA plus/minus sialidase; Tables S1, S6, S7) Tribrid mass spectrometer equipped with a FAIMS Pro ion mobility system and interfaced with an Easy-nLC1200 liquid chromatography system (all Thermo Fisher Scientific). Peptides were trapped on an Acclaim Pepmap 100 C18 trap column (100 µm x 2 cm, particle size 5 µm, Thermo Fisher Scientific) and separated on an in-house packed analytical column (35 cm x 75 µm, particle size 3 µm, Reprosil-Pur C18, Dr. Maisch) using a stepped gradient from 5% to 35% acetonitrile in 0.2% formic acid over 77 min at a flow of 300 nL/min. FAIMS Pro was alternating between the compensation voltages (CV) of -40, -60, and -80, and the same data-dependent settings were used at all CVs. The precursor ion mass spectra were acquired at a resolution of 120 000 and an *m/z* range of 375-1375. Using a cycle time of 1 second the most abundant precursors with charges 2–7 were isolated with an *m/z* window of 0.7 and fragmented by collision induced dissociation (CID) at 35%. Fragment spectra were recorded in the ion trap at Rapid scan rate. Dynamic exclusion was set to 60 sec. The ten most abundant MS2 fragment ions were isolated using multi-notch isolation for further MS3 fragmentation. MS3 fragmentation was performed using higher-energy collision dissociation (HCD) at 55% and the MS3 spectra were recorded in the Orbitrap at 50,000 resolution and an *m/z* range of 100–500.

#### Proteomic data analysis

Raw files were processed and analyzed with Proteome Discoverer (Ver 3.0, Thermo Scientific) The data was matched against *Homo Sapiens* SwissProt database (20,597 entries, Feb 2024) or a Nuclear database with selected proteins (7,224 entries), together with a contaminant database (250 entries, from Thermo Fisher) in some experiments (Peptide pulldown, PNA plus/minus sialidase; Tables S1, S7). Sequest was the search engine, precursor and fragment ion tolerance were set to 5 ppm and 0.6 Da, and tryptic peptides were accepted with 1 missed cleavage. Methionine oxidation was set as a variable modifications and cysteine carbamidomethylation, TMTpro on lysine and peptide N-termini were set as fixed modifications. Percolator was used for PSM validation with a strict FDR threshold of 1%. For quantification, TMT reporter ions were identified in the MS3 HCD spectra with 3 mmu mass tolerance and the TMT reporter intensity values for each sample were normalized on the total peptide amount. In study "PNA plus/minus sialidase" (Table S7), Inferys Rescoring node was used and the TMT reporter intensity values for each sample were normalized on total peptide amount. The SPS threshold was set to 65%, a Sequest HT threshold score of 2 was chosen. Only unique peptides were used for relative quantification and proteins were required to pass a protein FDR of 5%.

#### Synthesis and purification of Lys(Biotin)-PEG$_4$-DBCO-Cys-N50-sC18*

The peptide was chain assembled using Rink amide resin (substitution 0.66 mmol/g). All amino acid derivatives were acquired from Carbolution GmbH (St. Ingbert, Germany) and used without further purification, except for the Fmoc-Lys(Biotin)-OH which was acquired from Iris Biotech GmbH (FAA1443, Marktredwitz, Germany). The Sulfo DBCO-maleimide was acquired from Vector Laboratories inc. (CCT1230, Newark, CA, USA). The core Cys-N50-sC18* peptide sequence (CVQRKRQKLMPGLRKRLRKFRNK) was synthesized using automated SPPS on a Intavis MultiPep CF (CEM Corporation, Matthews, NC, USA). Amino acid couplings were carried out with the molar ratio of (4):(4):(8) of (Fmoc-protected amino acid):(HBTU):(NMM) at room temperature for 60 minutes and deprotection was achieved in 20% (v/v) piperidine in DMF for 15 minutes at room temperature. Fmoc-PEG$_4$-OH and Fmoc-Lys(Biotin)-OH were coupled manually with the molar ratio of (4):(4):(8) of (Fmoc-protected amino acid):(HATU):(DIPEA) at room temperature for 60 min, to ensure complete reactions, all couplings were carried out twice. Manual deprotection was achieved in 20% (v/v) piperidine in DMF for 30 minutes. The peptide proceeded to standard cleavage from resin using a mixture of TFA (92.5%), $H_2O$ (2.5%), TIPS (2.5%) and EDT (2.5%) for 3 hours at room temperature. TFA was removed using $N_2$ and the resultant residue was suspended in ice-cold diethyl ether. The mixture was then centrifuged (5 min, 4600 RPM) after which the supernatant was decanted into the waste. The remaining solid was washed twice by ice-cold diethyl ether and subjected to purification. For the purification and characterization of peptides, two eluent systems were used. Mobile phase A was 0.5% acetic acid in MQ-$H_2O$, mobile phase B was 0.1% acetic acid in ACN and detection was done at 214 nm. The crude peptide was dissolved in a mixture of ACN in $H_2O$ and purified by semi-preparative HPLC using a Waters 600 system (Waters, Milford, MA, USA) equipped with a C18 column (MultoKrom 100 – 5 C18, 5 µm particle size, 100 Å pore size, 250 x 20 mm, CS Chromatographie Service, Langerwehe, Germany) and a gradient of mobile phase A and mobile phase B from 0% B to 40% B over 50 min at 8 mL/min. Analytical RP-HPLC was carried out on a Waters XC e2695 system employing a Waters PDA 2998 diode array detector equipped with a ISAspher 100-3 C18 (C18, 3.0 µm particle size, 100 Å pore size, 50×4.6 mm, Isera GmbH, Düren, Germany) at a flow rate of 2 mL/min using a gradient of mobile phase A and mobile phase B from 10% B to 90% B over 10 min at 2 mL/min. The molecular weight of the purified peptides was confirmed by ESI mass spectrometry on a Waters Synapt G2-Si ESI mass spectrometer equipped with a Waters Acquity UPLC system using a Xela C18 column (C18, 1.7 µm particle size, 80 Å pore size, 50×3.0 mm, Isera GmbH). The sulfo DBCO-maleimide was attached by preparing a solution of peptide (2 mM in PBS, pH 7.4) and mixing with a solution of Sulfo DBCO-maleimide (2 mM in PBS, pH 7.4) reacting in a Thermomixer (Eppendorf SE, Hamburg, Germany) for 2 hours at room temperature and used without further purification. The coupling was confirmed using ESI-MS and analytical HPLC.

| **Peptide** | **Calculated Mw [g/mol]** | **Experimental Mw [g/mol]** | **tR [min]** |
|---|---|---|---|
| Lys(Biotin)-PEG$_4$-Cys-N50-sC18* | 3540.10 | 3540.04 | 2.54 |
| Lys(Biotin)-PEG$_4$-DBCO-Cys-N50-sC18* | 4277.56 | 4277.52 | 3.31 |

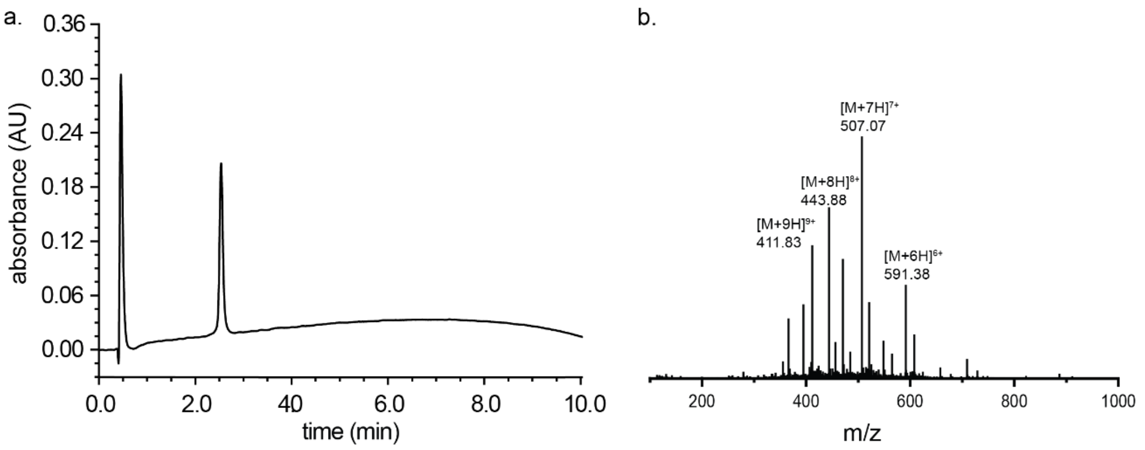

**A) HPLC chromatogram and B) mass spectrum for Lys(Biotin)-PEG4-Cys-N50-sC18*.**

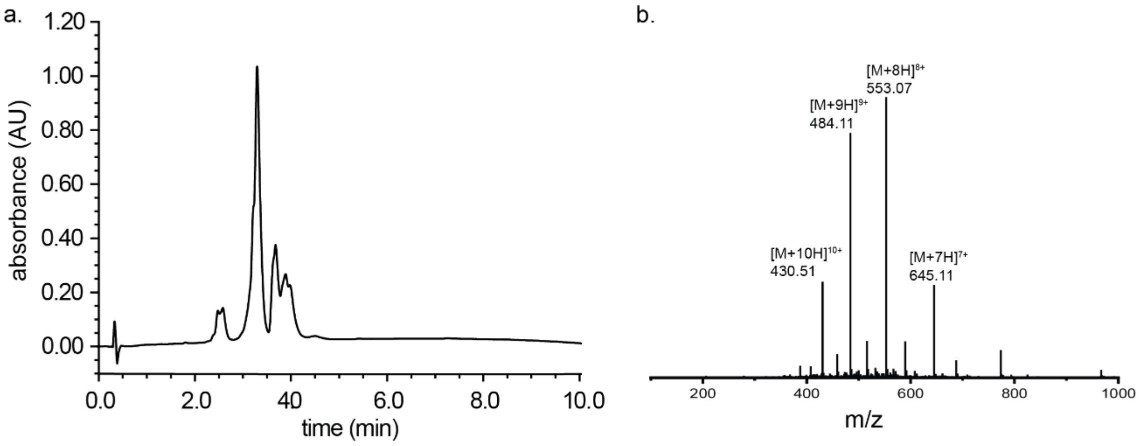

**A) HPLC chromatogram and B) mass spectrum for Lys(Biotin)-PEG4-DBCO-Cys-N50-sC18*.**

#### Design of the Lys(Biotin)-PEG4-DBCO-Cys-N50-sC18*

We aimed to design a peptide that could specifically label the incorporated azido-mannose moieties within the cell nucleus. To achieve this, the peptide contained three distinct features. Firstly, a core cell-penetrating peptide (CPP) sequence was chosen that combined the features of the nuclear localization signal and a shortened version of the CPP sC18*, which was reported to efficiently reach the nucleus in HeLa and MCF-7 cell lines.[1] Secondly, a DBCO moiety was envisioned to allow for a biorthogonal, copper free click reaction with the cell's nucleus.[2, 3] Lastly, a biotin functionality was incorporated to allow for facile detection of the peptide in assays such as flow cytometry, Western blotting and streptavidin pull-downs. In our initial attempt, the N50-sC18* sequence was chain assembled using automated solid-phase synthesis, after which a biotinylated lysine residue, a polyethylene glycol (PEG) spacer spanning four units, and a carboxylic acid derivative of DBCO were incorporated manually. However, this peptide appeared to not be functional—while it could reach the nucleus of the cell, it was not able to perform the click reaction with any azides. It has been described in literature that DBCO can spontaneously rearrange under acidic conditions, with low acid concentrations typically used in HPLC purification of peptides already enough to induce rearrangement to a non-reactive tetracyclic product.[4] Therefore, it is highly likely that during treatment with the TFA cocktail used to remove the peptide from the resin, the DBCO moiety had rearranged completely. Recently, it was reported that by addition of a Cu(I) salt to the TFA cleavage mixture, this acid-mediated rearrangement can be prevented.[5] Unfortunately, addition of this Cu(I) salt did not seem to affect the reactivity of the final DBCO-labeled peptide in our cellular assays and the peptide as a whole appeared inactive. To circumvent the issues associated with acid-mediated rearrangement of DBCO, we decided to incorporate the DBCO group after the peptide had been purified under acidic conditions. To this end, we incorporated a cysteine residue, amenable for simple maleimide conjugation using a Sulfo DBCO reagent. The sulfo variant of DBCO was chosen as it is reported to be beneficial for water solubility. To our delight, the purified, cysteine-containing peptide was functionalized by using an equimolar solution of the DBCO-maleimide within two hours. Now, the peptide appeared functional and could efficiently label the azides.

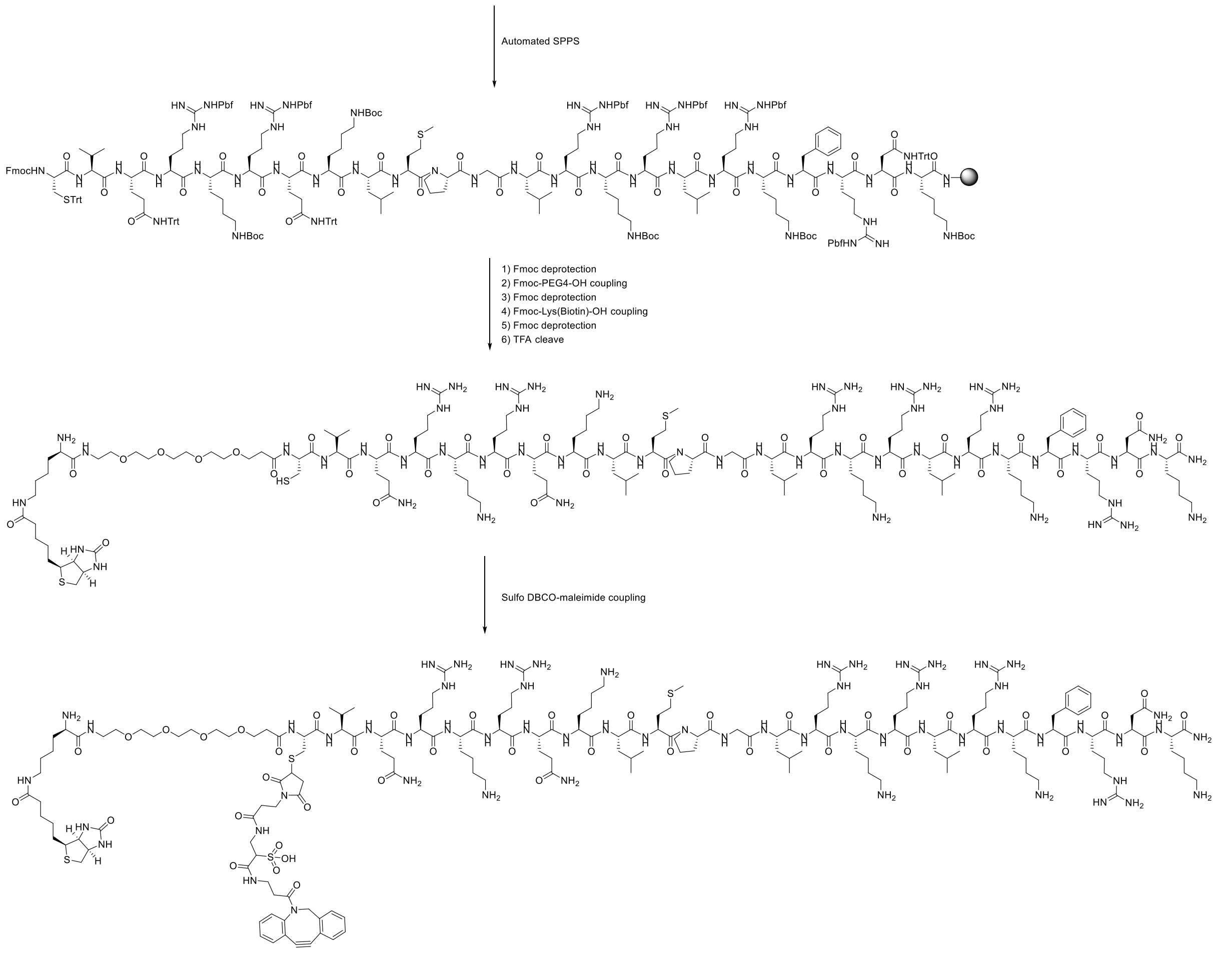

Synthetic scheme for Lys(Biotin)-PEG4-DBCO-Cys-N50-sC18*.

#### References

1. A. Gronewold, M. Horn and I. Neundorf, *Beilstein J. Org. Chem.*, 2018, **14**, 1378-1388.
2. H. Y. Yoon, D. Lee, D. K. Lim, H. Koo and K. Kim, *Adv. Mat.*, 2022, **34**, 2107192.
3. C. G. Gordon, J. L. Mackey, J. C. Jewett, E. M. Sletten, K. N. Houk and C. R. Bertozzi, *J. Am. Chem. Soc.*, 2012, **134**, 9199-9208.
4. M. Chigrinova, C. S. McKay, L.-P. B. Beaulieu, K. A. Udachin, A. M. Beauchemin and J. P. Pezacki, *Org. Biomol. Chem.*, 2013, **11**, 3436-3441.
5. P. W. Erickson, J. M. Fulcher, P. Spaltenstein and M. S. Kay, *Bioconjugate Chem.*, 2021, **32**, 2233-2244.
